# Kinetic Diagram Analysis: A Python Library for Calculating Steady-State Observables of Biochemical Systems Analytically

**DOI:** 10.1101/2024.05.27.596119

**Authors:** Nikolaus Carl Awtrey, Oliver Beckstein

**Affiliations:** Department of Physics, Arizona State University, Tempe AZ, USA; Center for Biological Physics, Arizona State University, Tempe AZ, USA

## Abstract

Kinetic diagrams are commonly used to represent biochemical systems in order to study phenomena such as free energy transduction and ion selectivity. While numerical methods are commonly used to analyze such kinetic networks, the diagram method by King, Altman and Hill makes it possible to construct exact algebraic expressions for steady-state observables in terms of the rate constants of the kinetic diagram. However, manually obtaining these expressions becomes infeasible for models of even modest complexity as the number of the required intermediate diagrams grows with the factorial of the number of states in the diagram. We developed *Kinetic Diagram Analysis* (`KDA`), a Python library that programmatically generates the relevant diagrams and expressions from a user-defined kinetic diagram. `KDA` outputs symbolic expressions for state probabilities and cycle fluxes at steady-state that can be symbolically manipulated and evaluated to quantify macroscopic system observables. We demonstrate the `KDA` approach for examples drawn from the biophysics of active secondary transmembrane transporters. For a generic 6-state antiporter model, we show how the introduction of a single leakage transition reduces transport efficiency by quantifying substrate turnover. We apply `KDA` to a real-world example, the 8-state free exchange model of the small multidrug resistance transporter EmrE of Hussey et al (*J General Physiology* **152** (2020), e201912437), where a change in transporter phenotype is achieved by biasing two different subsets of kinetic rates: alternating access and substrate unbinding rates. `KDA` is made available as open source software under the GNU General Public License version 3.

## 1 Introduction

Cellular processes at different scales are commonly represented as kinetic diagrams or kinetic graphs where distinct states form the nodes of the graph and the edges describe the reactions that interconvert between the states.^1–6^ Diagrams can be represented at different levels of detail, from individual biochemical reactions with associated rates to cycles and a generic undirected graph (Fig. 1). The discrete-state formalism requires systems with a separation of time scales such that equilibrium is achieved much faster within a state than between states. ^7^ System dynamics are approximated as discrete transitions between individual states. Each transition is quantified by its forward and reverse rate, resulting in a set of master equations that relates the change in the population of each state to fluxes between states via mass-action kinetics.^8,9^ Such kinetic models are general and suitable to quantitatively describe processes out of equilibrium that are driven by external sources of free energy. The underlying equations are commonly solved numerically.^10,11^ However, it is also possible to obtain exact algebraic solutions under steady state conditions (with equilibrium included as a special steady state) with the diagram method developed by King and Altman ^12^ and Hill. ^13,14^ The diagram method is applicable to cyclic diagrams and ultimately yields rational algebraic expressions for state probabilities and cycle fluxes in terms of products of rate constants. It consists of rules to decompose cyclic graphs into a large number of auxiliary graphs that correspond to individual terms in the algebraic expressions. Although the diagram method is restricted to steady state and cannot comment on the time evolution of the system, it offers the ability to directly identify and calculate functionally relevant fluxes and provides the functional form of all observables, amenable to symbolic manipulation.

**Figure 1:**
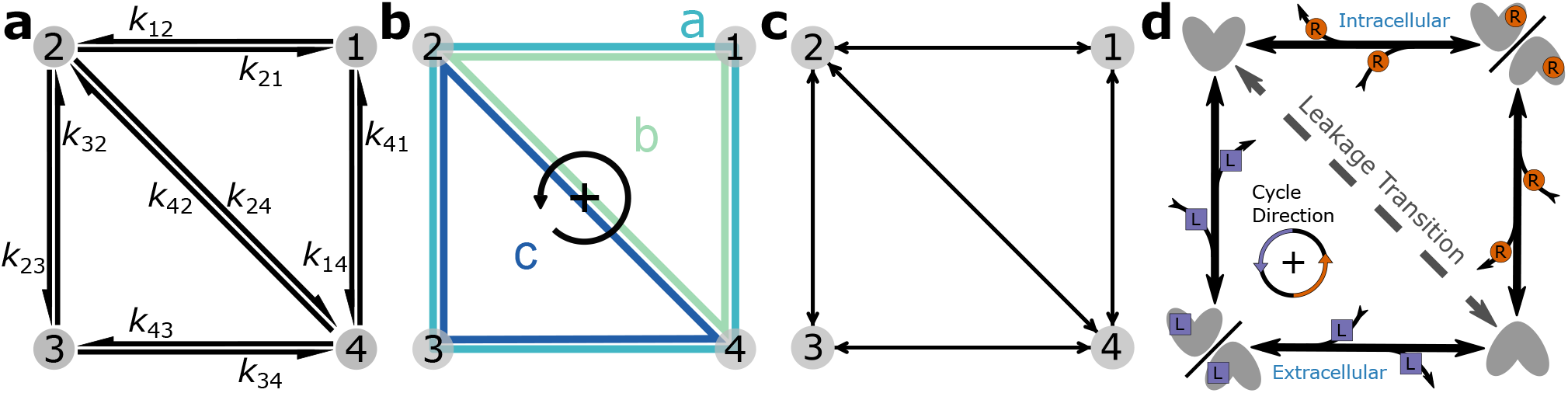
Kinetic Diagram Representations. (a) 4-state kinetic diagram with each one-way transition labeled with the corresponding reaction rate, *k*_*ij*_. (b) The same diagram highlighting the 3 distinct cycles, labeled *a, b*, and *c*, with the positive cycle direction defined as counter-clockwise. (c) `KDA` rendering of the same kinetic diagram (via `NetworkX`) with transition pairs (i.e., *i* → *j, j* → *i*) represented by double-arrows. (d) Simple 4-state antiporter model. Cycle *a* transports substrate L and driving ion R across the cell membrane in opposite directions. Leakage cycles *b* and *c* transport ligands down their electrochemical gradients independently. Fast-equilibrium between conformations is assumed for states 1 and 3.

The diagram method quantifies cyclic processes. They are represented by closed loops in the kinetic diagram called *kinetic cycles*. Cycles generally contain three or more states but can also be found in 2-state models with multiple transitions between the pair of states. Cycles spanning all states in a kinetic diagram are known as *Hamiltonian cycles*. They often describe the primary physiological function of the system; we call such cycles *productive cycles*. Simple diagrams generally have one cycle while complex diagrams may have several cycles or even several Hamiltonian cycles. Cycles are called *leakage cycles* when they offer an alternative pathway to the productive cycle since these cycles can reduce the efficiency of the primary function of the productive cycle. For example, the 4-state model in Fig. 1b contains three cycles: the Hamiltonian cycle *a* and two leakage cycles, *b* and *c*. If the primary function of this state model is achieved with completions of cycle *a*, cycles *b* and *c* will reduce the number of desired cycle completions by providing different pathways, ultimately reducing efficiency. Thus, quantifying these cycle completions can be important for understanding cyclic phenomena in these systems.

In practice, the diagram approach is difficult to carry out except for the simplest diagrams because of a combinatorial explosion of the intermediate graphs that must be constructed. For a diagram with *N* states, the number of intermediate subgraphs (and thus terms in the algebraic expressions) scale as roughly *N*!,^15^ where each subgraph represents a kinetic rate-product of *N* − 1 rates. However, generating graphs with the associated book keeping are easily handled by computers, suggesting a programmatic solution. Thus, to make the diagram method more accessible, we developed `Kinetic` `Diagram` `Analysis` (`KDA`), a Python package for programmatically generating the required subgraphs and symbolic expressions from a user-defined kinetic diagram. `KDA` leverages modern Python packages for graphs (`NetworkX`^16^) and symbolic expressions (`SymPy`^17^) to produce exact algebraic steady-state solutions for state probabilities and fluxes for diagrams of, in principle, arbitrary complexity. Our Python implementation is designed to be modular so that it can be easily integrated into existing workflows or used interactively in Jupyter notebooks; overall, we provide an accessible and interoperable solution to generate symbolic solutions of the diagram method.

Other software has been developed to generate the algebraic expressions for enzymes and transporters. Previous work performed by Pring^18^ and Rhoads,^19^ Lam and Priest, ^20^ Cornish-Bowden, ^21^ Kinderlerer and Ainsworth, ^22^ Straathof and Heijnen, ^23^ Fromm and Fromm,^24^ Varon et al.,^25^ Yago et al.,^26^ Qi et al.,^27^ and Loriaux et al. ^28^ have all focused on computer-based algorithms and programs for creating algebraic rate expressions. Additionally, recent work by Nam et al. ^29^ provides a general framework for determining steady states in terms of reaction rates using the Matrix-Tree theorems of graph theory. Currently the only programs available for download for computer-aided derivation of kinetic rate equations are `KAPattern`^27^ and `py-substitution`.^28^ `KAPattern` is a standalone application which uses the schematic method of King and Altman^12^ and the topological theory of graphs^30^ to generate King-Altman graphs and output kinetic rate equations. `py-substitution`, implemented in Matlab and Maple, uses linear algebra methods to derive steady state expressions for mass action models with arbitrary structure. While `KAPattern` and `py-substitution` yield similar outputs to `KDA`, our code and tests are open source and provide a high degree of interoperability via the Python ecosystem.

To demonstrate the use of `KDA` we are focusing on the biophysics of secondary active transporters,^31–33^ an area covered originally in Hill’s work^13,14^ with many recent applications of kinetic models.^1–6,34–37^ These membrane proteins use the free energy stored in a transmembrane ionic gradient to drive energetically uphill transport of a substrate (another ion or small molecule) by alternating between multiple protein conformations in a manner that is coupled to ion and substrate binding. Secondary active transporters are prime examples of free energy transduction in biology^14^ and serve as systems to study non-equilibrium behavior at the molecular scale. The transport process involves the protein moving through distinct states characterized by specific bound and unbound conformations. These conformations are categorized as either inward-facing (providing intracellular access) or outward-facing (allowing extracellular access). For instance, in an inward-facing conformation, a ligand in the intracellular solution can bind to the protein inducing a transition from an unbound inward-facing state to a bound inward-facing state. These binding events and accompanying conformational changes occur in a cyclic manner, enabling the transport of one or more ligands across the cell membrane per cycle completion. Cycles are broadly classified as either *symport* or *antiport* cycles: Symport cycles involve the simultaneous movement of both driving ion and substrate in the same direction (e.g. from intracellular to extracellular) while antiport cycles involve transport of the ion and substrate in opposite directions.

For instructional purposes we include a simple 4-state kinetic diagram representing an antiporter model (Fig. 1d). This model features two ligands, driving ion R and substrate L, where the gradient of the driving ion is stronger than the gradient of the substrate, as expressed with the concentrations [R_ext_]/[R_int_] ≫ [L_ext_]/[L_int_], and both gradients are directed from the external side to the internal side of the cell ([R_ext_] > [R_int_], [L_ext_] > [L_int_]). The system states are defined in the following way: 1 (driving ion-bound inward/outward-facing conformations with fast-equilibrium), 2 (inward-facing unbound state), 3 (substrate-bound inward/outward-facing conformations with fast-equilibrium), and 4 (outward-facing unbound state). The physiological function of this transporter is to use the energetically downhill influx of the driving ion (binding to the transporter 4 → 1, rapid isomerization of the transporter between outward facing and inward facing conformation, and internal release 1 → 2) to move substrate L against its gradient from the inside (binding reaction 2 → 3, isomerization, and release 3 → 4) to the outside. The cycle *a* (4 ↔ 1 ↔ 2 ↔ 3 ↔ 4) is the productive Hamiltonian cycle (where double arrows indicate that each reaction in the cycle is reversible). A positive (defined as counterclockwise in this diagram) completion of cycle *a* (4 → 1 → 2 → 3 → 4) moves driving ion R from outside to inside while simultaneously transporting substrate L from inside to outside against its concentration gradient, thus exhibiting antiport function. The model also contains a leakage transition 2 ↔ 4 resulting in a total of three cycles in the diagram (*a, b, c* in Fig. 1b). Leakage cycle *b* reduces the transport efficiency of the system by uncoupling the ligands, enabling R to move down its concentration gradient without including movement of substrate L; similarly, cycle *c* allows L to move from the outside to the inside down its own gradient, in the opposite direction compared to the productive cycle *a*. This simplified model is used as an instructional example throughout the paper although a more realistic antiporter model is discussed in the Results.

The paper is organized as follows: We begin with a review of cycle kinetics and intermediate diagram generation, closely following Hill. ^15^ We then introduce the algorithms for generating the various diagrams and summarize `KDA`’s capabilities. Next we validate the `KDA` symbolic solutions against numerical solutions from a matrix solver and evaluate the performance against the numerical approach. We then apply `KDA` to two examples, namely the transport cycles of active secondary transporters. For a simple generic 6-state antiporter model (similar to recent work by Kolomeisky and colleagues^37^), we show how the introduction of a single leakage transition reduces transport efficiency by directly calculating the turnover number of the transporter (productive cycle completions per unit time) as a function of the leak rate. We then study a real-world example, the 8-state free exchange model of the small multidrug resistance transporter EmrE from the work of Henzler-Wildman *et al*.^36^ where we confirm by direct calculation of turnover numbers that a change in transporter phenotype can be achieved by biasing two different subsets of kinetic rates, namely either rates related to alternating access or specific ligand unbinding rates. We conclude with a discussion of the equivalence of operational fluxes, which can only be obtained with the diagram approach, and sums of specific net transition fluxes, which can be computed with any approach.

## 2 Theory and Methods

We begin with reviewing the basics of cycle kinetics and the principles of the diagram method, following Hill. ^15^ We will then explain how `KDA` generates diagrams and symbolic expressions, and highlight some of the capabilities used in our analysis.

### 2.1 Kinetic Rates and Reactions

Kinetic diagrams are composed of individual reactions linked by common states where pairs of states follow unimolecular kinetics.^9,13^ These reactions are described by mass-action kinetics and corresponding forward (*k*_*ij*_) and reverse rate constants (*k*_*ji*_).^15^ We consider three types of rate constants: first-order, pseudofirst-order, and second-order. A first order reaction

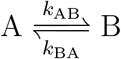

is characterized by the *first-order rate constants k*_AB_ and *k*_BA_ that are concentration-independent with units s^−1^. The time evolution of the concentrations [A] and [B] is described by the ordinary differential equations *d*[A]*/dt* = *−k*_*AB*_[A] + *k*_*BA*_[B] and *d*[B]*/dt* = *k*_*AB*_[A] *−k*_*BA*_[B] = *− d*[A]*/dt. Second order rate constants* quantify an elementary binding reaction of a reactant X with a molecule A to form a new molecule B

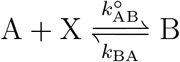

where 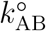 (units s^−1^M^−1^) and *k*_BA_ (units s^−1^) are the second-order forward and reverse rate constants. The time dependence is 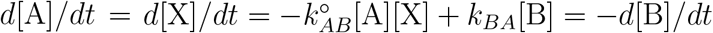. If the reactions do not change the concentration of X appreciably (i.e., *d*[X]*/dt≈* 0) then one can formally define the *pseudo-first-order rate constant*

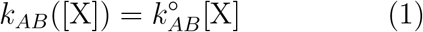

for the differential equation *d*[A]*/dt* = *− k*_*AB*_([X])[A] + *k*_*BA*_[B], which effectively corresponds to a first order reaction

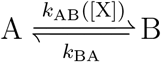

with a concentration-dependent forward rate “constant” *k*_AB_([X]). It is often possible to rewrite all second order reactions as pseudo-first order reactions and thus simplify the resulting system of differential equations.

### 2.2 Rate Constant Interdependence and Thermodynamic Consistency

When constructing kinetic diagrams the set of rate constants must be self-consistent.^15,38^ For *any* cycle in a kinetic model, the sum of the Gibbs free energy differences along the cycle equals zero in equilibrium (equilibrium cycle closure or Wegscheider condition ^39^)

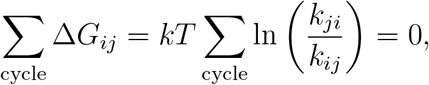

where the cycle is a closed loop in the kinetic diagram. The rate constants are first or pseudo first-order, where the pseudo first-order rates contain the second-order rates in Eq. 1. Combining logarithms and dividing through by *kT* yields

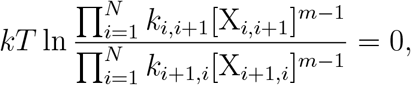

where *N* is the number of states in the cycle, X_*i,i*+1_ is the identity of the excess reactant in the reaction going from state *i* to *i* + 1, and *m* is the reaction order (e.g. *m* = 1 for first-order, *m* = 2 for second-order). It is implied that indices exceeding *N* are cyclic (i.e., *N* + *i* → *i*). First and second-order rates are considered *intrinsic* as they are separate from the chemical concentrations; thus for cycles with a chemical gradient, the terms can be separated out

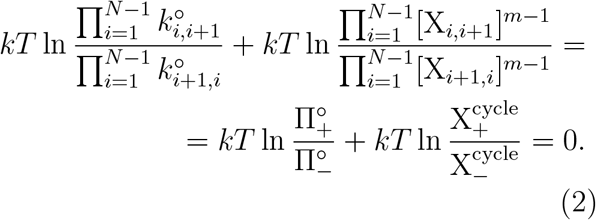

Here 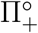 and 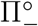 are the forward and reverse rate-products for the intrinsic rates in the cycle, respectively, and 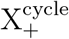 and 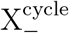 are the forward and reverse concentration products for the cycle, respectively. Note the ratio of the forward/reverse concentration products is equal to the equilibrium constant, 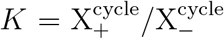. With the intrinsic system parameters separated, the remaining term is the chemical driving force for the cycle

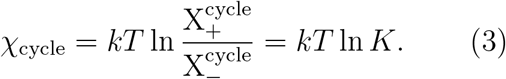

Expressing Eq. 2 in its simplest form,

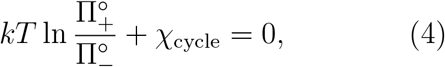

we can see that since *χ*_cycle_ is zero at equilibrium (since *K* is one at equilibrium), the remaining intrinsic term must hold equal to zero. Simplifying Eq. 4, the resultant expression (Hill’s kinetic cycle closure^14,15^)

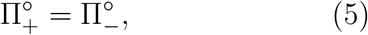

must hold for all cycles in a kinetic diagram under equilibrium and non-equilibrium conditions to remain consistent with thermodynamics. The intrinsic forward and reverse rate-products in any cycle must be equal because the intrinsic parameters do not contribute to the driving force of any cycle. Eq. 5 demonstrates that the rate constants in a kinetic model are interdependent and cannot be altered without due consideration. ^38^

### 2.3 Driving Kinetic Cycles

Kinetic cycles are driven by thermodynamic driving forces, which dictate the preferred direction of a cycle when present. The driving force *χ* (Eq. 3) is calculated from the difference of chemical potentials for a given ligand. In principle, other external driving forces such as electrostatic potential and mechanical forces can also be considered but for simplicity we restrict the following discussion to purely chemical potential (concentration) differences. For example, for a system involving some ligand M with inside and outside concentrations M_i_ and M_o_, the chemical potentials are related to the concentrations (in the infinite dilution limit)

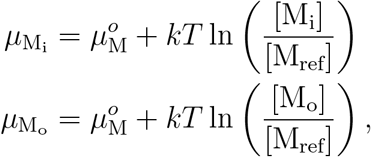

where 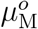 is the standard chemical potential for ligand M and M_ref_ is a reference concentration. The difference yields the thermodynamic driving force driving M from inside to outside:

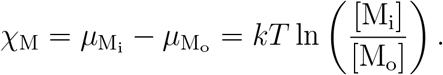

Cycles are only driven by external concentration gradients (chemical potential differences) and the internal rates cannot contribute as these cancel exactly due to Eq. 5.

In equilibrium all cycle driving forces vanish, *χ* = 0, due to the condition of detailed balance (all transition pairs in the system have equal forward and reverse fluxes between states, i.e., *k*_*ij*_*p*_*i*_ = *k*_*ji*_*p*_*j*_ where *p*_*i*_ is the probability to find the system in state *i*).^15^ In equilibrium the state probabilities remain constant, *dp*_*i*_*/dt* = 0. In general, out of equilibrium the cycle driving forces are non-zero and the state probabilities change with time. Steady state is a special non-equilibrium state where the state probabilities do not change (*dp*_*i*_*/dt* = 0) even though net fluxes between states exists with non-zero driving cycle forces *χ* ≠ 0. If the system can be described exclusively by first order and pseudo-first order reactions then the system will always evolve towards a fixed steady state at long times, regardless of initial conditions. The diagram approach then yields the steady state solution and thus characterizes the long-time behavior of the system described by the kinetic diagram.

### 2.4 State Probabilities from Kinetic Diagrams

In order to obtain the steady-state state probabilities for a kinetic diagram we start with the master equations, the set of linear differential equations which relate the change in probabilities *p*_*i*_ to the probability flux between states,

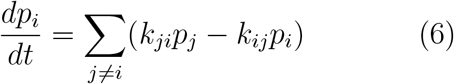

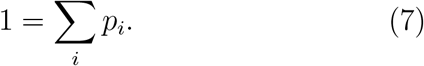

For a system with *N* states there are *N* master equations where *N* − 1 are linearly independent.^15^ The normalization condition, Eq. 7, is required to fully determine the system. The master equations for the 4-state model in Fig. 1 are included in Section 1.1 of the Supporting Information. At steady-state the master equations Eq. 6 reduce to a system of linear equations that can be solved directly with matrix methods. Alternatively, the master equations can be integrated using an ODE solver until a steady-state is reached. In both cases, a numerical solution for the steady state probabilities is obtained.

The diagram approach produces an exact algebraic solution. Broadly speaking, it requires the creation of sets of two types of diagrams: *partial diagrams* and *directional diagrams*. Partial diagrams are spanning trees of the kinetic graph and form the basis for constructing the directional diagrams, each of which are a special type of subgraph of their respective parent partial diagram. Because spanning trees are subgraphs of the original kinetic diagram it follows that all diagrams are also subgraphs of the kinetic diagram, i.e., in the process no edges are added that do not exist in the original problem.

#### 2.4.1 Partial Diagrams

For a kinetic diagram G with vertices V and edges E, the partial diagrams are the set of all spanning trees for G where each spanning tree is a minimally connected and maximally acyclic subgraph of G. In other words, each partial diagram contains all vertices V, has |E| = |V| −1 edges, and no closed loops. Each partial diagram has a unique configuration of edges resulting in a unique pathway connecting all vertices. As an example, the 4-state model in Fig. 1 has a total of 8 partial diagrams where each partial diagram contains all 4 vertices and 3 edges (Fig. 2). For more complex kinetic diagrams, the number of partial diagrams can be enumerated using Kirchhoff’s matrix theorem.^40^ Partial diagrams have no physical significance but rather serve as intermediates for constructing the directional diagrams covered in the next section.

**Figure 2:**
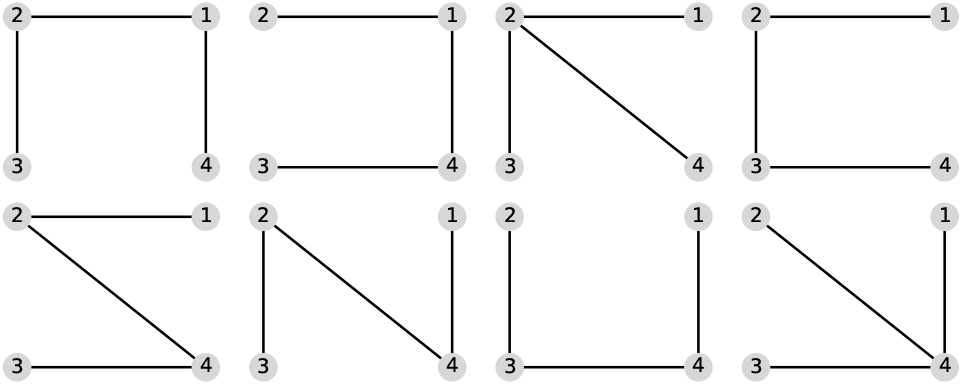
Partial diagrams for the 4-state model in Fig. 1. Partial diagrams are the set of all maximally connected and minimally acyclic undirected subgraphs (i.e., spanning trees) for the parent kinetic diagram. For the 4-state model there are |V| = 4 vertices, and thus each spanning tree contains |V| − 1 edges and no closed loops.

#### 2.4.2 Directional Diagrams

Directional diagrams are directed subgraphs of partial diagrams which represent the kinetic rate-products used to create the steady-state algebraic expressions for state probabilities and fluxes. For a partial diagram with vertices V (same as kinetic diagram) there are |V| child directional diagrams, where each directional diagram has a unique target state. Edges in directional diagrams represent rates, e.g., the edge from vertex (state) *i* to *j* represents rate *k*_*ij*_. Edges are oriented along the path to the target state creating |V| unique directed pathways along the kinetic diagram (per partial diagram). For example, the 4-state model in Fig. 1 has 8 partial diagrams and thus 32 directional diagrams (Fig. 3). The complete set of directional diagrams contains all unique directed pathways to all states in the kinetic diagram. Each directional diagram represents a unique rate-product that is formed by multiplying the rates for all edges in the diagram.

**Figure 3:**
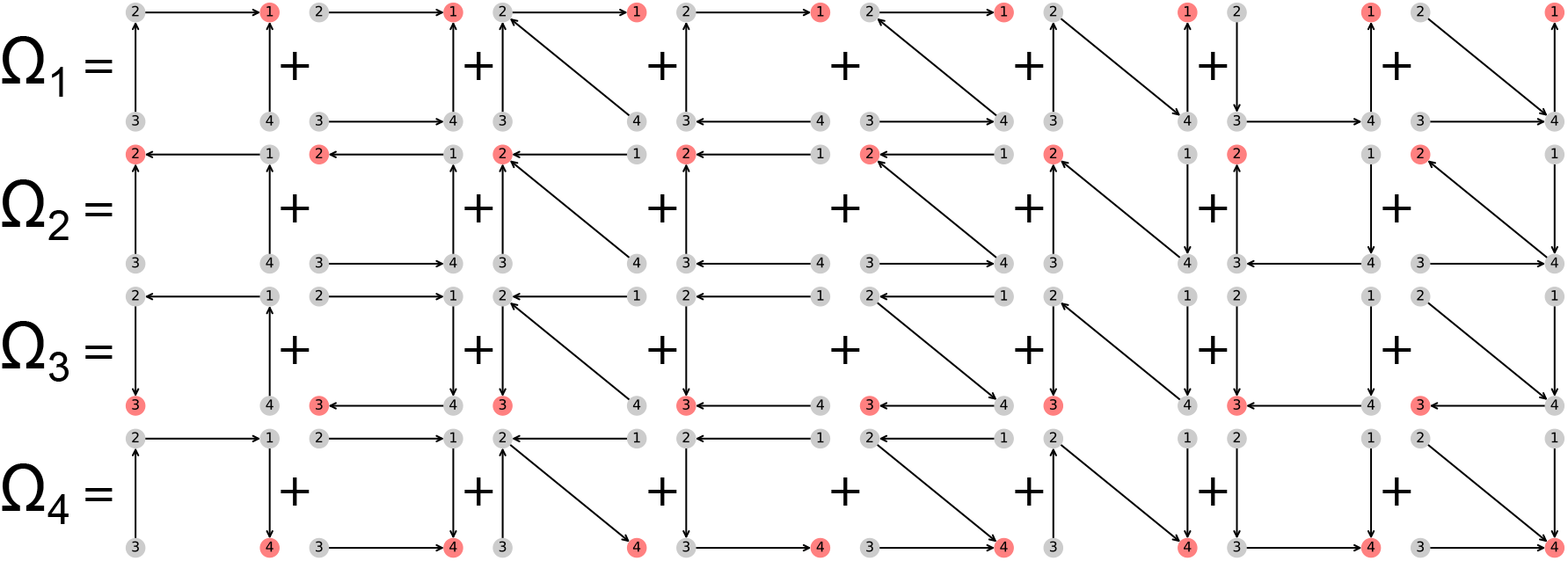
Directional diagrams for the 4-state model in Fig. 1. Directional diagrams represent rate-products used in steady-state expressions, where the rate-product is the product of all edge weights (i.e., kinetic rates) in the diagram. Directional diagrams are organized such that rows correspond to a single target state (colored coral) while columns contain the directional diagrams that share a parent partial diagram. The sum of each row yields the unnormalized expression for the corresponding state probability at steady-state, denoted Ω_*i*_, where the normalization factor is Σ = _*i*_ Ω_*i*_. Explicit rate-products are listed in Supporting Information.

The steady-state algebraic expressions for the state probabilities are created directly from the directional diagrams. Subsets of the directional diagrams are summed based on a common target state. For some state *i* the sum of directional diagrams (i.e., rate-products) with target state *i* yields the unnormalized state probability expression for state *i*,

Ω_*i*_ = ∑ directional diagrams for state *i*.

The state probabilities are

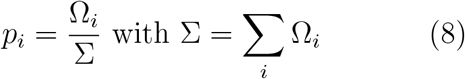

where the normalization factor Σ is the sum of all directional diagrams. For example, for the 4-state model in Fig. 1, the state probability expression for state 1 is found by summing the directional diagrams in the top row of Fig. 3 and dividing by the sum of all directional diagrams. Section 1.2 of the Supporting Information contains the complete set of unnormalized state probability expressions for the 4-state model.

### 2.5 Fluxes from Kinetic Diagrams

The diagram method produces expressions for different flux types in terms of the rates of the kinetic diagram. The three flux types, *transition fluxes, cycle fluxes*, and *operational fluxes* quantify different levels of steady-state activity in the system. The **cycle flux** is the most fundamental since all steady-state activity of the system can be accounted for in terms of cycle fluxes.^15^ The one-way cycle flux 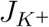 is the mean number of *K* cycles completed in the positive direction per second for any cycle *K* in the kinetic diagram. Cycle direction determinations follow from the chosen convention for the kinetic diagram (e.g. counter-clockwise cycles are positive), which must be consistent for all calculations.

*Net cycle fluxes* are the difference between the forward and reverse one-way cycle fluxes for cycle *K*,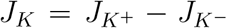. To create the exact expressions for the net cycle fluxes the flux diagrams and directional diagrams must be created. *Flux diagrams* are cycle-specific subgraphs of a kinetic diagram constructed from the target cycle and the pathways leading to that cycle. As an analogy, while directional diagrams represent every pathway to a given state, flux diagrams represent every pathway to a given cycle. Thus, while the value of a directional diagram is the rate-product of all pathways leading to a target state, the value of a flux diagram is the rate-product of all pathways leading to the target cycle (denoted Σ_*K*_) weighted by a cycle contribution. The cycle contribution is the difference between the forward and reverse rate-products (Π_+_− Π_−_) in the target cycle and determines the direction of the net cycle flux. Namely, *J*_*K*_ > 0 when Π_+_ > Π_−_. Thus, the net cycle flux for a cycle *K* is

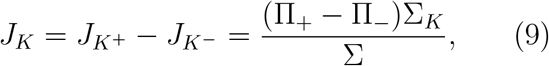

where (Π_+_− Π_−_)Σ_*K*_ is the sum of all flux diagrams for cycle *K* and Σ is the sum of all directional diagrams for the kinetic diagram (Eq. 8). For cycles where additional pathways cannot be drawn without creating an additional closed loop there are no flux diagrams, and thus Σ_*K*_ = 1.

As an example, in the 4-state model in Fig. 1 cycles *b* and *c* have two corresponding flux diagrams while cycle *a* has none (Fig. 4). Thus, Σ_*K*_ for cycles *b* and *c* are Σ_b_ = *k*_32_ + *k*_34_ and Σ_c_ = *k*_12_ + *k*_14_ while Σ_a_ = 1. Including the cycle contributions yields the net cycle flux expressions

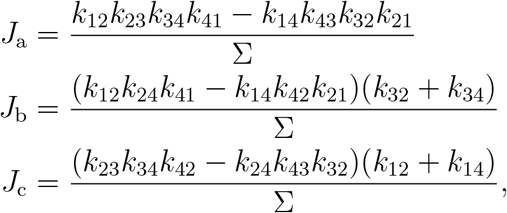

where Σ is the sum of all directional diagrams (Fig. 3).

**Figure 4:**
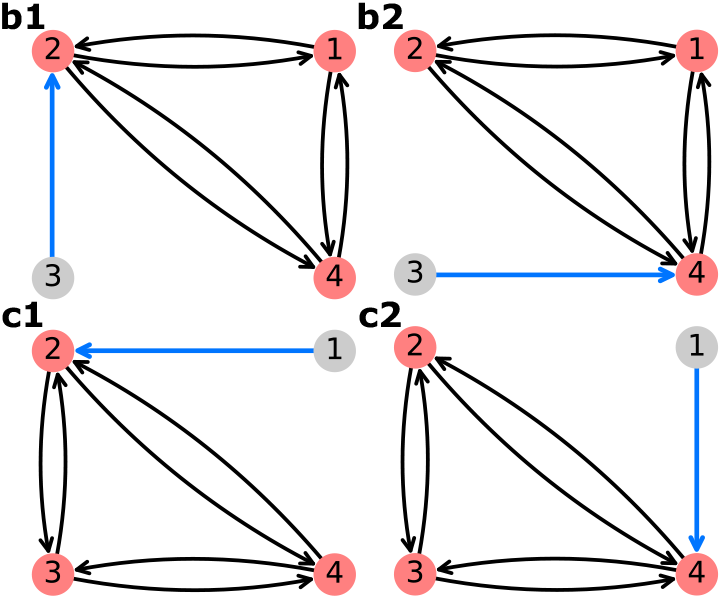
Flux diagrams for the 4-state model in Fig. 1. Flux diagrams are cycle-specific directed subgraphs of a kinetic diagram that represent expressions used to create steady-state net cycle flux expressions. Flux diagrams are composed of two parts: the target cycle from the original diagram (shown with coral-colored vertices) and pathways leading to the cycle (shown in blue). The flux diagram expression is the product of the non-cycle edge weights weighted by the difference of the forward and reverse rate-products in the target cycle. Flux diagrams with common target cycles (e.g. diagrams b1 and b2 or c1 and c2) are summed to create the unnormalized net cycle flux expressions, where the normalization expression derives from the sum of the directional diagrams for the kinetic diagram.

**Transition fluxes** are one-way probability fluxes for any transition *i* → *j* in the kinetic diagram and are the most local of the three flux types. The one-way transition flux is expressed *j*_*ij*_ = *k*_*ij*_*p*_*i*_ while the *net transition flux* is

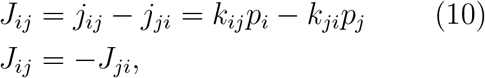

where *p*_*i*_ and *p*_*j*_ are the steady-state state probabilities. At equilibrium, *J*_*ij*_ = 0. Since net transition fluxes are expressed in terms of rates and state probabilities the exact expressions can be determined once the state probability expressions are defined.

Net transition fluxes can be calculated from net cycle flux expressions. For a given transition *i* → *j*, the net transition flux can be expressed as the sum of net cycle fluxes whose cycle traverses the transition *i* → *j*.^15^ For example, using the 4-state model in Fig. 1, the net transition flux for transition 2 → 4 is *J*_24_ = *J*_b_*− J*_c_. Similarly for 1 → 2, *J*_12_ = *J*_a_ + *J*_b_. Net cycle flux signs are determined by the chosen cycle convention. Since the counter-clockwise direction is positive for all cycles, the positive cycle direction for cycle *c* traverses 4 → 2 and thus requires a negative sign when calculating the net transition flux *J*_24_. While transition flux expressions can be expressed in terms of net cycle fluxes, the net cycle fluxes for all cycles cannot necessarily be deduced from the net transition fluxes because net cycle fluxes can outnumber net transition fluxes for complex kinetic diagrams.^15^

**Operational fluxes** correspond to specific processes in a kinetic diagram. Every operational flux has a conjugate thermodynamic driving force with the positive direction chosen the same for both. ^15^ Operational fluxes quantify individual processes in the kinetic model such as ligand transport, and thus can often be expressed in terms of net transition fluxes. The relationships between operational and transition fluxes are generally intuited from the diagram but can be found directly from the master equations evaluated at steady-state. ^15^ For example, considering the antiporter model in Fig. 1d, the net rate of appearance of substrate L on the outside (i.e., transition 3 → 4) is *J*_L_ = *J*_34_ (and also *J*_L_ = *J*_34_ = *J*_23_). For complex diagrams (i.e., diagrams with multiple processes corresponding to the thermodynamic driving force) several transition fluxes will contribute additively to the operational flux. ^15^

Operational fluxes can be expressed in terms of net cycle fluxes. The operational flux (in terms of net cycle fluxes) is found by summing the net cycle fluxes for any cycles with a net contribution to the given process. Continuing with the previous example, since positive completions of cycles a and c transport substrate L from inside to outside, the operational flux is *J*_L_ = *J*_a_ + *J*_c_. Similarly for the driving ion R, the operational flux is *J*_R_ = *J*_a_ + *J*_b_ where the positive direction moves the ion from the outside to the inside. More generally, for some ligand *x*

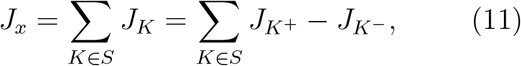

where *S* is the set of contributing cycles. Oneway operational fluxes are found by summing the corresponding one-way cycle fluxes (*J*_*K*_+, *J*_*K*_−) for all contributing cycles.

In summary, there are three different levels of interdependent fluxes with both one-way and net variants. Of the three flux types only transition and operational fluxes may be directly observable due to a change in state of the system. While individual net cycle fluxes are not generally observable, the set of net cycle fluxes contain all the steady-state information of the system and are thus capable of producing the expressions for any of the flux types. ^15^

### 2.6 KDA Algorithms

Here we describe the algorithms used by Kinetic Diagram Analysis to generate the partial, directional, and flux diagrams. All algorithms use graph objects from `NetworkX`,^16^ denoted G(V, E). V and E are the sets of vertices and edges in G, respectively, where each edge weight (i.e., kinetic rate *k*_*ij*_) is stored for later retrieval. Graph objects follow a principle of minimum required complexity. For example, partial diagrams are represented by `NetworkX.Graph` objects while directional and flux diagrams are represented by `NetworkX.MultiDiGraph` objects since they require directed edges. For each diagram type, the set of diagrams generated by `KDA` are converted into algebraic expressions (typically rate-products from the edges in the diagram) and combined together algebraically in accordance with the diagram method (i.e., Eqs. 8 and 9).

#### 2.6.1 Generating Partial Diagrams

With the goal of finding the set of all spanning trees, the `KDA` partial diagram algorithm uses what is broadly categorized as a “test and select” method^40^ where all subgraphs of the kinetic diagram are created and checked against the criteria of a spanning tree (Algorithm 1). To generate subgraphs for a kinetic diagram with vertices V, first the unique undirected edges within the kinetic diagram are found. This set of undirected edges is then used to create all possible combinations of |V|− 1 edges with the function `get_combinations_of`, which uses the built-in Python `itertools` module. Each edge combination yields a potential partial diagram that is subsequently checked against the spanning tree criteria. Spanning tree verification is carried out via the `is_spanning_tree` function, which uses `NetworkX`. Valid spanning trees are kept and passed along for directional diagram generation.

#### 2.6.2 Generating Directional Diagrams

To generate directional diagrams `KDA` implements an iterative algorithm over partial diagrams and their nodes (Algorithm 2). Directional diagrams are bidirectional subgraphs of partial diagrams where each partial diagram has |V| child directional diagrams. Thus the focus of the algorithm is finding the appropriate set of directional edges to include for each child directional diagram. We iterate over the partial diagrams and collect all paths from the leaf nodes (sources) to the target node.

Within the directional diagram algorithm, the pathways to each target node are found using `get_all_simple_path_edges` (Algorithm 3). This algorithm uses `NetworkX.all_simple_path_edges` which implements a modified depth-first-search algorithm to generate paths.^41^ Since directional diagrams always have a single target but potentially multiple sources, `all_simple_path_edges` is used in a reverse fashion to exploit its ability to handle multiple targets. The returned paths have their direction reversed and redundant edges removed since the paths may overlap.

##### Algorithm 1

`generate_partial_diagrams(G(V,` `E))`

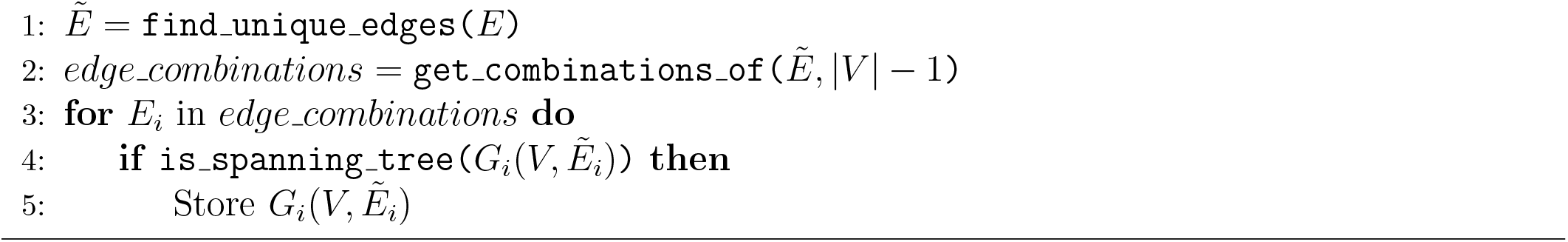

##### Algorithm 2

`generate_directional_diagrams(G(V,` `E))`

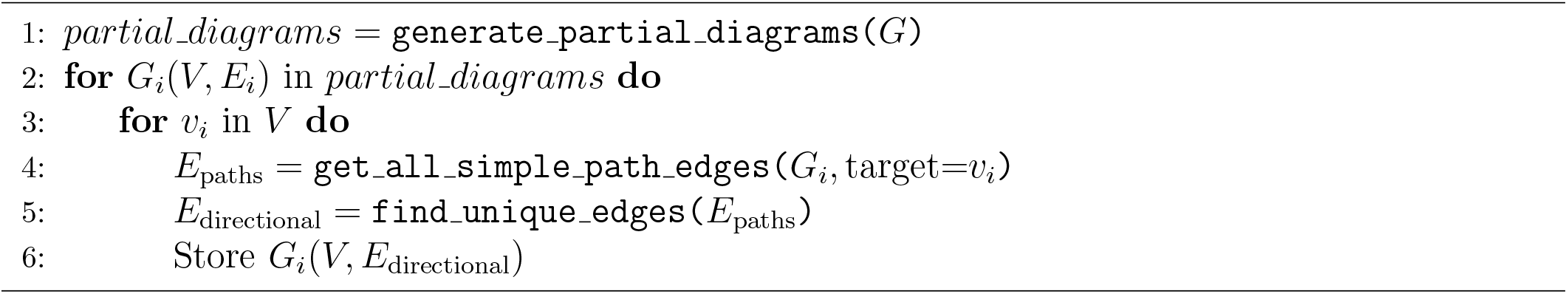

#### 2.6.3 Generating Flux Diagrams

Flux diagrams are used to generate net cycle flux expressions, i.e., Eq. 9. The process for creating flux diagrams is similar to the directional diagrams algorithm since both require finding pathways to a “target”. While directional diagrams have a target state, flux diagrams have a target cycle (i.e., a set of target states).

The `KDA` flux diagram algorithm (Algorithm 4) generates flux diagrams from an input kinetic diagram and a specified target cycle. The procedure for constructing a flux diagram for target cycle *K* and kinetic diagram G(V,E) is as follows. First the cycle edges must be isolated. Since flux diagrams contain only cycle *K*, the initial step is to categorize the edges E into two groups: cycle edges, which include edges for both cycle directions, and non-cycle edges, which are unidirectional. The cycle edges are preserved for later use whereas the non-cycle edges serve as the foundation for identifying potential edge paths.

To determine the pathways, first the number of directional edges for each pathway is calculated, |E_path_| = |V|−|V_cycle_|. The combinations of |E_path_| non-cycle edges are then used to create the possible pathways. Akin to Algorithm 1 a “test and select” approach^40^ is employed where each edge combination (i.e., pathway) is individually examined for validity. For each set of non-cycle edges the potential pathways to each target cycle node are aggregated. These pathway edges are checked to ensure they match the expected number of edges and to eliminate redundancy since pathways may overlap. If all edges are indeed distinct, a valid flux diagram was created and can be stored.

Within the flux diagram algorithm, the pathways to each target cycle node are found using Algorithm 5. To collect the pathways to each target cycle node from the combinations of non-cycle edges, the non-cycle edges are used to create pathways from the furthest neighbor of the target to the target node. For each target node, redundant edges are removed from the generated path and the resultant edges are kept for checking downstream in Algorithm 4.

Algorithm 6 was implemented to find the pathways from the furthest neighbor of the target node to the target node itself. First the non-cycle edge combinations are used by finding the edges that contain the target state. From this set of edges, the set of neighbors (within the set of non-cycle edges) are located. If neighbors are found, the non-adjacent edges are then located and used to create the set of edges with the appropriate direction (towards the target state). These edges are then stored, and the algorithm recursively iterates over each neighbor until no further neighbors are found.

##### Algorithm 3

`get_all_simple_path_edges(G(V,` `E),` `target)`

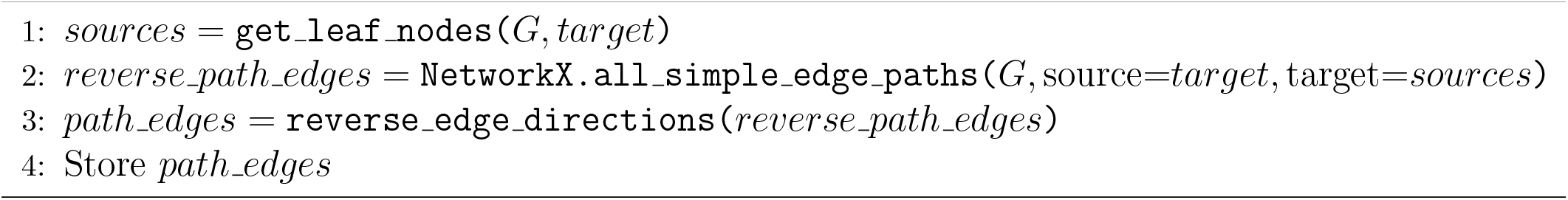

##### Algorithm 4

`generate_flux_diagrams(G(V,` `E),` `V`_cycle_ )

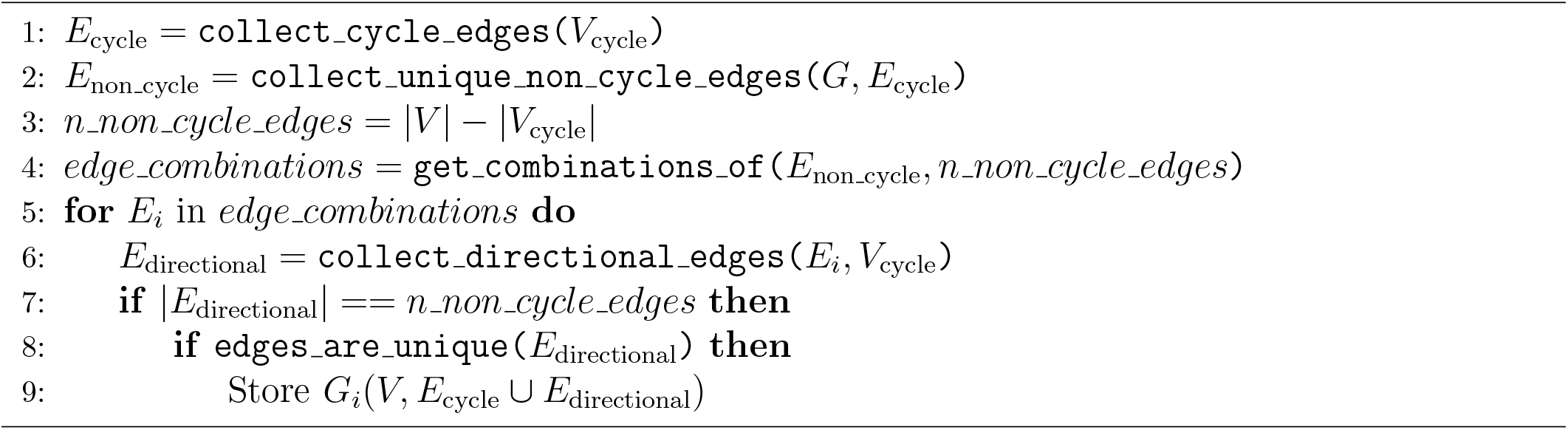

### 2.7 KDA Capabilities

`KDA` has methods to build and analyze kinetic diagrams. `KDA` uses `NetworkX`^16^ to construct diagrams, the `SymPy`^17^ library to construct and manipulate algebraic expressions, and both `NumPy`^42^ and `SciPy`^43^ for fast, vectorized array operations and numerical solving. `KDA` is written in Python 3 with continuous integration running a pytest-based test suite. The `KDA` tests run on Python 3.9 - 3.11 with 99% code coverage.

#### 2.7.1 Kinetic Diagram Generation

A typical workflow in `KDA` begins by defining the rate matrix for the kinetic model since it is used to construct the kinetic diagram (`NetworkX.MultiDiGraph` object). The rate matrix is similar to a connectivity matrix in structure but matrix elements represent the kinetic rate *k*_*ij*_ instead of vertex adjacency. While the rate matrix includes the edge weights, the connectivity matrix is sufficient to generate the kinetic diagram since edges are represented symbolically. Using Python code the kinetic diagram is generated by defining the connectivity or rate matrix as a `NumPy` array, creating the `MultiDiGraph` object, and using a `KDA` utility to create the edges (Fig. 5). The kinetic diagram serves as the central object in `KDA` since it enables the creation of the partial, directional and flux diagrams, as well as the state probability and net cycle flux expressions.

**Figure 5:**
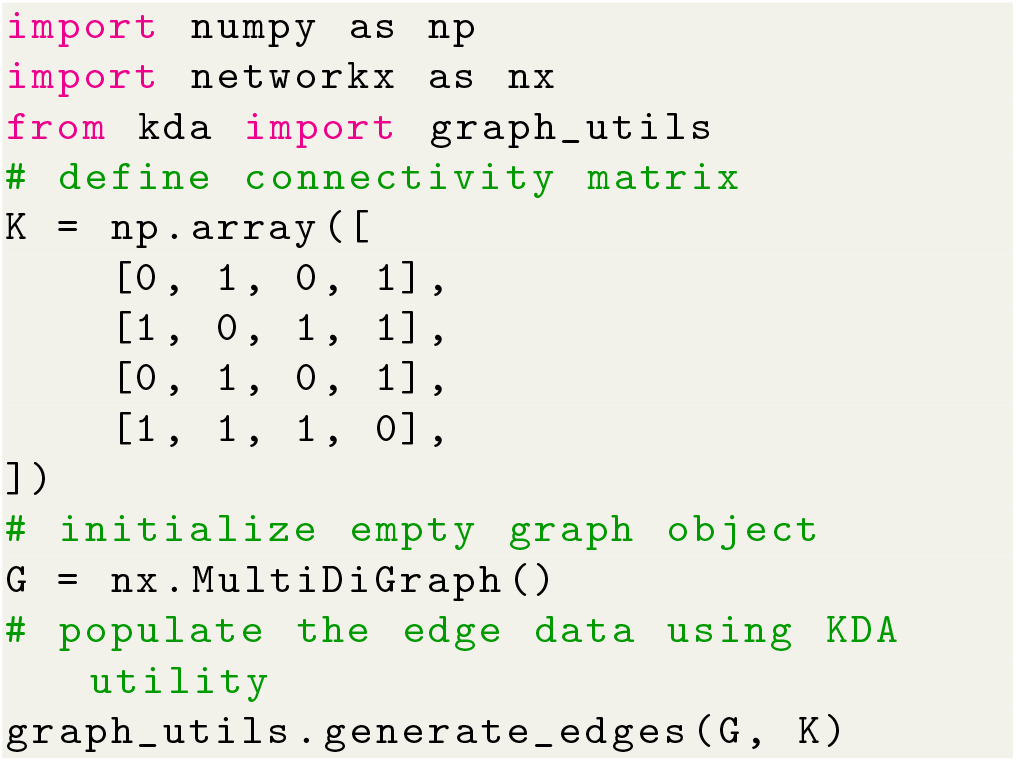
Python code for generating the kinetic diagram for the 4-state model (Fig. 1) using `KDA`.

##### Algorithm 5

`collect_directional_edges(E,` `V`_cycle_ )

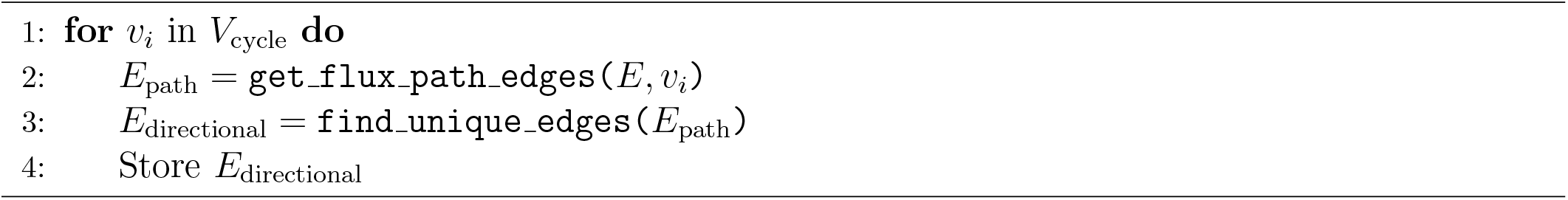

##### Algorithm 6

`get_flux_path_edges(E,` `v)`

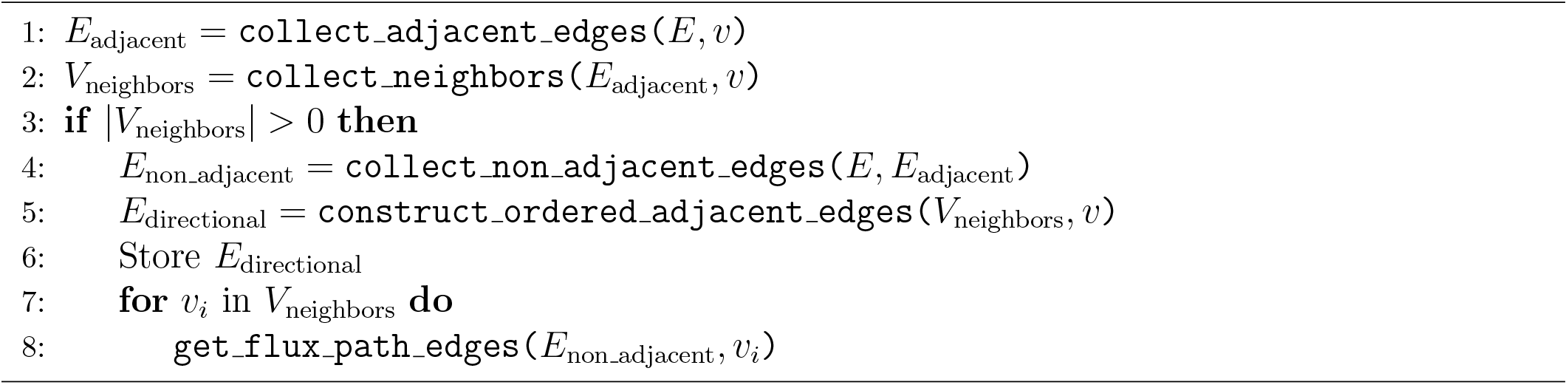

#### 2.7.2 Algebraic Expression Generation

Algebraic expressions generated by `KDA` are handled by `SymPy`^17^ enabling programmatic variable substitution and conversion to Python lambda functions. Starting with the code example in Fig. 5, the set of state probability expressions for the 4-state model can be generated with a single function call (Fig. 6). The state probability expression for state 1 is shown with the expression clipped due to its complexity. The output expressions for the 4-state model contain 8 terms in the numerator and 32 in the denominator, where the denominator is simply the sum of the numerators for all states in the kinetic diagram. The `KDA` function `calc_state_probs` generates the state probabilities as `SymPy` expressions by implementing Algorithm 2, in accordance with Eq. 8. For example, the probability for state 1 is

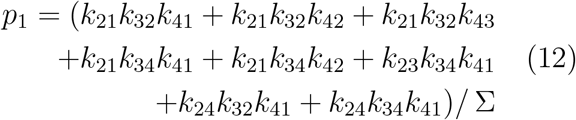

where Σ is the sum of all state multiplicities. The complete set of state multiplicity expressions can be found in Section 1.2 of the Supporting Information.

**Figure 6:**
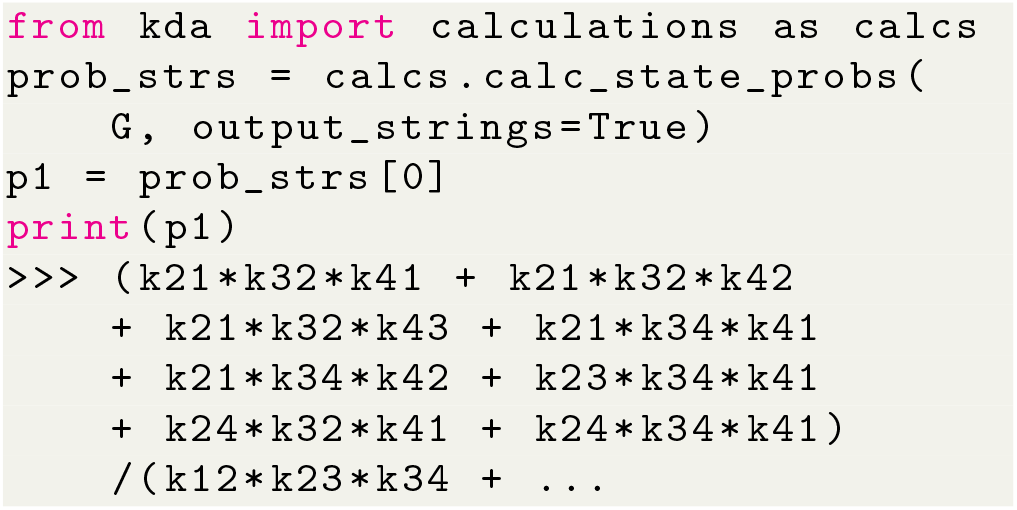
Python code for generating state probability expressions for the kinetic diagram in Fig. 1 using `KDA`. Graph object `G` is defined in Fig. 5.

With the expressions generated, `SymPy` is leveraged to perform variable substitutions and programmatic expression simplification. For example, using the 4-state antiporter model (Fig. 1d) we may set the concentration of ligand on inside and outside (L_int_, L_ext_) and driving ion (R_int_, R_ext_) and assume the same (symmetrical) binding rates (*L*_on_ for ligand and *R*_on_ for driving ion) and unbinding rates (*L*_off_, *R*_off_) on both sides of the membrane, and symmetrical leakage rates (*k*_24_ = *k*_42_ = *k*_leak_). We then apply these assumptions directly to our original expression by substituting the generic rates *k*_*ij*_ with appropriate expressions of our parameters of interest, as shown in Python code in Fig. 7; see Section 1.2 in Supporting Information for the mathematical rate expressions that correspond to the code shown here. With the variable substitutions applied to Eq. 12, the simplified state probability is now specific to the system:

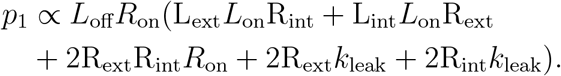

**Figure 7:**
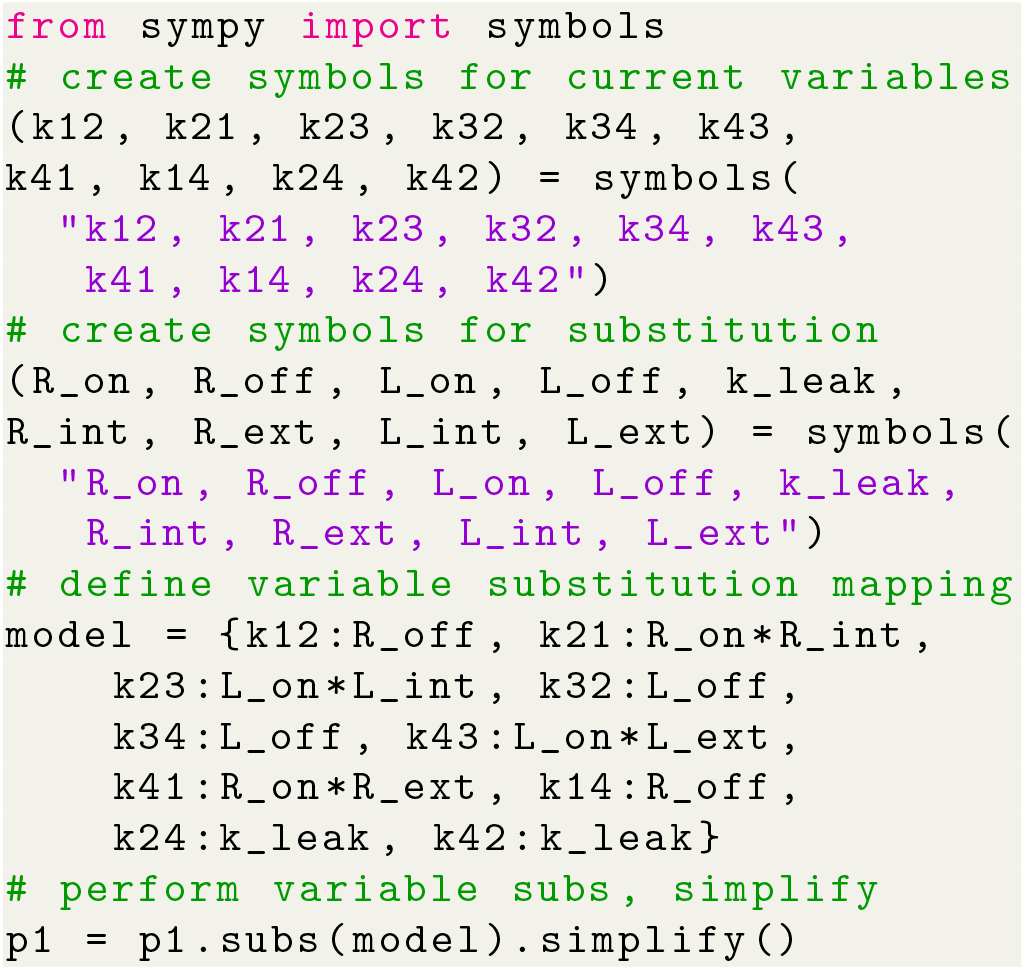
Python code for substituting state probability expression variables using `SymPy`. Initial state probability expression `p1` is defined in Fig. 6.

The expression is normalized with a similarly computed Σ (see Section 1.2 of the Supporting Information). For fast evaluation the simplified expression can be converted into a Python lambda function.

The same steps that were used for generating the state probability expressions (i.e., expression generation, substitutions, simplification) are also carried out to generate expressions for net cycle fluxes. Both state probabilities and net cycle fluxes require the use of the kinetic diagram object, but net cycle fluxes require a user-defined target cycle and cycle order (Fig. 8). The cycle is defined by a list of states (index zero) which can be listed using `KDA` utilities. The cycle order denotes the positive direction of the cycle by providing two of the cycle nodes in the order of the positive cycle direction. The `KDA` function `calculations.calc_net_cycle_flux` implements Algorithms 2 and 4 to construct the relevant diagrams and generate the `SymPy` expressions according to Eq. 9. The final expression for the net cycle flux of cycle *a* (using Fig. 8) is

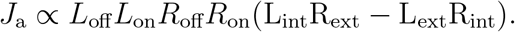

**Figure 8:**
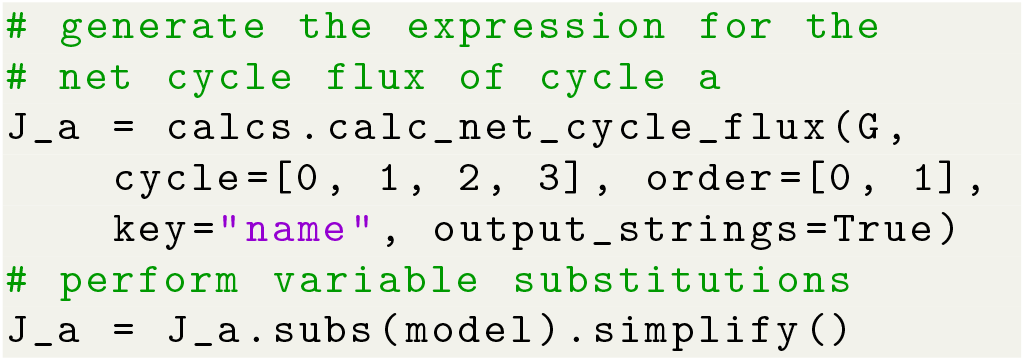
Python code for generating the net cycle flux expression for cycle a in Fig. 1 using `KDA`. Graph object `G` is defined in Fig. 5 and the model parameters `model` are defined in Fig. 7.

The expression is normalized using the same simplified expression as the state probabilities, included in Section 1.2 of the Supporting Information.

#### 2.7.3 Building Operational Flux Expressions

Of the expressions created with `KDA`, the operational flux expressions are the most difficult to create because they typically require multiple other flux expressions to be created beforehand. Like cycle and transition fluxes, the exact operational flux expressions can be created using `KDA`, but unlike these fluxes a manual approach is used to construct operational flux expressions because they rely on hand-picked cycles or transitions.

Starting with the *net cycle flux approach*, the algebraic expressions for the net cycle flux of each cycle have to be created and combined. This is accomplished by first determining which cycles contribute to the process of interest (e.g. ligand transport). In general, if a single cycle completion achieves the desired outcome (e.g. transports a ligand across the cell membrane) it is a contributing cycle. For example, continuing with the 4-state antiporter model (Fig. 1d), cycle *a* transports ligands R and L and thus is a contributor to both, while cycles *b* and *c* contribute to only ligand R and L transport, respectively. Focusing on the driving ion R, the desired outcome is the transport of R from outside to inside. Since positive cycle completions of both cycles *a* and *b* result in transport of R from outside to inside, using Eq. 11 the operational flux for R is *J*_R_ = *J*_a_ + *J*_b_. For substrate L the desired outcome is transport in the opposite direction (i.e., inside to outside), and thus *J*_L_ = *J*_a_ + *J*_c_. Thus, with the net cycle flux expressions defined the operational fluxes are constructed by simply summing the net cycle flux expressions for all contributing cycles, with careful consideration of cycle direction.

The *net transition flux approach* offers a simpler path to expression creation for kinetic models with multiple cycles but requires intuition of the kinetic model. Assuming the contributing transitions for the process of interest have been determined the expressions for the net transition fluxes for each transition must first be created. For a net transition flux *J*_*ij*_, the expressions for the state probabilities for both states *i* and *j* must be constructed using the process described in Section 2.7.2, then combined according to Eq. 10. The final expression for the operational flux is found by summing the net transition fluxes for the contributing transitions (Fig. 9). For example, in the 4-state antiporter model transitions 4 → 1 and 1 → 2 are the contributing binding/unbinding transitions for ligand R. Thus, the operational flux for R is expressed in terms of either transition *J*_R_ = *J*_4,1_ = *J*_1,2_. Similarly for substrate L, the relevant transitions are 2 → 3 and 3 → 4, therefore *J*_L_ = *J*_2,3_ = *J*_3,4_.

**Figure 9:**
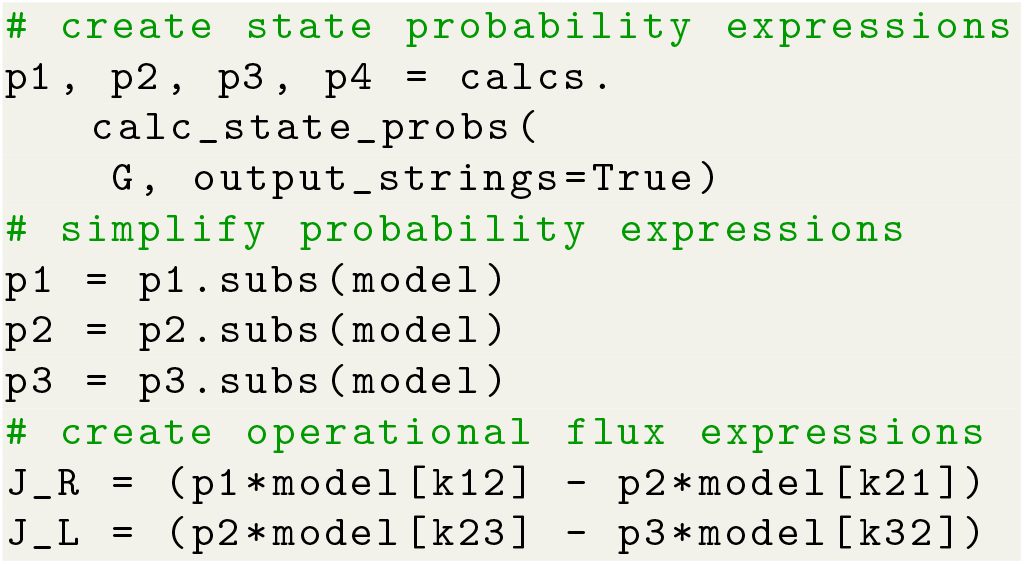
Python code for generating the operational flux expressions in terms of transition fluxes for ligands R and L for the 4-state antiporter model (Fig. 1d). The operational fluxes for R and L are *J*_R_ = *p*_1_*k*_12_ *− p*_2_*k*_21_ and *J*_L_ = *p*_2_*k*_23_*− p*_3_*k*_32_, respectively. The graph object `G` is defined in Fig. 5 and the model parameters `model` are defined in Fig. 7.

In summary, the exact operational flux expressions can be found via `KDA` using either net cycle fluxes or net transition fluxes. The net cycle flux approach requires knowledge of the contributing cycles while the net transition flux approach requires the knowledge of the contributing transitions. The net cycle flux approach requires the generation of net cycle fluxes for all contributing cycles while the net transition flux approach only requires state probability expressions for the states of interest. For complex models the net cycle fluxes may be cumbersome since many cycles will have to be inspected. For either method, the resultant operational flux expressions can be simplified and converted to Python lambda functions for fast iteration through parameter sets.

#### 2.7.4 Plotting Diagrams and Cycles

`KDA` uses the `Matplotlib`^44^ and `NetworkX`^16^ libraries for creating diagram and cycle figures. The kinetic, partial, directional, and flux diagrams can be plotted using `KDA` utilities, as well as cycles. Diagrams and cycles can be plotted individually or as panels, with features for highlighting nodes and cycles of interest. Diagrams can be displayed with bidirectional arrows or separate arrows based on user preference.

#### 2.7.5 Numerical Solvers

`KDA` has methods to resolve the steady-state state probabilities numerically for validation and testing purposes. These numerical solvers operate on the kinetic matrix to programmatically generate the set of master equations and solve them using singular value de-composition, ODE integration, and direct matrix solving. The ODE solver implements `SciPy.integrate.solve_ivp` with the LSODA integration method^43,45,46^ to solve for *p*_*i*_(*t*) since chemical kinetic differential equations are commonly stiff.^47^ For the matrix solver, the `NumPy`^42^ function `NumPy.linalg.svd` is used to find the linearly dependent differential equation from the kinetic differential equations, at which point it is replaced with the probability normalization equation (Eq. 7). The final matrix is solved using `NumPy.linalg.solve`.

### 2.8 Data Sharing and Software Used

In this work we used Python 3.9 with the following packages: `NetworkX`^16^ version 3.2.1, `SymPy`^17^ version 1.12, `NumPy`^42^ version 1.25.2, `SciPy`^43^ version 1.11.2, `Matplotlib`^44^ version 3.7.2, and `KDA` version 0.3.0. All packages were installed from PyPi with the exception of `KDA` which was installed from source. Version 1.1 of the `KAPattern`^27^ Windows application was downloaded from https://vpr.sites.uofmhosting.net/software/kapattern.

The source code for Kinetic Diagram Analysis can be found on GitHub at github.com/Becksteinlab/kda and is archived under DOI 10.5281/zenodo.5826393. The code and data for all figures is located in the `KDA` examples repository. The `KDA` examples repository can be found at github.com/Becksteinlab/kda-examples and is archived under DOI 10.5281/zenodo.6437043.

## 3 Results and discussion

We first validate `KDA` and compare its performance to direct numerical calculations. Using two applications from the biophysics of secondary active transporters we demonstrate the potential of the diagram method to obtain exact expressions for the relevant observables, namely operational fluxes. We conclude with a discussion of an alternative approach to calculate operational fluxes.

### 3.1 KDA Validation and Performance

We validated and benchmarked `KDA` on a set of 875 randomly generated kinetic diagrams of varying degree and order. All kinetic diagram data were collected using a Windows 10 PC with an AMD Ryzen 3900X CPU and G.SKILL Ripjaws V Series DDR4 RAM (2×32GB).

For each random diagram, we ensured its thermodynamic validity by applying the following checks: (1) The kinetic diagram used a thermodynamically consistent set of rates at equilibrium that were generated with the `MultiBind`^38^ library. (2) Since the net transition and net cycle fluxes are zero under equilibrium conditions, the net cycle flux and net transition flux for all cycles and transition pairs, respectively, were calculated and checked to be numerically close to zero. Operational fluxes were excluded since they are necessarily zero if all other net fluxes are zero. (3) Lastly, the number of generated partial and directional diagrams were checked against the expected values calculated using Kirchhoff’s matrix theorem.^40^

#### 3.1.1 Validation

With the set of random diagrams, two validation analyses were performed. (1) We compared `KDA` state probability expressions against the results from the `KDA` matrix solver (detailed in Section 2.7.5) and (2) we directly compared `KDA`-generated symbolic expressions to expressions produced with the `KAPattern` software.^27^ To quantify the discrepancy between `KDA` and matrix solver numerical results, the accuracy was quantified by calculating the root-meansquare deviations (RMSD) between the `KDA` state probabilities and the matrix solver probabilities as

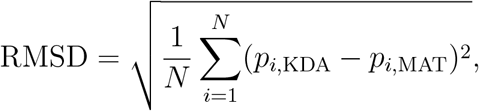

where *N* is the number of states in the diagram. For the comparison to `KAPattern`, five of the randomly generated diagrams (a 3-state, 4-state, 5-state, 6-state, and 7-state) were ran through both `KDA` and `KAPattern` and the unnormalized state probability expressions were directly compared for each state using `SymPy`.^17^ To compare the accuracy of the state probability calculations across graphs of varying degree the RMSD values for all kinetic diagrams were grouped by graph degree and placed in box and whisker plots highlighting the minimum, maximum, and median RMSD values on a number of states basis. As the number of states increases, the median and maximum RMSD values trend upward from 10^−16^ and 10^−13^ for 3-state case to 10^−14^ and 10^−11^ for the 20-state case, respectively (Fig. 10a). For the 3-state diagrams, the median RMSD is close to machine floating point precision (i.e., 10^−16^) and thus numerically indistinguishable from zero, while the max RMSD is relatively larger. All generated 3-state diagrams are identical and the state probability expressions are known since they are easily verified by hand. Thus, since the `KDA` 3-state solution is known to be correct by comparison against the manually calculated solution, the relatively larger maximum RMSD is likely attributable to the pseudoinverse calculation in the matrix solver. The trends of increasing median and maximum RMSD are further explained by the increased number of floating point operations required to evaluate the state probability expressions numerically. As graph degree increases the number of directional diagrams increases, and thus the number of floating point number operations is increased. Furthermore, RMSD values are dominated by the largest difference between a pair of *p*_*i*_ values so the resultant RMSD is approximately equal to the largest probability difference between methods. Thus, since the largest RMSD value for any diagram is 10^−11^ the `KDA` state probabilities for all kinetic diagrams agree with the matrix solver to better than 10^−11^. Overall `KDA` and the matrix solver solutions show good agreement and are in line with expectations for numerical evaluations.

**Figure 10:**
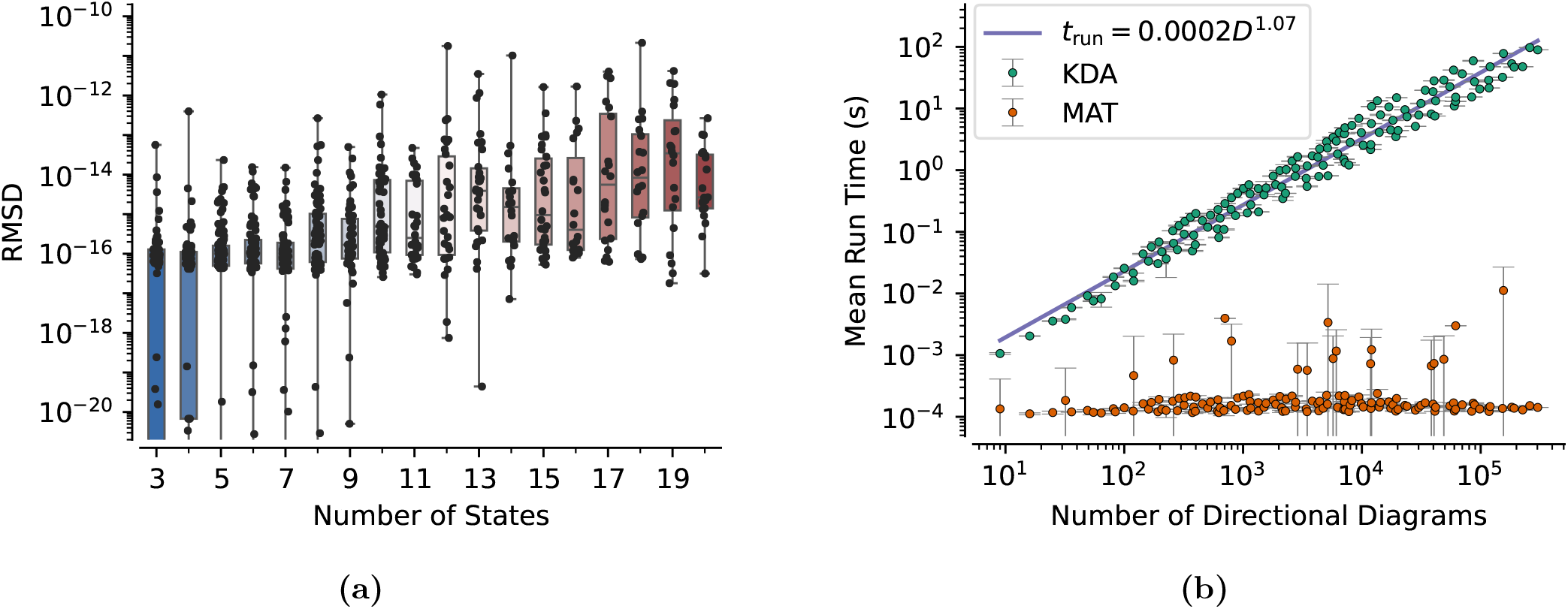
`KDA` State Probability Validation and Performance. (a) Box and whisker plots of the root-mean-square deviations for the `KDA` and matrix solution state probabilities for a range of graph degrees. Points are shown for every generated kinetic diagram. Boxes include 50% of the data while whiskers indicate the minimum and maximum RMSD values. (b) Average run time for the Kinetic Diagram Analysis (KDA) and matrix (MAT) solution state probability calculations as a function of the number of directional diagrams required to generate the state probability expressions. Both methods yield a numerical solution starting from a common rate matrix, where the `KDA` run time includes the time to generate all relevant diagrams and expressions. The average is calculated for sets of random diagrams with identical numbers of directional diagrams, where the standard deviation of the set is calculated for error. The fit line gives the average run time *T* as a function of the number of directional diagrams *D* for the `KDA` solution.

Since numerical calculations have inherent error due to floating point number calculations, the `KDA` and `KAPattern` symbolic expressions were directly compared using the `SymPy` computer algebra system.^17^ Since the normalization factor is a sum of the state multiplicities (Eq. 7), the state multiplicity for each state was directly compared. `SymPy` proved the expressions 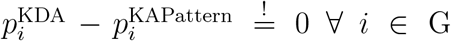 for the 3-state, 4-state, and 5-state diagrams, i.e., `KDA` and `KAPattern` yielded identical symbolic expressions. For the 6 and 7-state diagrams, disagreements were found between the `KDA` and `KAPattern` expressions. In these cases, the `KAPattern` expressions contained invalid rate-products that included either an additional rate, mutually exclusive rates (e.g. the forward and reverse rates for a given transition), or both (see Section 4 of the Supporting Information for details).

Overall, the `KDA` state probability expressions were shown to be numerically correct for different typical inputs and agreed with the symbolic expressions from `KAPattern` for the cases where `KAPattern` did not produce incorrect output.

#### 3.1.2 Performance

In conjunction with the accuracy analysis, `KDA` performance was separately measured against the matrix solver and `KAPattern`. For the matrix solver comparison, the run times for `KDA` and the matrix solver were recorded, where the `KDA` run times include the generation of underlying diagrams, expressions, and numerical evaluation. `KDA` performance is measured as a function of state probability expression complexity, where expression complexity is quantified by the number of directional diagrams generated. The number of directional diagrams is used since the number of rate-product terms in a given expression scales with the number of directional diagrams. Namely, for a kinetic diagram with *N* states and *D* directional diagrams, the number of terms in a state probability expression is 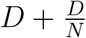. For the `KAPattern` analysis, `KDA` performance was measured at run time while the `KAPattern` run time was measured using a stopwatch.

For the matrix solver comparison, the average run times for `KDA` and the matrix solver were calculated for graphs with equivalent numbers of directional diagrams. As the number of directional diagrams increases, matrix solver performance is nearly constant while `KDA` performance increases nearly linearly (Fig. 10b). Notably, the `KDA` performance trends with the relationship *t*_run_ = 0.0002*D*^1.07^ where *t*_run_ is the run time and *D* is the number of directional diagrams. Despite the matrix solver outperforming `KDA` across the range, the average run times for `KDA` remained below one second until the number of directional diagrams exceeded 2000. For reference, the 4-state antiporter model (Fig. 1d) has only 32 directional diagrams and thus requires approximately 8.16*×* 10^−2^ s to generate and evaluate the expressions. It is worth noting that the vast majority of the `KDA` run time is spent generating the diagrams and expressions, whereas expression evaluation requires less than 1 millisecond for most models. A typical `KDA` workflow only requires expressions to be created once and thus subsequent evaluations would be in the same run time regime as the matrix solver, allowing for efficient parameter searches for a given kinetic model.

For the `KAPattern` performance comparison, `KAPattern` required less than 0.5 s to generate the expressions for all kinetic diagrams (3-7 states) while `KDA` required 0.17 s and 22.93 s for the 3 and 7-state kinetic diagrams, respectively. The superior `KAPattern` performance is likely due to the use of more efficient algorithms, as well as a known `KDA` bottleneck where expression parsing with `SymPy` is consuming the majority of the run time for symbolic outputs (93% for the 7-state diagram). This bottleneck will be addressed in future versions of `KDA`. Regarding algorithmic improvements, the `KDA` algorithms for generating partial and directional diagrams (Algorithms 1 and 2) contain a loop over possible edge combinations, which roughly scale as *N*! for kinetic diagrams with *N* states. This approach leads to increased run time since there are many edge combinations that create invalid subgraphs, resulting in unnecessary iterations. Other algorithms are available which could avoid or reduce these unnecessary iterations.^40^ Algorithms developed by Char, ^48^ Sen Sarma et al.,^49^ Naskar et al., ^50^ or Onete^51^ offer alternative “test and select” methods, but our method is a proof of concept and sufficiently fast for our applications.

### 3.2 Applications

Here we apply the diagram method to increasingly complex models to quantify changes in system function using steady-state fluxes. For each model the steady-state fluxes are calculated using `KDA` and expressed in terms of select internal system parameters. Internal parameters are then varied to observe changes in system function (i.e., rate biasing). We begin with a 6-state antiporter model similar to the model studied by Berlaga and Kolomeisky ^37^ to demonstrate leakage effects on system function by varying a single parameter, the leakage rate. We then use the 8-state free exchange model studied by Hussey, Thomas and Henzler-Wildman^36^ to observe the effects of rate biasing on transporter phenotype using two different approaches: varying alternating access rates, and varying ligand unbinding rates. We complete our analysis by demonstrating how the results of some of our applications can be found more simply using net transition fluxes.

We note that in the following we only consider kinetic diagrams that contain edges that are part of cycles, i.e., we do not include “dangling nodes” or “kinetic trap states” that are only connected by a single reversible reaction to another node. Adding such a node generally affects the magnitude of cycle fluxes and thus operational fluxes, as shown explicitly for a simple 3-state model in Section 2 in Supporting Information. Even though we do not show examples beyond the 3-state model, the diagrammatic method and `KDA` can also be used to exactly solve the steady state of such systems.

#### 3.2.1 Effective Leakage in a Sodium Proton Antiporter

To demonstrate the potential of the diagram method with `KDA`, we used two similar kinetic models to investigate the impact of introducing a leakage transition on ligand turnover and transport stoichiometry. These models closely resemble the model studied by Berlaga and Kolomeisky. ^37^ Serving as the control case is a generic single-cycle 6-state antiporter model. In comparison, the second model incorporates an additional leakage transition resulting in a total of three kinetic cycles. The detailed kinetic diagram for both models is shown in Fig. 11a with the individual diagrams highlighted in Fig. 11b.

**Figure 11:**
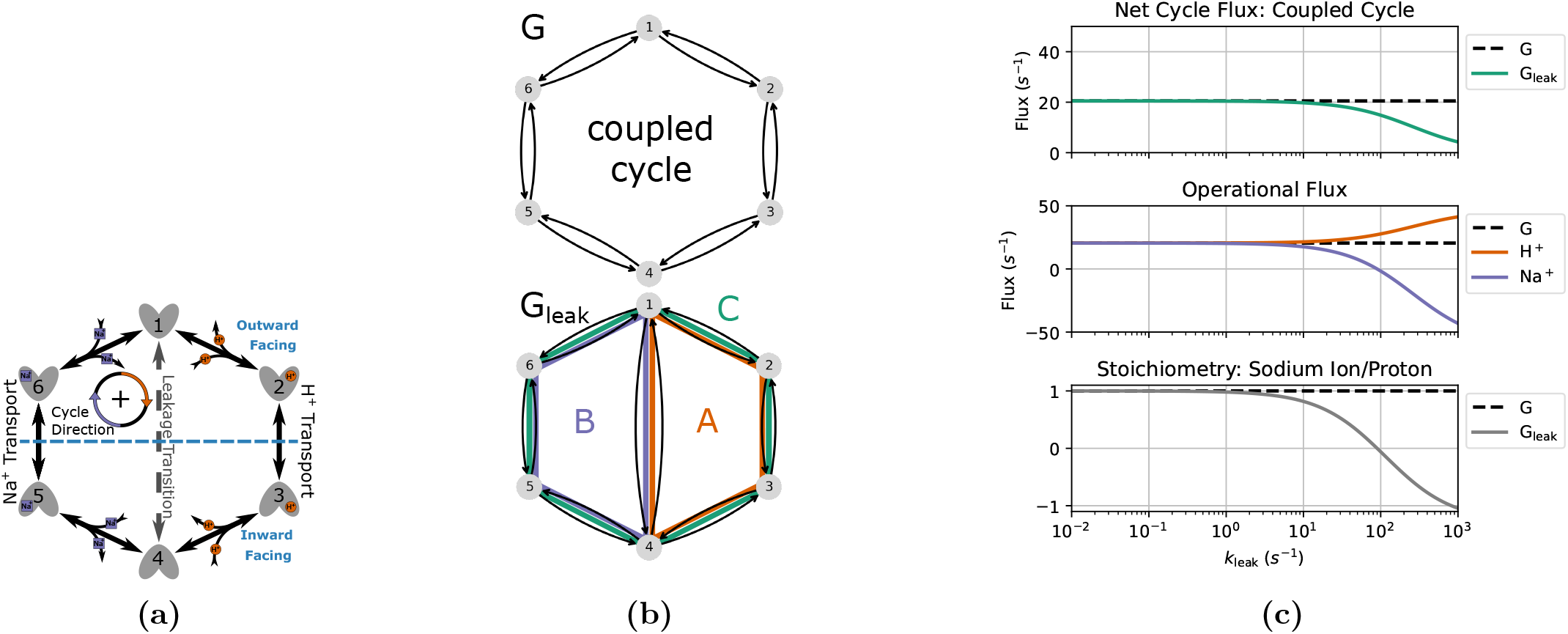
Effects of Leakage on a Generic 6-state Antiporter Model. (a) Kinetic model for a sodium proton antiporter. The dominant Hamiltonian cycle direction is clockwise. (b) 6-state diagram (top) and 6-state diagram with a leakage transition (bottom). Sodium and proton leakage cycles B and A are labeled in lavender and orange, respectively, with the coupled cycle shown in green. (c) Top to bottom: net cycle fluxes for the coupled cycle of both 6-state models, operational fluxes for sodium ions and protons, and the stoichiometry (sodium ion per proton transported), all as a function of the leakage transition rate.

The control model, G, contains only one cycle with a locked 1:1 stoichiometry. This results in the operational flux for both ligand species being identical to the net cycle flux for the coupled cycle (i.e.,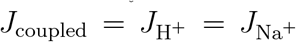). The leakage model, G_leak_, contains three cycles: A, B and C. Cycles A and B are leakage cycles, spontaneously transporting ligands down their concentration gradients. Cycle C is the same coupled cycle in G. Thus, the operational fluxes for G_leak_ are

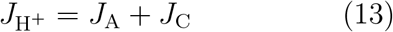

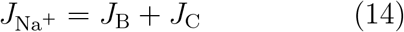

The coupled cycle is defined such that a positive cycle completion results in the transport of a single sodium ion from inside (intracellular) to outside (extracellular), and a proton from outside to inside. The sodium and proton gradients are both inward-facing where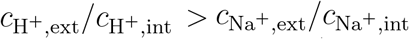, resulting in the clockwise transitions (i.e., positive cycle completions) being dominant for the coupled cycle.

A series of assumptions were made for both models to simplify calculations: proton and sodium ion binding and unbinding rates are assumed uniform for all reactions and the voltage dependence is ignored, conformational change (alternating access) rates are separately symmetric for sodium and proton transport processes, and the leakage transition rates are symmetric (Table 1). Proton and sodium binding rate estimates are based on the sodium proton antiporter NHA2 due to its electroneutral transport and experimental data availability. Binding rate estimates based on NHA2 are only used to provide a set of realistic parameters for the 6-state model and not to represent a full model of NHA2. The leakage rate, *k*_leak_, is symmetric (i.e., *k*_1,4_ = *k*_4,1_) and varied to show the effect of increasing the transition probability for the leakage transition.

**Table 1:** Rate Constant Definitions for Sodium Proton Antiporter Model.

| Parameter | Transition | Process | Value | Unit | Source |
| --- | --- | --- | --- | --- | --- |
| $k_{\text{on,H}^+}^{\circ}$ | $1 \rightarrow 2$ | Proton-on | $1.120 \times 10^{10}$ | $\text{M}^{-1}\text{s}^{-1}$ | Estimate<sup>a</sup> |
| | $3 \leftarrow 4$ | Proton-on | $1.120 \times 10^{10}$ | $\text{M}^{-1}\text{s}^{-1}$ | Estimate<sup>a</sup> |
| $k_{\text{off,H}^+}$ | $1 \leftarrow 2$ | Proton-off | $1.775 \times 10^3$ | $\text{s}^{-1}$ | Estimate<sup>b</sup> |
| | $3 \rightarrow 4$ | Proton-off | $1.775 \times 10^3$ | $\text{s}^{-1}$ | Estimate<sup>b</sup> |
| $k_{\text{on,Na}^+}^{\circ}$ | $4 \rightarrow 5$ | Sodium-on | $3.011 \times 10^9$ | $\text{M}^{-1}\text{s}^{-1}$ | Estimate<sup>a</sup> |
| | $6 \leftarrow 1$ | Sodium-on | $3.011 \times 10^9$ | $\text{M}^{-1}\text{s}^{-1}$ | Estimate<sup>a</sup> |
| $k_{\text{off,Na}^+}$ | $4 \leftarrow 5$ | Sodium-off | $9.033 \times 10^7$ | $\text{s}^{-1}$ | Estimate<sup>c</sup> |
| | $6 \rightarrow 1$ | Sodium-off | $9.033 \times 10^7$ | $\text{s}^{-1}$ | Estimate<sup>c</sup> |
| $k_{\text{AA,H}^+}$ | $2 \rightarrow 3$ | Alternating access | 69 | $\text{s}^{-1}$ | Ref. 52 |
| | $2 \leftarrow 3$ | Alternating access | 69 | $\text{s}^{-1}$ | Ref. 52 |
| $k_{\text{AA,Na}^+}$ | $5 \rightarrow 6$ | Alternating access | 350 | $\text{s}^{-1}$ | Ref. 52 |
| | $5 \leftarrow 6$ | Alternating access | 350 | $\text{s}^{-1}$ | Ref. 52 |
| $k_{\text{leak}}$ | $1 \rightarrow 4$ | Leakage | $10^{-2} - 10^2$ | $\text{s}^{-1}$ | Range<sup>d</sup> |
| | $1 \leftarrow 4$ | Leakage | $10^{-2} - 10^2$ | $\text{s}^{-1}$ | Range<sup>d</sup> |
| $\text{pH}_{\text{ext}}$ | $1 \rightarrow 2$ | Proton-on | 5.5 | | Ref. 53 |
| $\text{pH}_{\text{int}}$ | $3 \leftarrow 4$ | Proton-on | 8.5 | | Ref. 53 |
| $c_{\text{Na}^+, \text{ext}}$ | $6 \leftarrow 1$ | Sodium-on | 150 | mM | Ref. 53 |
| $c_{\text{Na}^+, \text{int}}$ | $4 \rightarrow 5$ | Sodium-on | 10 | mM | Ref. 53 |

<sup>a</sup> Calculated by estimating the flux through a disk, $k_{\text{on},X}^{\circ} = 4RD_X N_A$ .<sup>54</sup> $R$ , the radius of the disk, was given an approximate value of $5\text{\AA}$ using the crystal structure of NHA2. The diffusion coefficients, $D_X$ , are $9.3 \times 10^{11} \text{\AA}^2 \text{s}^{-1}$ and $2.5 \times 10^{11} \text{\AA}^2 \text{s}^{-1}$ for $\text{H}^+$ and $\text{Na}^+$ , respectively.<sup>55,56</sup> Final values are calculated using $N_A$ , the Avogadro constant, and conversion factor $1 \text{ liter} = 10^{27} \text{\AA}^3$ .
<sup>b</sup> Calculated $k_{\text{off}} = k_{\text{on}}^{\circ} 10^{-\text{p}K_a}$ , with a $\text{p}K_a$ of 6.8.<sup>52</sup>
<sup>c</sup> Calculated $k_{\text{off}} = k_{\text{on}}^{\circ} K_D$ , with a $\text{Na}^+$ binding affinity $K_D = 30 \text{ mM}$ .<sup>53</sup>
<sup>d</sup> Rates are varied in accordance with the thermodynamic consistency condition, Eq. 5.

To observe the effects of leakage cycles on the 6-state antiporter model (Fig. 11a) the steady-state operational fluxes (i.e., Eqs. 13-14) were calculated as a function of the leakage rate *k*_leak_ with fixed substrate concentration gradients (Table 1). The expectation was to observe efficient antiport behavior (i.e., 1:1 stoichiometry) for the control model and an overall reduction in transport efficiency for the leakage model as the leakage rate is increased.

For the control model G positive fluxes for both ligands are observed with 1:1 stoichiometry (Fig. 11c). With only a single coupled cycle available the stoichiometry is locked 1:1 resulting in the expected ideal antiport behavior.

For the leakage model G_leak_, as the leakage rate is increased we observed an increase in proton flux and decrease in sodium flux (relative to the control model) resulting in an overall reduction in stoichiometry (Fig. 11c). These flux changes are characteristic of the introduction of a leakage cycle, where each ligand is now able to effectively bypass the productive (i.e., coupled) cycle and flow in accordance with their respective concentration gradients, decreasing the efficiency of the transporter. As the leakage rate approaches *k*_leak_ = 100 s^−1^ this leakage effect is strong enough to effectively uncouple transport, where antiport behavior is no longer observed (Fig. 11c). In this uncoupled regime the net cycle fluxes for the leakage cycles (A and B) dominate over the coupled cycle flux resulting in both ligands flowing from outside to inside in accordance with their concentration gradients. This decoupling is further demonstrated by the stoichiometry, where as the sodium leakage cycle flux overcomes the coupled cycle flux (i.e., *J*_B_ > *J*_C_) the stoichiometry drops below zero. Overall a shift in transporter function from sodium efflux with ideal stoichiometry to sodium influx with poor transport efficiency was observed.

#### 3.2.2 Rate Bias Effects on EmrE Phenotype

Here we apply the diagram method to a realistic model of a drug-proton antiporter with experimentally measured rates, the so-called Free Exchange Model of EmrE.^36^ EmrE is an ideal candidate for our analysis as the rate constants for all microscopic steps of the transporter cycle are available together with a numerical analysis of the model.^36^

EmrE is a proton-coupled multidrug efflux pump from *Escherichia coli*, ^36,57,58^ broadly categorized as a secondary active transporter. ^59^ EmrE uses the inward-facing proton gradient of *E. coli* to drive drug efflux.^60^ EmrE typically exhibits an antiport phenotype but has been shown to adopt additional states atypical for a pure antiporter.^36,59^ The Free Exchange Model of EmrE incorporates all experimentally observed states including minor and major EmrE conformations, as well as leakage pathways that allow for cotransport and exchange of both ion and substrate. ^36^ The inclusion of all pathways allows us to observe which pathways are preferential (e.g. symport, antiport) under different environmental conditions. ^33^ Henzler-Wildman et al. discuss a minimal 8-state model with a single protonation event and a 10-state model with two protonation events. Here we are focusing on the 8-state model (Fig. 12a) as the more parsimonious model while also serving as a good comparison to the 6-state antiporter model from the previous section.

**Figure 12:**
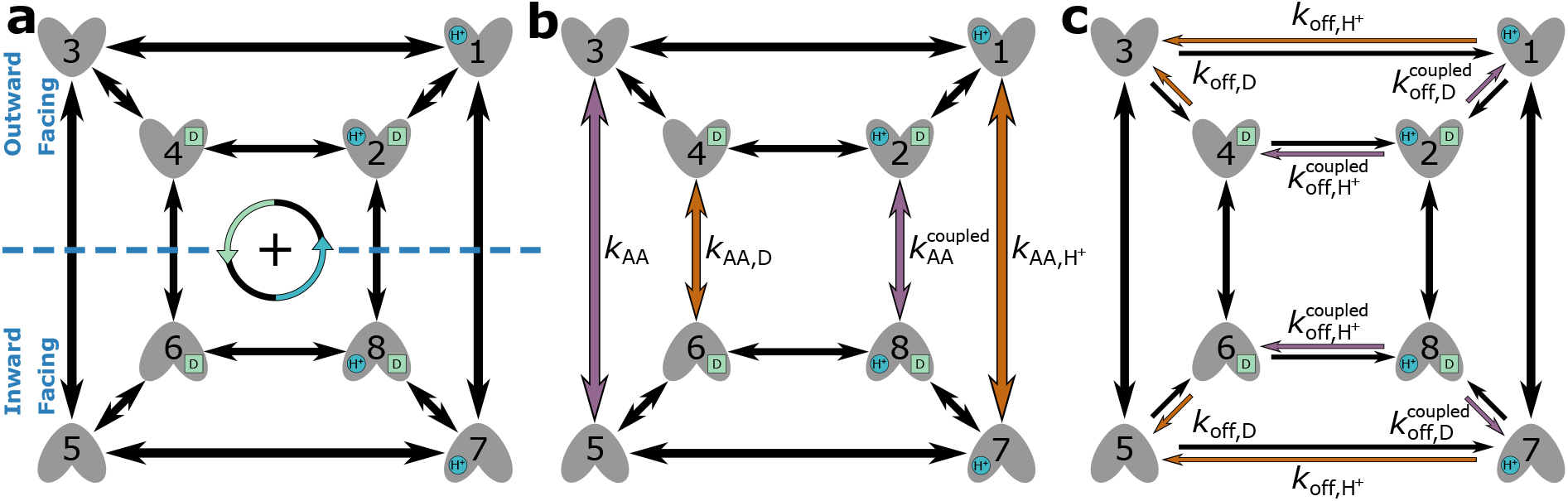
Free Exchange Model of EmrE. (a) Kinetic model of EmrE. Proton and drug molecules are shown in blue and green, respectively, with the positive cycle direction defined as counter-clockwise (CCW). (b) Kinetic model of EmrE with alternating access transitions highlighted. Symport and antiport biased transition arrows are colored purple and orange, respectively. (c) Kinetic model of EmrE with substrate unbinding transitions highlighted. Symport and antiport biased transition arrows are colored purple and orange, respectively.

The original study^36^ modeled EmrE by numerically solving a system of coupled nonlinear ordinary differential equations for a range of different rates. They showed that biasing specific kinetic rates increased probabilities of specific pathways along the kinetic diagram affecting overall transporter function; multiple calculations were ran under different biasing conditions including biased alternating access rates and substrate unbinding (off) rates. Calculations were initialized with an outward-facing proton gradient 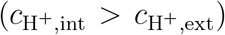 and vanishing drug concentration gradient (*c*_D,int_ = *c*_D,ext_), with a constant pH. The drug gradient was used to determine transporter phenotype by comparing the final (i.e., steady-state) drug concentration against the initial drug concentration. With a constant pH, at steady-state an inward-facing drug gradient (*c*_D,int_ > *c*_D,ext_) indicated the antiport phenotype while an outward-facing drug gradient (*c*_D,int_ < *c*_D,ext_) indicated the symport phenotype.

Instead, in our analysis we directly characterize changes in EmrE phenotype by calculating the operational fluxes for both ion and substrate using system parameters from the original study. ^36^ Since steady-state fluxes are calculated directly, equal and constant internal and external drug concentrations are maintained for all calculations.

To calculate the net cycle fluxes for EmrE first the unique cycles in the kinetic diagram were identified. The 8-state kinetic diagram for EmrE contains 28 unique cycles (Fig. 13) where 16 contribute to proton transport, 16 contribute to drug transport, and 4 contribute to neither. Transport contribution is determined by tracking substrate location throughout the cycle; cycles for which a single cycle completion results in the net transport of one or both substrates across the membrane are considered contributors. For example, cycle 1 has a net transport of a single proton since it transports a proton from outside to inside in a counter-clockwise cycle completion, and thus is a contributing cycle to the proton turnover. Cycles with a net transport of both substrates contribute to both operational flux calculations, while cycles with zero net transport do not contribute to either. The operational fluxes for protons and drug molecules were calculated using Eq. 11,

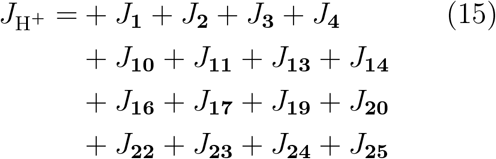

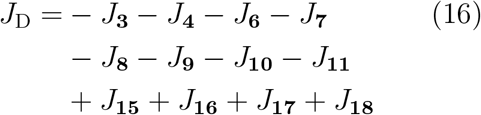

where *J*_*i*_ is the net cycle flux of cycle *i* in Fig 13. Both operational flux expressions have a term for each contributing cycle resulting in 16 terms per expression. Net cycle flux signs are determined according to the cycle direction definition, where the positive cycle direction was defined as counter-clockwise (CCW).

**Figure 13:**
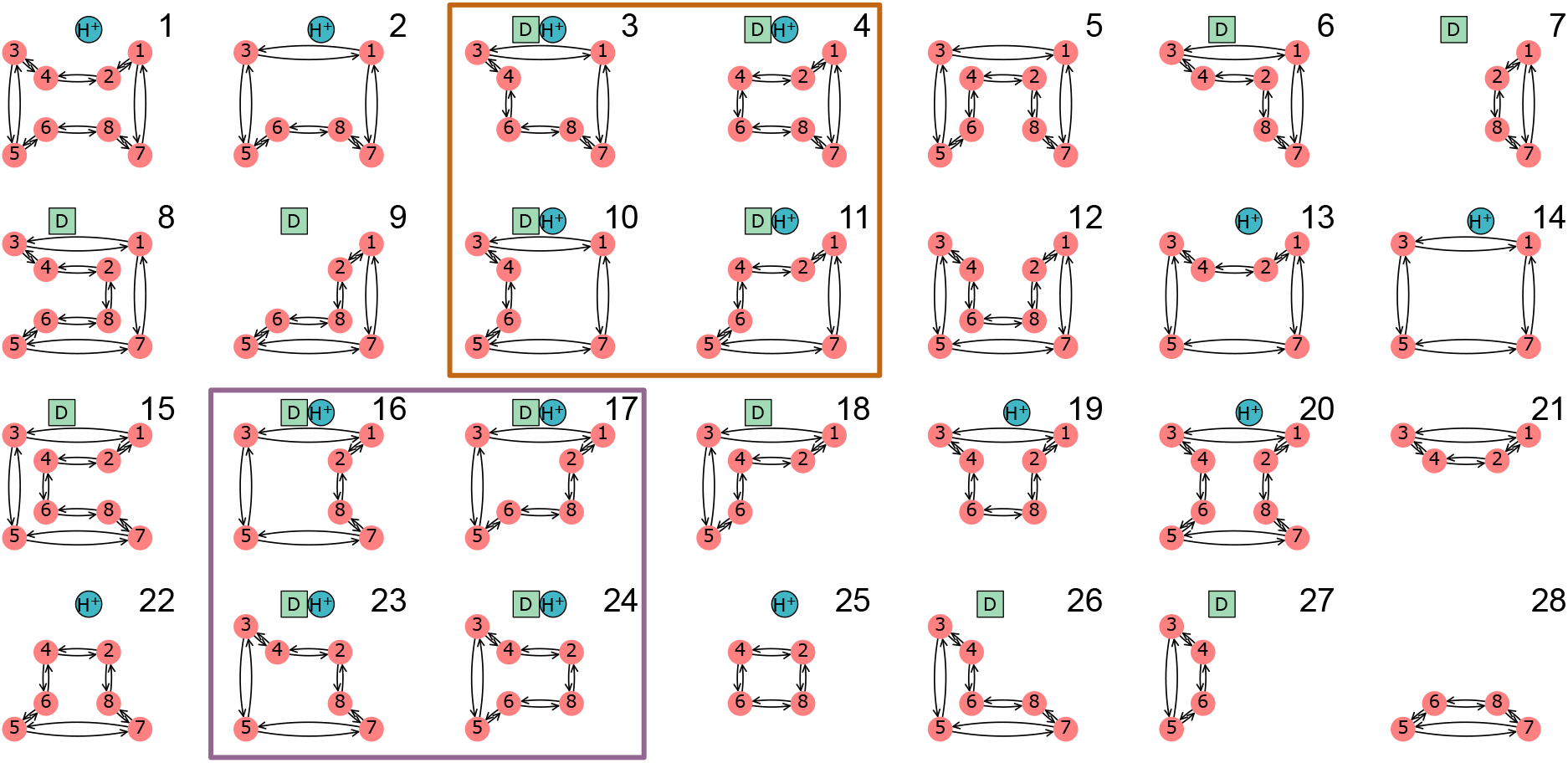
EmrE Free Exchange Model Net Transport by Cycle. All cycles for the 8-state model of EmrE with the net cycle transport shown above each cycle. Cycles which contribute to proton and drug transport are labeled with blue and green polygons, respectively. Symport cycles (boxed in purple) transport both a proton and a drug molecule in the same direction, while antiport cycles (boxed in orange) transport the proton in the opposing direction of the drug molecule. Positive and negative contributions are defined as counter-clockwise cycle completions which transport a substrate from inside to outside or outside to inside, respectively.

Operational flux calculations were performed under different parameter sets taken from the original study. ^36^ Two subsets of system parameters were varied independently to observe phenotype changes: alternating access rates (Fig. 12b) and substrate unbinding (off) rates (Fig. 12c). Alternating access rate biasing influences coupling directly by increasing relative transition probabilities of either antiport or symport pathways. Substrate off-rate biasing influences transport indirectly by changing relative preference for unbinding the doubly bound states versus single bound states on one side of the membrane. The parameter sets for both analyses are included in Table 2.

**Table 2:** Rate Constant Definitions for EmrE Free Exchange Model.

| Parameter | Transition | Process | Value (Fig. 14) | Value (Fig. 15) | Unit |
| --- | --- | --- | --- | --- | --- |
| $k_{\text{on,H}^+}^{\circ}$ | $3 \rightarrow 1$ | Proton-on | $1 \times 10^{10}$ | $1 \times 10^{10}$ | $\text{M}^{-1}\text{s}^{-1}$ |
| | $5 \rightarrow 7$ | Proton-on | $1 \times 10^{10}$ | $1 \times 10^{10}$ | $\text{M}^{-1}\text{s}^{-1}$ |
| | $4 \rightarrow 2$ | Proton-on | $1 \times 10^{10}$ | $1 \times 10^{10}$ | $\text{M}^{-1}\text{s}^{-1}$ |
| | $6 \rightarrow 8$ | Proton-on | $1 \times 10^{10}$ | $1 \times 10^{10}$ | $\text{M}^{-1}\text{s}^{-1}$ |
| $k_{\text{off,H}^+}^{\text{EH}}$ | $3 \leftarrow 1$ | Proton-off | $1 \times 10^3$ | $10^{-2} - 10^8$ † | $\text{s}^{-1}$ |
| | $5 \leftarrow 7$ | Proton-off | $1 \times 10^3$ | $10^{-2} - 10^8$ † | $\text{s}^{-1}$ |
| $k_{\text{off,H}^+}^{\text{EHD}}$ | $4 \leftarrow 2$ | Proton-off | $1 \times 10^3$ | $10^{-2} - 10^8$ † | $\text{s}^{-1}$ |
| | $6 \leftarrow 8$ | Proton-off | $1 \times 10^3$ | $10^{-2} - 10^8$ † | $\text{s}^{-1}$ |
| $k_{\text{on,D}}^{\circ}$ | $3 \rightarrow 4$ | Drug-on | $1 \times 10^7$ | $1 \times 10^7$ | $\text{M}^{-1}\text{s}^{-1}$ |
| | $5 \rightarrow 6$ | Drug-on | $1 \times 10^7$ | $1 \times 10^7$ | $\text{M}^{-1}\text{s}^{-1}$ |
| | $1 \rightarrow 2$ | Drug-on | $1 \times 10^7$ | $1 \times 10^7$ | $\text{M}^{-1}\text{s}^{-1}$ |
| | $7 \rightarrow 8$ | Drug-on | $1 \times 10^7$ | $1 \times 10^7$ | $\text{M}^{-1}\text{s}^{-1}$ |
| $k_{\text{off,D}}^{\text{ED}}$ | $3 \leftarrow 4$ | Drug-off | 1 | $10^{-4} - 10^6$ † | $\text{s}^{-1}$ |
| | $5 \leftarrow 6$ | Drug-off | 1 | $10^{-4} - 10^6$ † | $\text{s}^{-1}$ |
| $k_{\text{off,D}}^{\text{EHD}}$ | $1 \leftarrow 2$ | Drug-off | 1 | $10^{-4} - 10^6$ † | $\text{s}^{-1}$ |
| | $7 \leftarrow 8$ | Drug-off | 1 | $10^{-4} - 10^6$ † | $\text{s}^{-1}$ |
| $k_{\text{AA}}^{\text{EH}}$ | $7 \rightarrow 1$ | Alternating access | $10^{-3} - 10^5$ † | 1, 10, 100, 1000 | $\text{s}^{-1}$ |
| | $7 \leftarrow 1$ | Alternating access | $10^{-3} - 10^5$ † | 1, 10, 100, 1000 | $\text{s}^{-1}$ |
| $k_{\text{AA}}^{\text{ED}}$ | $6 \rightarrow 4$ | Alternating access | $10^{-3} - 10^5$ † | 1, 10, 100, 1000 | $\text{s}^{-1}$ |
| | $6 \leftarrow 4$ | Alternating access | $10^{-3} - 10^5$ † | 1, 10, 100, 1000 | $\text{s}^{-1}$ |
| $k_{\text{AA}}^{\text{E}}$ | $5 \rightarrow 3$ | Alternating access | $10^{-3} - 10^5$ † | 1, 10, 100, 1000 | $\text{s}^{-1}$ |
| | $5 \leftarrow 3$ | Alternating access | $10^{-3} - 10^5$ † | 1, 10, 100, 1000 | $\text{s}^{-1}$ |
| $k_{\text{AA}}^{\text{EHD}}$ | $8 \rightarrow 2$ | Alternating access | $10^{-3} - 10^5$ † | 1, 10, 100, 1000 | $\text{s}^{-1}$ |
| | $8 \leftarrow 2$ | Alternating access | $10^{-3} - 10^5$ † | 1, 10, 100, 1000 | $\text{s}^{-1}$ |
| $\text{pH}_{\text{ext}}$ | $3 \rightarrow 1$ | Proton-on | 7.5 | 7.5 | |
| | $4 \rightarrow 2$ | Proton-on | 7.5 | 7.5 | |
| $\text{pH}_{\text{int}}$ | $5 \rightarrow 7$ | Proton-on | 6.5 | 6.5 | |
| | $6 \rightarrow 8$ | Proton-on | 6.5 | 6.5 | |
| $c_{\text{D,ext}}$ | $3 \rightarrow 4$ | Drug-on | $25 \times 10^{-9}$ | $25 \times 10^{-9}$ | M |
| | $1 \rightarrow 2$ | Drug-on | $25 \times 10^{-9}$ | $25 \times 10^{-9}$ | M |
| $c_{\text{D,int}}$ | $5 \rightarrow 6$ | Drug-on | $25 \times 10^{-9}$ | $25 \times 10^{-9}$ | M |
| | $7 \rightarrow 8$ | Drug-on | $25 \times 10^{-9}$ | $25 \times 10^{-9}$ | M |

† Rates are varied in accordance with the thermodynamic consistency condition, Eq. 5.

##### 3.2.2.1 Rate Biasing via Alternating Access Rates

We first investigate if changing the conformational change rates (Fig. 12b) is sufficient to switch EmrE phenotype from symport to antiport. Transporter phenotype was determined based on the directions of operational fluxes, where same-sign fluxes (cotransport) indicate the symport phenotype and opposite-sign fluxes (exchange) indicate the antiport phenotype. Operational fluxes (i.e., Eqs. 15-16) were calculated as a function of *R*_AA_,

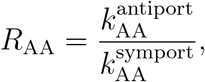

where *R*_AA_ > 1 favors antiport, *R*_AA_ < 1 favors symport, and

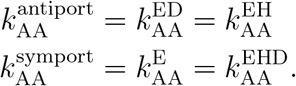

Calculations were performed with fixed substrate concentration gradients (Table 2, column 4), where the alternating access rates (e.g.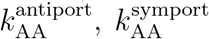) were varied across the range 10^−3^ − 10^5^ s^−1^ in accordance with the thermodynamic consistency condition Eq. 5.

Starting with the unbiased regime (*R*_AA_ = 1), peak proton flux of 2.6 s^−1^ is reached with zero drug flux, resulting in a stoichiometry (i.e., drug transported per proton) of zero (Fig. 14a). As the alternating access rates are biased towards symport (i.e., *R*_AA_ < 1), the drug flux increases to 0.1 s^−1^ and the proton flux decreases to− 0.1 s^−1^. Biasing towards antiport (i.e., *R*_AA_ > 1) results in the drug flux approaching 0.1 s^−1^ while the proton flux decreases to 0.1 s^−1^. For both symport and antiport-favored regimes, the substrate fluxes result in an approach to ideal 1:1 stoichiometry.

**Figure 14:**
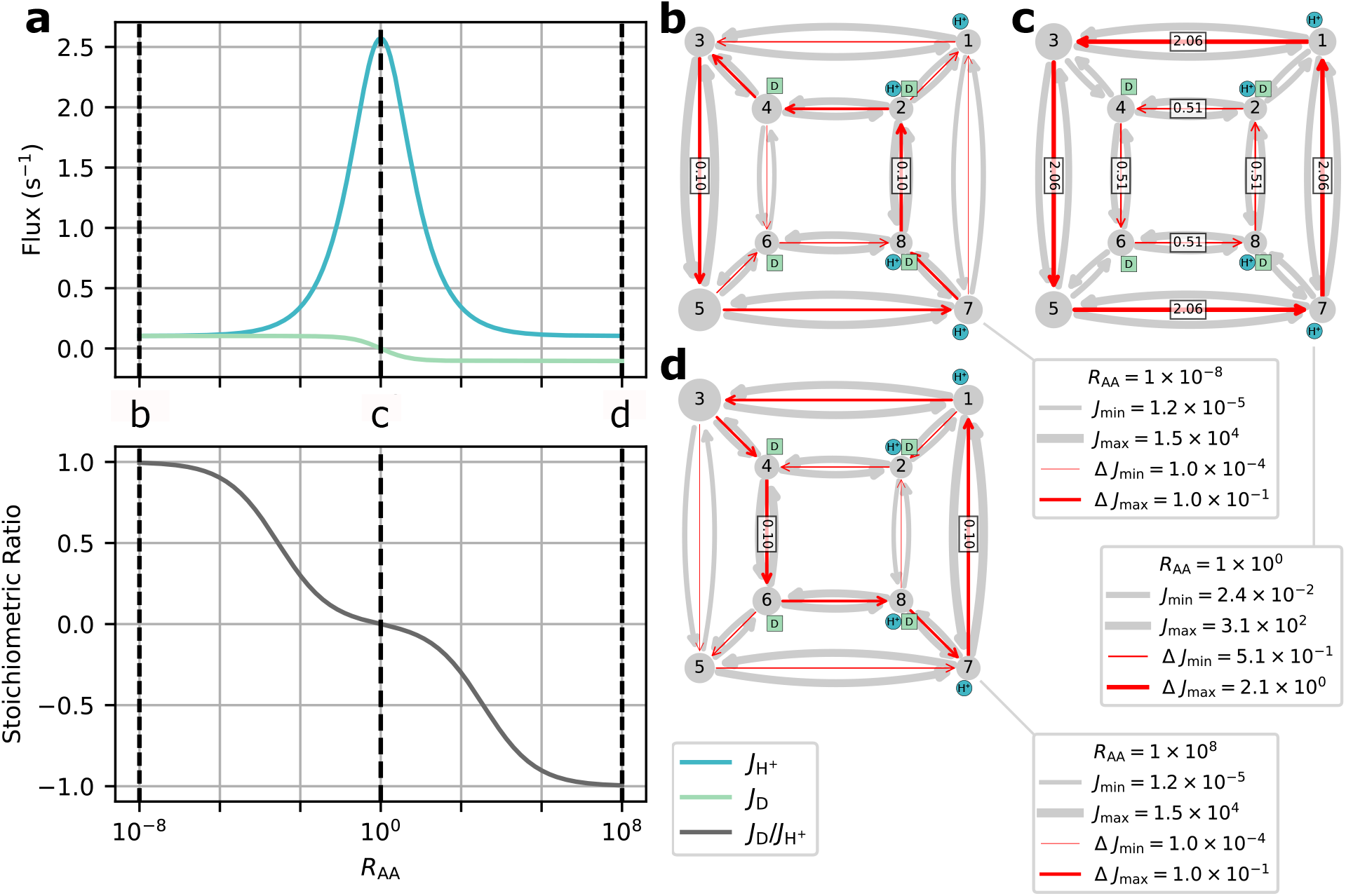
Alternating Access Rate Biasing Affects EmrE Phenotype. (a) Operational fluxes and stoichiometric ratio for protons and drug molecules as a function of the alternating access biasing parameter *R*_AA_. Operational fluxes are calculated from a symport-biased regime (*R*_AA_ < 1) to an antiport-biased regime (*R*_AA_ > 1). (b-d) Kinetic diagrams of EmrE with the transition fluxes (gray arrows) and net transition fluxes (red arrows) overlaid. The cases shown (in order) are for ideal symport (*R*_AA_≪ 1), unbiased/uncoupled (*R*_AA_ = 1), and ideal antiport (*R*_AA_≫ 1), respectively, with text boxes displayed for net transition fluxes of magnitude *J*_*ij*_ > 0.02 s^−1^. Nodes and arrows are min-max scaled according to Eqs. S3 and S4 from the Supporting Information, respectively.

In the different biasing regimes specific pathways along the kinetic diagram contributed much more to the overall operational fluxes than others (see Table S1 of the Supporting Information). Because cycles are specific collections of transitions that each represent a change of state of the system, a stronger cycle contribution indicates a given sequence of transitions (i.e., a *molecular mechanism*) is more likely than another. For example, in the symport-favored regime cycles 16, 17, 23, and 24 (Fig. 13) all contributed strongly with cycle 23 as the primary contributor (Fig. 14b). Cycle 23 points to the importance of internal proton binding and external proton release under symport-biased conditions. In the unbiased regime, cycles 25 and especially 14 were the primary contributors to the operational flux (Fig. 14c). In the antiport-biased regime, cycle 3 was the largest contributor (Fig. 14d) together with 4, 10 and 11, demonstrating a preference for external proton release and internal proton binding in the antiport regime.

Our findings are consistent with the original study, ^36^ where biasing alternating access rates results in a change in transport phenotype from symport to antiport based on the biasing direction. Both symport and antiport phenotypes exhibited a peak drug turnover of 0.1 s^−1^. While the observed drug turnover is relatively slow for a transporter, work by Yerushalmi et al.^58^ reported a comparable value for EmrE turnover of 14 min^−1^ (i.e., 0.23 s^−1^) under typical conditions.

##### 3.2.2.2 Rate Biasing via Substrate Off-rates

Instead of biasing conformational change rates of the model to affect transporter phenotype, here we investigate if similar phenotypical changes can be achieved by indirect manipulation of the substrate unbinding rates (as proposed in Hussey et al^36^). These rates depend on substrate concentrations and thus are more easy to manipulate experimentally or in a physiological context. Following the prior analysis, transporter phenotype was determined from the directions of operational fluxes (i.e., same-sign fluxes indicate symport and opposite-sign fluxes indicate antiport). Operational fluxes (Eqs. 15-16) were calculated as a function of *R*_off_,

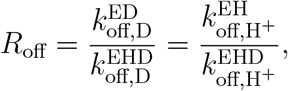

where *R*_off_ < 1 favors antiport and *R*_off_ > 1 favors symport. Calculations were performed with fixed substrate concentration gradients (Table 2, column 5), where substrate off-rates(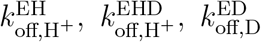, and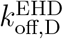, Table 2) for protons and drugs were kept uniform (e.g.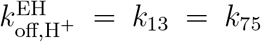) and varied from 10^−2^ − 10^8^ s^−1^ and 10^−4^− 10^6^ s^−1^, respectively. For all calculations the alternating access rate was varied uniformly,

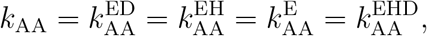

where the rate was set to one of four distinct values, 1 s^−1^, 10 s^−1^, 100 s^−1^, or 1000 s^−1^. All rates (i.e., substrate unbinding, alternating access) were varied in accordance with the thermodynamic consistency condition Eq. 5.

In the unbiased case (*R*_off_ = 1), the proton gradient drives a positive proton flux that increases with increasing alternating access rate *k*_AA_ but in the absence of coupling, the net drug flux remains zero and hence the stoichiometric ratio is also zero (Fig. 15a). With substrate off-rates biased towards antiport (*R*_off_ < 1), the positive proton flux decreases relative to *R*_off_ = 1 but drives a drug flux in the negative direction (Fig. 15a), eventually resulting in 1:1 stoichiometry for *R*_off_ < 10^−8^. Both substrate fluxes increase in magnitude with increasing alternating access rate. Biasing towards symport (*R*_off_ > 1) also decreases the proton flux although the drug flux exhibits a clear maximum around *R*_off_ *≈* 10^2^ and vanishes for *R*_off_ ≫ 1 (Fig. 15a), in contrast to the antiport regime (*R*_off_ < 1). Furthermore, increasing alternating access rates for *R*_off_ > 1 does not result in a monotonic increase in the peak drug flux. Instead the largest peak drug fluxes are observed for *k*_AA_ = 10 s^−1^ and *k*_AA_ = 100 s^−1^, while the *k*_AA_ = 1 s^−1^ and *k*_AA_ = 1000 s^−1^ cases result in lower peak drug fluxes.

**Figure 15:**
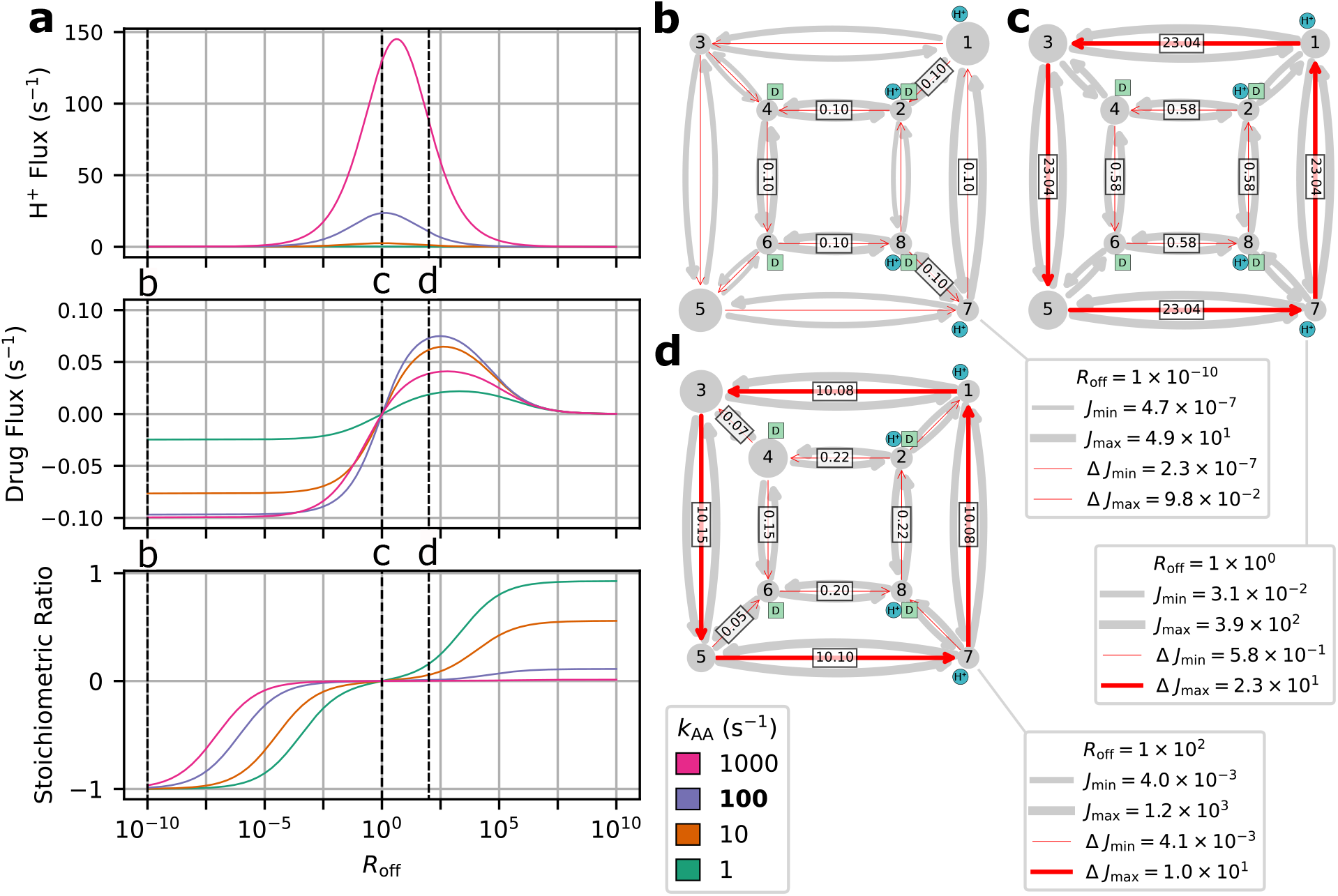
Substrate Unbinding Pathway Biasing Affects EmrE Function. (a) Operational fluxes and stoichiometric ratio for protons and drug molecules as a function of the biasing parameter *R*_off_ and the alternating access rate *k*_AA_. Operational fluxes are calculated from an antiport regime, *R*_off_ < 1, to a symport regime, *R*_off_ > 1. (b-d) Kinetic diagrams of EmrE with the transition fluxes (gray arrows) and net transition fluxes (red arrows) overlaid. The cases shown (in order) are for ideal antiport (*R*_off_ ≪ 1), unbiased/uncoupled (*R*_off_ = 1), and maximum symport drug flux (*R*_off_ = 100), respectively, with text boxes displayed for net transition fluxes of magnitude *J*_*ij*_ > 0.02 s^−1^. All fluxes are calculated for the *k*_AA_ = 100 s^−1^ case. Nodes and arrows are min-max scaled according to Eqs. S3 and S4 from the Supporting Information, respectively.

For the unbiased and antiport-favored regimes (*R*_off_ ≤ 1), a subset of cycles account for nearly all the operational flux. We will focus only on the *k*_AA_ = 100 s^−1^ case for simplicity. In the unbiased case, the proton leakage cycles 14 and 25 (Fig. 13; see also Table S2 of the Supporting Information) were the primary contributors to the operational flux (Fig. 15c).

With strong antiport biasing, proton leakage remains present (cycle 14) but the coupled antiport cycle 4 is now the dominant mechanism (Fig. 15b). With mild symport biasing (i.e., *R*_off_ = 100), a mixture of proton leakage cycles (14, 25) and coupled symport cycles (24 and 23) contribute to both operational fluxes (Fig. 15d).

Our findings are consistent with the original study, ^36^ where the indirect biasing of substrate unbinding rates results in a change in transport phenotype from antiport to symport. While the overall transport behavior was observed, antiport and symport phenotypes had notable differences in behavior and efficiency. The antiport regime exhibits a saturation behavior, where peak drug flux is reached and maintained for a range of *R*_off_ values. The symport-favored regime exhibits a different behavior, where mild off-rate biasing results in an initial drug flux peak, and further off-rate biasing results in an approach to zero drug flux. In terms of efficiency, strong antiport biasing results in peak drug flux (0.10 s^−1^) while approaching ideal 1:1 stoichiometry. Thus, in the highly antiport-biased regime there is efficient usage of protons to drive drug transport. In the symport regime, the peak drug flux (0.07 s^−1^) is relatively smaller and is only observed with poor stoichiometry (< 0.2 drugs/proton). Thus, for the symport-favored regime, ideal stoichiometry is only achieved under strong symport biasing where the drug fluxes approach zero. This suggests that while both antiport and symport regimes ultimately resulted in coupled drug transport, the symport biasing requires more fine tuning of system parameters to achieve efficient drug transport.

In the antiport-favored regime single cycles were observed to be preferential while the symport regime exhibited a mixture of cycles with relatively similar fluxes. For the peak antiport drug flux case (*R*_off_ = 10^−10^) the substrate fluxes were driven primarily by single cycles. Under symport-favored biasing, the peak drug flux occurs near *R*_off_ = 100 for the *k*_AA_ = 100 s^−1^ case. In this narrow symport regime, the substrate fluxes were driven by several cycles each. dominant overall. The reason for the lower efficiency of symport in this model is that the symport cycles compete with uniport cycles, in particular, proton leakage cycles. With increasing alternating access rate, leakage increases together with the coupled symport pathways and leading to overall inefficient transport as suggested by the original study. ^36^ The cycle analysis clearly demonstrates the sensitivity of symport pathways to leakage when the alternating access rates are a similar order of magnitude as the substrate off-rates (for *R*_off_ = 100 the drug off-rates are 10^1^− 10^3^ s^−1^), resulting in less efficient cotransport and smaller drug fluxes.

### 3.3 Operational Fluxes in Terms of Net Transition Fluxes

The diagrammatic method provides a clear prescription for how to compute operational fluxes from net cycle fluxes. The operational fluxes ultimately quantify the system’s *function* and play the role of observables while the net cycle fluxes are not observable but represent the fundamental theoretical quantities to the diagrammatic method. In the previous sections we used `KDA` to analyze different transporter models using operational fluxes calculated from net cycle fluxes. Here we discuss an alternative approach that employs net transition fluxes to create the operational flux expressions, as previously discussed in Section 2.7.3. The primary advantage of using transition fluxes is that they are generally easier to obtain because only the rates and the state probabilities, which are straightforward to calculate numerically, are required.

Unlike for the derivation of operational fluxes from net cycle fluxes, a prescription for forming operational fluxes from net transition fluxes is less well documented. Hill directly writes down the appropriate expressions as “physically obvious” although he also shows that, in general, the operational/transition flux relationships can be derived from the master equations evaluated at steady-state. ^15^ George et al.^5^ derive the relevant operational fluxes based on their ansatz

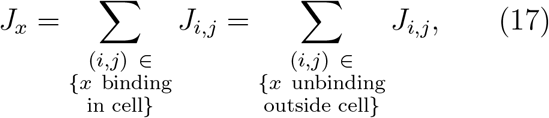

where *J*_*x*_ is the net transport for a ligand *x* and the flux is computed from the sum of all net transition fluxes that either lead to binding or unbinding of the ligand. Eq. 17 intuitively captures the relevant fluxes and we ultimately use the same ansatz to obtain operational flux expressions but we will show how we can use `KDA` symbolic expressions to prove the equivalence between the net cycle flux and net transition flux expressions for operational fluxes for cases when this is difficult to do manually.

All the diagram method fluxes (transition, cycle, and operational) can be expressed in terms of one another. For simple models like the 6-state antiporter model (Fig. 11b, model G) the relationships between all levels of fluxes can be written down directly. The 6-state model contains only one cycle (cycle C, Fig. 11b) and therefore one net cycle flux. Thus, the relationships to the operational fluxes are straightforward, 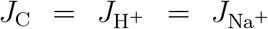Additionally, since net transition fluxes are sums of net cycle fluxes, all counter-clockwise net transition fluxes along cycle C are equal to the net cycle flux, *J*_C_ = *J*_6,5_ = *J*_5,4_ = … = *J*_1,6_.

In general, the operational/transition flux relationships are derived from the master equations evaluated at steady-state^15^ but for more complex models these relationships are not necessarily obvious. However, some of the flux relationships may be intuited from the kinetic model directly (Eq. 17) and then proven to be equivalent to the net cycle expressions. For example, for the leakage variant of the 6-state antiporter model (Fig. 11b, model G_leak_) we guess that the operational fluxes are expressed in terms of single net transition fluxes involving the binding/unbinding of the proton (*J*_1,2_, *J*_3,4_) or sodium ion (*J*_6,1_, *J*_4,5_) or their transport across the membrane (*J*_2,3_, *J*_5,6_)

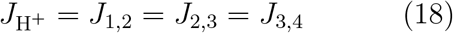

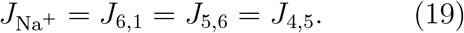

We prove Eqs. 18–19 by finding the net transition fluxes in terms of net cycle fluxes (using the relationship from Section 2.5, *J*_*i,j*_ is equal to the sum of net cycle fluxes whose cycle traverses the transition *i* → *j*),

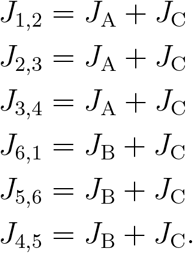

then comparing to the net cycle flux sums from the original operational flux relationships (Eqs. 13–14):

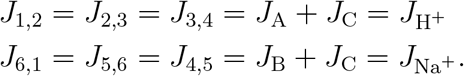

For more complicated models like the 8-state model of EmrE (Fig. 12) we first inspect the binding/unbinding transitions to guess the operational flux relationships. For the EmrE model there are two extracellular binding/unbinding reactions for each ligand: transition pairs 1 ↔ 3 and 2 ↔ 4 unbind protons externally while transition pairs 2 ↔ 1 and 4 ↔ 3 bind external drugs. Summing the net transition fluxes for these transitions yields the operational fluxes

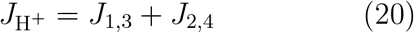

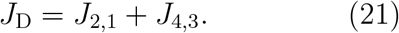

as sums of multiple transition fluxes, where each net transition flux contributes additively to the operational flux. A similar analysis of the intracellular reactions of EmrE would yield different but equivalent operational flux expressions. These flux relationships need to be shown to be equivalent to the ones derived from net cycle fluxes. In Section 5 of the Supporting Information we explicitly prove Eqs. 20–21 by expressing the net transition fluxes in terms of net cycle fluxes and calculating the appropriate sums.

However, the manual calculations are tedious and error-prone, even for models of moderate complexity such as the EmrE 8-state one, so here we show that we can prove the equivalence of cycle and transition flux expressions for the operational fluxes using the `SymPy` computer algebra system^17^ with the symbolic expressions generated by `KDA`. The operational flux expressions for the 6-state antiporter model (Fig. 11a, model G_leak_) and the 8-state model of EmrE (Fig. 12) were generated with `KDA` (see Section 8 of the Supporting Information for the full code for each model). `SymPy` proved the expressions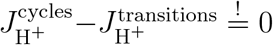 and 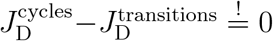for both the 6-state antiporter model and 8-state model of EmrE as shown in Fig. 16.

**Figure 16:**
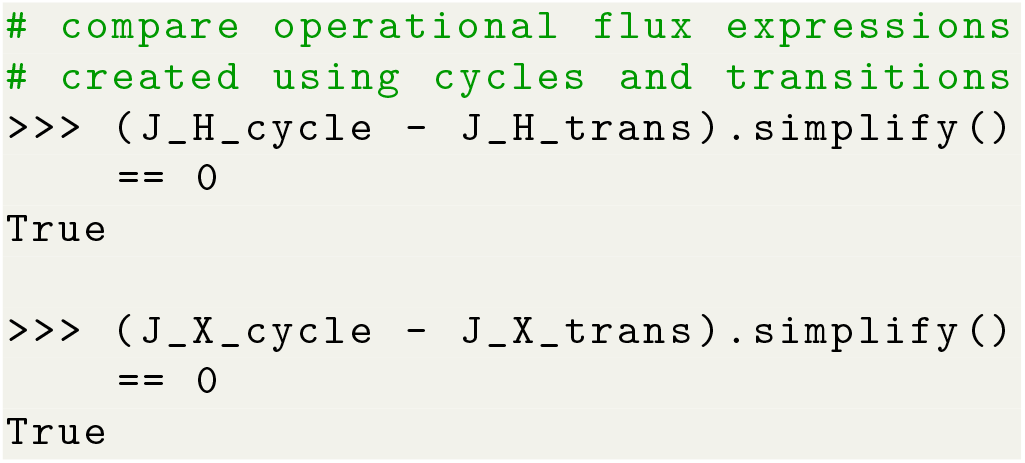
Proof of equivalence of operational flux expressions using computer algebra with `SymPy`. Python code which compares the `KDA`-generated operational flux expressions (created using cycles or transitions). The full code for the 6-state antiporter model (Fig. 11b, model G_leak_) and the 8-state model of EmrE (Fig. 12) are included in Section 8 of the Supporting Information.

Finally, as an alternative demonstration of the equivalence of the transition and cycle flux calculations we also include a numerical comparison; the advantage of the numerical approach is that it can be easily implemented as a validation check in the absence of symbolic expressions and it provides a sense of numerical sensitivity. For the 6-state model, the numerical results were calculated for several values of the leakage rate (*k*_leak_), along with the difference between each operational flux and its corresponding net transition flux (from Eqs. 18-19). For EmrE, the fluxes (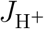 and *J*_D_) were calculated for both rate biasing cases (alternating access and substrate off-rate biasing) using the net cycle flux expressions (i.e., Eqs. 15-16) from the original analysis. For the 6state model the observed operational flux differences were exactly zero for both substrate species, demonstrating agreement between the two operational flux methods (see Table S3 in the Supporting Information). For EmrE, the typical difference observed is close to machine floating point precision, and thus computationally indistinguishable from zero (see Table S4 in the Supporting Information). For several cases the EmrE flux differences are relatively larger. Since the expressions are algebraically equivalent, these numerical disagreements are likely attributable to floating point arithmetic error that accumulates when mixing Python doubles with `SymPy`^17^ expressions.

## 4 Conclusion

The diagram method developed by King and Altman^12^ and Hill^13,14^ yields exact rational algebraic expressions to directly calculate functionally relevant steady-state fluxes observables of any kinetic model. Our `Kinetic_Diagram_Analysis` (`KDA`) Python package programmatically generates the necessary graphs and expressions from a user-defined kinetic diagram, a task that is otherwise infeasible for diagrams of even modest complexity due to the combinatorially large number of intermediate diagrams required. Our Python implementation is designed to be modular so that it can be easily integrated into existing workflows or used interactively in Jupyter notebooks; overall, we provide an accessible and interoperable solution to generate symbolic solutions of the diagram method.

To demonstrate the capability of the diagram method and validate `KDA` outputs we applied our methods to models for secondary active transporter proteins of increasing complexity. With a 6-state antiporter model (similar to the model studied by Berlaga and Kolomeisky ^37^) we demonstrated how the addition of a leakage transition to a generic antiporter model decreases transport efficiency (i.e., number of driving sodium ions transported per transported proton) while substrate and driving ion fluxes increase in the direction of their concentration gradients. Using the 8-state free exchange model of EmrE^36^ we confirmed the major conclusions of the original study, namely the ability to switch the model between antiporter and symporter phenotype by changes to specific rates, by directly computing physiologically relevant transport turnover numbers under different conditions. We further demonstrated how the experimentally observable operational fluxes can be either calculated from cycle fluxes in a general manner within the framework of the diagram method or as sums of judiciously chosen transition fluxes, which can be computed outside the diagram method. In this case, `KDA` can be used to rigorously prove the equivalence of a cycle flux and a transition flux formulation of an operation flux via symbolic computer algebra with `SymPy`.^17^

`KDA` is still early in development and thus has many avenues for improvement. For example, the “test and select” spanning tree algorithm used for partial diagram generation could be replaced with a more computationally efficient version that does not require the generation of all edge combinations, or one of the many other spanning tree algorithms, ^40^ thus improving performance for larger graphs. Although it is unlikely that `KDA` will ever achieve the speed of a pure numerical solution, it can guarantee exact solutions that are amenable to symbolic manipulation. Future work in this direction may explore directly taking derivatives to calculate the effect of changes in one or more variables on, e.g., operational fluxes, or introducing new external driving forces by replacing a rate law with an alternative functional form, e.g., to make rates dependent on mechanical external forces^61^ or membrane voltages.^62^

The fundamental assumptions underlying `KDA` are that all processes are reversible (have forward and backward rates) and all rates are thermodynamically consistent.^38^ Under these conditions, `KDA` will generate exact algebraic expressions under steady-state conditions. The diagram method is not limited to biochemical systems, and while our analysis focuses on transporter models of increasing complexity, other kinetic networks can be analyzed using the same methods.

We demonstrated that `KDA` can provide useful results for typical transporter models from the literature. Perhaps `KDA`’s most appealing feature is its ability to generate exact symbolic expressions that can be further analyzed and manipulated with a computer algebra system or used to efficiently generate exact numerical results for different external parameters. Thus, `KDA` is ideally suited for computing reference values to validate other approximate schemes and to explore theoretical models for non-equilibrium processes.

## Acknowledgement

The authors thank Chenou Zhang for his help with finding experimental rates for a sodium proton antiporter, and Ian Kenney for his assistance with recursive programming, ion diffusion theory, and comments on the manuscript. Research reported in this work was supported by the National Institute Of General Medical Sciences of the National Institutes of Health under Award No R01GM118772. The funders had no influence on the research or the writing of the manuscript.

## Supporting Information Available

The Supporting Information is available free of charge at https://.

4-state leakage model master equation derivation; 4-state leakage model unnormalized state probability expressions; Comparison of two models to observe effects of dangling nodes on flux magnitudes; Min-max normalization for plotting steady-state observables over kinetic diagrams; `KDA` vs. `KAPattern` expression disagreement; EmrE flux relationships; EmrE net cycle flux numerical data; 6-state antiporter model and 8-state model of EmrE operational flux comparison numerical data; Operational flux expression comparison code for the 6-state antiporter model and 8-state model of EmrE (PDF)

## References

(1) Mescam, M.; Vinnakota, K. C.; Beard, D. A. Identification of the catalytic mechanism and estimation of kinetic parameters for fumarase. The Journal of Biological Chemistry 2011, 286, 21100–21109.

(2) Iwahara, J.; Kolomeisky, A. B. Discrete-state stochastic kinetic models for target DNA search by proteins: Theory and experimental applications. Biophysical Chemistry 2021, 269, 106521.

(3) Reinhardt, C. R.; Konstantinovsky, D.; Soudackov, A. V.; Hammes-Schiffer, S. Kinetic model for reversible radical transfer in ribonucleotide reductase. Proceedings of the National Academy of Sciences 2022, 119, e2202022119.

(4) Bisignano, P.; Lee, M. A.; George, A.; Zuckerman, D. M.; Grabe, M.; Rosenberg, J. M. A kinetic mechanism for enhanced selectivity of membrane transport. PLOS Computational Biology 2020, 16, e1007789.

(5) George, A.; Bisignano, P.; Rosenberg, J. M.; Grabe, M.; Zuckerman, D. M. A systems-biology approach to molecular machines: Exploration of alternative transporter mechanisms. PLOS Computational Biology 2020, 16, e1007884.

(6) Kinz-Thompson, C. D.; Lopez-Redondo, M. L.; Mulligan, C.; Sauer, D. B.; Marden, J. J.; Song, J.; Tajkhorshid, E.; Hunt, J. F.; Stokes, D. L.; Mindell, J. A.; Wang, D. N.; Gonzalez, R. L. Elevator mechanism dynamics in a sodium-coupled dicarboxylate transporter. 2022; https://www.biorxiv.org/content/10.1101/2022.05.01.490196v2.

(7) Gunawardena, J. A linear framework for time-scale separation in nonlinear biochemical systems. PloS One 2012, 7, e36321.

(8) Segel, I. H. Enzyme kinetics: behavior and analysis of rapid equilibrium and steady state enzyme systems; New York : Wiley, 1993.

(9) Cornish-Bowden, A. Fundamentals of Enzyme Kinetics, 4th ed.; John Wiley & Sons, 2013.

(10) Johnson, K. A. Chapter 23 Fitting Enzyme Kinetic Data with KinTek Global Kinetic Explorer. In Methods in Enzymology ; Johnson, M. L., Brand, L., Eds.; Computer Methods Part B; Academic Press, 2009; Vol. 467; pp 601–626.

(11) Bevc, S.; Konc, J.; Stojan, J.; Hodošček, M.; Penca, M.; Praprotnik, M.; Janežič, D. ENZO: A Web Tool for Derivation and Evaluation of Kinetic Models of Enzyme Catalyzed Reactions. PLOS ONE 2011, 6, e22265, Publisher: Public Library of Science.

(12) King, E. L.; Altman, C. A Schematic Method of Deriving the Rate Laws for Enzyme-Catalyzed Reactions. The Journal of Physical Chemistry 1956, 60, 1375–1378.

(13) Hill, T. L. Studies in irreversible thermodynamics IV. Diagrammatic representation of steady state fluxes for unimolecular systems. Journal of Theoretical Biology 1966, 10, 442–459.

(14) Hill, T. L. Free Energy Transduction in Biology ; Elsevier, 1977.

(15) Hill, T. L. Free Energy Transduction and Biochemical Cycle Kinetics; Springer-Verlag: New York, 1989.

(16) Hagberg, A. A.; Schult, D. A.; Swart, P. J. Exploring Network Structure, Dynamics, and Function using NetworkX. Proceedings of the 7th Python in Science Conference. Pasadena, CA USA, 2008; pp 11–15.

(17) Meurer, A. et al. SymPy: symbolic computing in Python. PeerJ Computer Science 2017, 3, e103.

(18) Pring, M. The simulation and analysis by digital computer of biochemical systems in terms of kinetic models. 3. Generator programming. Journal of Theoretical Biology 1967, 17, 436–440.

(19) Rhoads, D. G.; Pring, M. The simulation and analysis by digital computer of biochemical systems in terms of kinetic models. IV. Automatic derivation of enzymic rate laws. Journal of Theoretical Biology 1968, 20, 297–313.

(20) Lam, C. F.; Priest, D. G. Enzyme kinetics. Systematic generation of valid King-Altman patterns. Biophysical Journal 1972, 12, 248–256.

(21) Cornish-Bowden, A. An automatic method for deriving steady-state rate equations. Biochemical Journal 1977, 165, 55–59.

(22) Kinderlerer, J.; Ainsworth, S. A computer program to derive the rate equations of enzyme catalysed reactions with up to ten enzyme-containing intermediates in the reaction mechanism. International Journal of Bio-Medical Computing 1976, 7, 1–20.

(23) Straathof, A. J. J.; Heijnen, J. J. Derivation of Enzymatic Rate Equations Using Symbolic Software. Biocatalysis and Biotransformation 1997,

(24) Fromm, S. J.; Fromm, H. J. A Two-Step Computer-Assisted Method for Deriving Steady-State Rate Equations. Biochemical and Biophysical Research Communications 1999, 265, 448–452.

(25) Varon, R.; Garcia-Sevilla, F.; Garcia-Moreno, M.; Garcia-Canovas, F.; Peyro, R.; Duggleby, R. G. Computer program for the equations describing the steady state of enzyme reactions. Computer applications in the biosciences: CABIOS 1997, 13, 159–167.

(26) Yago, J. M.; García Sevilla, F.; Garrido del Solo, C.; Duggleby, R. G.; Varón, R. A Windows program for the derivation of steady-state equations in enzyme systems. Applied Mathematics and Computation 2006, 181, 837–852.

(27) Qi, F.; Dash, R. K.; Han, Y.; Beard, D. A. Generating rate equations for complex enzyme systems by a computer-assisted systematic method. BMC Bioinformatics 2009, 10, 238.

(28) Loriaux, P. M.; Tesler, G.; Hoffmann, A. Characterizing the Relationship between Steady State and Response Using Analytical Expressions for the Steady States of Mass Action Models. PLoS Computational Biology 2013, 9, e1002901.

(29) Nam, K.-M.; Martinez-Corral, R.; Gunawardena, J. The linear framework: using graph theory to reveal the algebra and thermodynamics of biomolecular systems. Interface Focus 2022, 12, 20220013.

(30) Seshu, S.; Reed, M. B. Linear graphs and electrical networks; Addison-Wesley, 1961.

(31) Henderson, R. K.; Fendler, K.; Poolman, B. Coupling efficiency of secondary active transporters. Current Opinion in Biotechnology 2019, 58, 62–71.

(32) Drew, D.; North, R. A.; Nagarathinam, K.; Tanabe, M. Structures and General Transport Mechanisms by the Major Facilitator Superfamily (MFS). Chemical Reviews 2021, 121, 5289–5335.

(33) Beckstein, O.; Naughton, F. General principles of secondary active transporter function. Biophysics Reviews 2022, 3, 011307.

(34) Mayes, H. B.; Lee, S.; White, A. D.; Voth, G. A.; Swanson, J. M. J. Multi-scale Kinetic Modeling Reveals an Ensemble of Cl–/H+ Exchange Pathways in ClC-ec1 Antiporter. Journal of the American Chemical Society 2018, 140, 1793–1804.

(35) Burtscher, V.; Schicker, K.; Freissmuth, M.; Sandtner, W. Kinetic Models of Secondary Active Transporters. International Journal of Molecular Sciences 2019, 20, 5365.

(36) Hussey, G. A.; Thomas, N. E.; Henzler-Wildman, K. A. Highly coupled transport can be achieved in free-exchange transport models. The Journal of General Physiology 2020, 152, e201912437.

(37) Berlaga, A.; Kolomeisky, A. B. Molecular Mechanisms of Active Transport in Antiporters: Kinetic Constraints and Efficiency. The Journal of Physical Chemistry Letters 2021, 12, 9588–9594.

(38) Kenney, I. M.; Beckstein, O. Thermodynamically consistent determination of free energies and rates in kinetic cycle models. Biophysical Reports 2023, 3, 100120.

(39) Wegscheider, R. Über simultane Gleichgewichte und die Beziehungen zwischen Thermodynamik und Reactionskinetik homogener Systeme. Monatshefte für Chemie und verwandte Teile anderer Wissenschaften 1901, 22, 849–906.

(40) Chakraborty, M.; Chowdhury, S.; Chakraborty, J.; Mehera, R.; Pal, R. K. Algorithms for generating all possible spanning trees of a simple undirected connected graph: an extensive review. Complex & Intelligent Systems 2019, 5, 265–281.

(41) Sedgewick, R. Algorithms in C, part 5: graph algorithms, third edition, 3rd ed.; Addison-Wesley Professional, 2001.

(42) Harris, C. R. et al. Array programming with NumPy. Nature 2020, 585, 357–362.

(43) Virtanen, P. et al. SciPy 1.0: fundamental algorithms for scientific computing in Python. Nature Methods 2020, 17, 261–272.

(44) Hunter, J. D. Matplotlib: A 2D Graphics Environment. Computing in Science & Engineering 2007, 9, 90–95, Conference Name: Computing in Science & Engineering.

(45) Hindmarsh, A. C. ODEPACK, A Systematized Collection of ODE Solvers. IMACS Transactions on Scientific Computation 1983, 1, 55–64.

(46) Petzold, L. Automatic Selection of Methods for Solving Stiff and Nonstiff Systems of Ordinary Differential Equations. SIAM Journal on Scientific and Statistical Computing 1983, 4, 136–148.

(47) Gillespie, D.; Petzold, L. Numerical Simulation for Biochemical Kinetics. In System Modeling in Cellular Biology: From Concepts to Nuts and Bolts; The MIT Press, 2006.

(48) Char, J. Generation of Trees, Two-Trees, and Storage of Master Forests. IEEE Transactions on Circuit Theory 1968, 15, 228–238.

(49) Rakshit, A.; Sarma, S. S.; Sen, R. K.; Choudhury, A. An Efficient Tree-Generation Algorithm. IETE Journal of Research 1981, 27, 105–109.

(50) Basuli, K.; Sarma, S.; Naskar, S. Generation of All Spanning Trees of a Simple, Symmetric Connected Graph. SSRN Electronic Journal 2009,

(51) Onete, C. E.; Onete, M. C. C. Enumerating all the spanning trees in an un-oriented graph - A novel approach. 2010 XIth International Workshop on Symbolic and Numerical Methods, Modeling and Applications to Circuit Design (SM2ACD). 2010; pp 1–5.

(52) Călinescu, O.; Paulino, C.; Kühlbrandt, W.; Fendler, K. Keeping It Simple, Transport Mechanism and pH Regulation in Na+/H+ Exchangers. The Journal of Biological Chemistry 2014, 289, 13168–13176.

(53) Matsuoka, R.; Fudim, R.; Jung, S.; Zhang, C.; Bazzone, A.; Chatzikyriakidou, Y.; Robinson, C. V.; Nomura, N.; Iwata, S.; Landreh, M.; Orellana, L.; Beckstein, O.; Drew, D. Structure, mechanism and lipid-mediated remodeling of the mammalian Na+/H+ exchanger NHA2. Nature Structural & Molecular Biology 2022, 29, 108–120.

(54) Crank, J. The Mathematics of Diffusion; Clarendon Press, 1979.

(55) Amdursky, N.; Lin, Y.; Aho, N.; Groenhof, G. Exploring fast proton transfer events associated with lateral proton diffusion on the surface of membranes. Proceedings of the National Academy of Sciences 2019, 116, 2443–2451.

(56) Lewis, O. L.; Keener, J. P.; Fogelson, A. L. A physics-based model for maintenance of the pH gradient in the gastric mucus layer. American Journal of Physiology-Gastrointestinal and Liver Physiology 2017, 313, G599–G612.

(57) Purewal, A. Nucleotide sequence of the ethidium efflux gene from Escherichia coli. FEMS Microbiology Letters 1991, 82, 229–231.

(58) Yerushalmi, H.; Lebendiker, M.; Schuldiner, S. EmrE, an Escherichia coli 12-kDa Multidrug Transporter, Exchanges Toxic Cations and H+ and Is Soluble in Organic Solvents. Journal of Biological Chemistry 1995, 270, 6856–6863.

(59) Robinson, A. E.; Thomas, N. E.; Morrison, E. A.; Balthazor, B. M.; Henzler-Wildman, K. A. New free-exchange model of EmrE transport. Proceedings of the National Academy of Sciences of the United States of America 2017, 114, E10083– E10091.

(60) Morrison, E. A.; Robinson, A. E.; Liu, Y.; Henzler-Wildman, K. A. Asymmetric protonation of EmrE. The Journal of General Physiology 2015, 146, 445–461.

(61) Bell, G. I. Models for the Specific Adhesion of Cells to Cells: A theoretical framework for adhesion mediated by reversible bonds between cell surface molecules. Science 1978, 200, 618–627.

(62) Läuger, P.; Jauch, P. Microscopic description of voltage effects on ion-driven cotransport systems. The Journal of Membrane Biology 1986, 91, 275–284.

