## Supporting Information for "Kinetic Diagram Analysis: A Python Library for Calculating Steady-State Observables of Biochemical Systems Analytically"

**Abstract**

The Supplementary Information contains the detailed example of the simple 4-state model, investigates the effect of nodes in kinetic graphs that are not part of cycles ("dangling nodes"), discusses discrepancies in the symbolic expressions calculated by `KDA` and the `KAPattern` software, provides all quantitative data for the EmrE cycle models, and more details on the equivalence of calculating operational fluxes via cycle fluxes or transition fluxes.

### 1 4-state Example Model

Here we include the master equations and unnormalized state probability expressions for the 4-state kinetic diagram in Fig. 1 in the main paper.

#### 1.1 Master Equation Derivation

For the 4-state model, the kinetic rate matrix $K$ is expressed

$$K = \begin{bmatrix} 0 & k_{12} & 0 & k_{43} \\ k_{21} & 0 & k_{23} & k_{24} \\ 0 & k_{32} & 0 & k_{34} \\ k_{34} & k_{42} & k_{43} & 0 \end{bmatrix},$$

where the kinetic rate matrix is similar to an adjacency matrix in structure, but instead of each element representing the number of edges connected the elements represent the kinetic rate. Applying a simple conversion to the rate matrix,

$$K'_{ij} = \begin{cases} K_{ji}, & \text{if } i \neq j, \\ -\sum_{j} K_{ij}, & \text{if } i = j, \end{cases}$$

we get a matrix which can be used to construct the set of master equations by simply taking the product with the state probabilities (i.e. $\dot{P} = K'P$):

$$\begin{bmatrix} \dot{p_1} \\ \dot{p_2} \\ \dot{p_3} \\ \dot{p_4} \end{bmatrix} = \begin{bmatrix} -(k_{12}+k_{43}) & k_{21} & 0 & k_{34} \\ k_{12} & -(k_{21}+k_{23}+k_{24}) & k_{32} & k_{42} \\ 0 & k_{23} & -(k_{32}+k_{34}) & k_{43} \\ k_{43} & k_{24} & k_{34} & -(k_{34}+k_{42}+k_{43}) \end{bmatrix} \begin{bmatrix} p_1 \\ p_2 \\ p_3 \\ p_4 \end{bmatrix} \quad \text{(S1)}$$

Multiplying out the matrices in Eq. S1 yields the set of master equations for the 4-state model:

$$\begin{aligned} \dot{p_1} &= -p_1(k_{12}+k_{43}) + p_2 k_{21} + p_4 k_{34} \\ \dot{p_2} &= p_1 k_{12} - p_2(k_{21}+k_{23}+k_{24}) + p_3 k_{32} + p_4 k_{42} \\ \dot{p_3} &= p_2 k_{23} - p_3(k_{32}+k_{34}) + p_4 k_{43} \\ \dot{p_4} &= p_1 k_{43} + p_2 k_{24} + p_3 k_{34} - p_4(k_{34}+k_{42}+k_{43}). \end{aligned}$$

#### 1.2 State Probability Expressions

The unnormalized state probability expressions for the 4-state model are defined,

$$\begin{aligned} \Omega_1 &= k_{21}k_{32}k_{41} + k_{21}k_{32}k_{42} + k_{21}k_{32}k_{43} + k_{21}k_{34}k_{41} \\ &\quad + k_{21}k_{34}k_{42} + k_{23}k_{34}k_{41} + k_{24}k_{32}k_{41} + k_{24}k_{34}k_{41} \\ \Omega_2 &= k_{12}k_{32}k_{41} + k_{12}k_{32}k_{42} + k_{12}k_{32}k_{43} + k_{12}k_{34}k_{41} \\ &\quad + k_{12}k_{34}k_{42} + k_{14}k_{32}k_{42} + k_{14}k_{32}k_{43} + k_{14}k_{34}k_{42} \\ \Omega_3 &= k_{12}k_{23}k_{41} + k_{12}k_{23}k_{42} + k_{12}k_{23}k_{43} + k_{12}k_{24}k_{43} \\ &\quad + k_{14}k_{21}k_{43} + k_{14}k_{23}k_{42} + k_{14}k_{23}k_{43} + k_{14}k_{24}k_{43} \\ \Omega_4 &= k_{12}k_{23}k_{34} + k_{12}k_{24}k_{32} + k_{12}k_{24}k_{34} + k_{14}k_{21}k_{32} \\ &\quad + k_{14}k_{21}k_{34} + k_{14}k_{23}k_{34} + k_{14}k_{24}k_{32} + k_{14}k_{24}k_{34}, \end{aligned}$$

where the normalization expression is the sum of all expressions,

$$\begin{aligned} \Sigma &= k_{12}k_{23}k_{34} + k_{12}k_{23}k_{41} + k_{12}k_{23}k_{42} + k_{12}k_{23}k_{43} \\ &\quad + k_{12}k_{24}k_{32} + k_{12}k_{24}k_{34} + k_{12}k_{24}k_{43} + k_{12}k_{32}k_{41} \\ &\quad + k_{12}k_{32}k_{42} + k_{12}k_{32}k_{43} + k_{12}k_{34}k_{41} + k_{12}k_{34}k_{42} \\ &\quad + k_{14}k_{21}k_{32} + k_{14}k_{21}k_{34} + k_{14}k_{21}k_{43} + k_{14}k_{23}k_{34} \\ &\quad + k_{14}k_{23}k_{42} + k_{14}k_{23}k_{43} + k_{14}k_{24}k_{32} + k_{14}k_{24}k_{34} \\ &\quad + k_{14}k_{24}k_{43} + k_{14}k_{32}k_{42} + k_{14}k_{32}k_{43} + k_{14}k_{34}k_{42} \\ &\quad + k_{21}k_{32}k_{41} + k_{21}k_{32}k_{42} + k_{21}k_{32}k_{43} + k_{21}k_{34}k_{41} \\ &\quad + k_{21}k_{34}k_{42} + k_{23}k_{34}k_{41} + k_{24}k_{32}k_{41} + k_{24}k_{34}k_{41}. \end{aligned}$$

Using the 4-state antiporter model (Fig. 1d in the main paper), if we assume the same (symmetrical) binding rates ($L_{\text{on}}$ for ligand and $R_{\text{on}}$ for driving ion) and unbinding rates

($L_{\mathrm{off}}$, $R_{\mathrm{off}}$) on both sides of the membrane, and symmetrical leakage rates,

$$k_{24} = k_{42} = k_{\mathrm{leak}}$$

together with the concentration of ligand on inside and outside ($\mathrm{L}_{\mathrm{int}}$, $\mathrm{L}_{\mathrm{ext}}$) and driving ion ($\mathrm{R}_{\mathrm{int}}$, $\mathrm{R}_{\mathrm{ext}}$), the remaining rates become

$$\begin{aligned}
k_{12} &= k_{14} = R_{\mathrm{off}},\\
k_{21} &= R_{\mathrm{on}}\mathrm{R}_{\mathrm{int}},\\
k_{23} &= L_{\mathrm{on}}\mathrm{L}_{\mathrm{int}},\\
k_{32} &= k_{34} = L_{\mathrm{off}},\\
k_{43} &= L_{\mathrm{on}}\mathrm{L}_{\mathrm{ext}},\\
k_{41} &= R_{\mathrm{on}}\mathrm{R}_{\mathrm{ext}},
\end{aligned}$$

and the normalization expression simplifies to

$$\begin{aligned}
\Sigma = {} & 2\mathrm{L}_{\mathrm{ext}}\mathrm{L}_{\mathrm{int}}L_{\mathrm{on}}^2 R_{\mathrm{off}} + \mathrm{L}_{\mathrm{ext}}L_{\mathrm{off}}L_{\mathrm{on}}\mathrm{R}_{\mathrm{int}}\mathrm{R}_{\mathrm{on}} + 2\mathrm{L}_{\mathrm{ext}}L_{\mathrm{off}}L_{\mathrm{on}}R_{\mathrm{off}}\\
& + \mathrm{L}_{\mathrm{ext}}L_{\mathrm{on}}\mathrm{R}_{\mathrm{int}}R_{\mathrm{off}}\mathrm{R}_{\mathrm{on}} + 2\mathrm{L}_{\mathrm{ext}}L_{\mathrm{on}}R_{\mathrm{off}}k_{\mathrm{leak}} + \mathrm{L}_{\mathrm{int}}L_{\mathrm{off}}L_{\mathrm{on}}\mathrm{R}_{\mathrm{ext}}\mathrm{R}_{\mathrm{on}}\\
& + 2\mathrm{L}_{\mathrm{int}}L_{\mathrm{off}}L_{\mathrm{on}}R_{\mathrm{off}} + \mathrm{L}_{\mathrm{int}}L_{\mathrm{on}}\mathrm{R}_{\mathrm{ext}}R_{\mathrm{off}}\mathrm{R}_{\mathrm{on}} + 2\mathrm{L}_{\mathrm{int}}L_{\mathrm{on}}R_{\mathrm{off}}k_{\mathrm{leak}}\\
& + 2L_{\mathrm{off}}\mathrm{R}_{\mathrm{ext}}\mathrm{R}_{\mathrm{int}}\mathrm{R}_{\mathrm{on}}^2 + 2L_{\mathrm{off}}\mathrm{R}_{\mathrm{ext}}R_{\mathrm{off}}\mathrm{R}_{\mathrm{on}} + 2L_{\mathrm{off}}\mathrm{R}_{\mathrm{ext}}\mathrm{R}_{\mathrm{on}}k_{\mathrm{leak}}\\
& + 2L_{\mathrm{off}}\mathrm{R}_{\mathrm{int}}R_{\mathrm{off}}\mathrm{R}_{\mathrm{on}} + 2L_{\mathrm{off}}\mathrm{R}_{\mathrm{int}}\mathrm{R}_{\mathrm{on}}k_{\mathrm{leak}} + 8L_{\mathrm{off}}R_{\mathrm{off}}k_{\mathrm{leak}}.
\end{aligned}$$

#### 2 Effects of Dangling Nodes on Flux Magnitudes

To explore the effects of adding nodes to diagrams, we created a simple model along with a similar variant with an additional, "dangling" node, for comparison. Dangling nodes do not belong to a cycle but still affect the magnitude of net cycle fluxes and thus operational fluxes. Fig. S1 shows the two kinetic diagrams, a 4-state and 3-state, where the 4-state

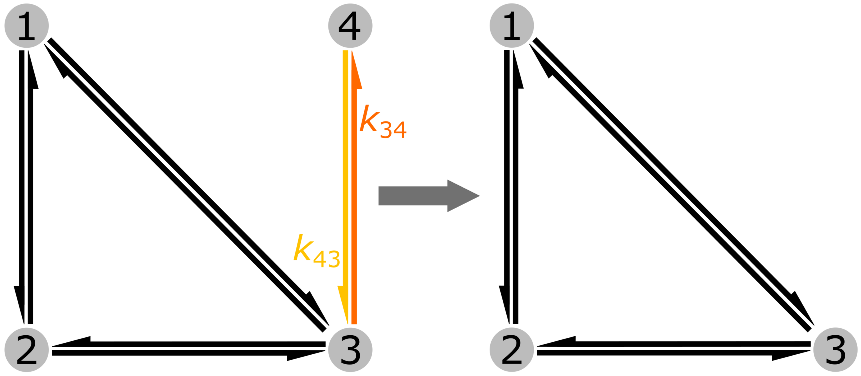


**Figure S1:** 4-state, dangling node model (left) and 3-state model (right).

model is the 3-state model with an additional, dangling node.

To illustrate the effect of adding a dangling node to the 3-state model, we will start by defining the unnormalized state probabilities for both models:

$$
\begin{aligned}
\Omega_1^{\text{3-state}} &= k_{21}k_{31} + k_{21}k_{32} + k_{23}k_{31} \\
\Omega_2^{\text{3-state}} &= k_{12}k_{31} + k_{12}k_{32} + k_{13}k_{32} \\
\Omega_3^{\text{3-state}} &= k_{12}k_{23} + k_{13}k_{21} + k_{13}k_{23} \\
\Omega_1^{\text{4-state}} &= k_{21}k_{31}k_{43} + k_{21}k_{32}k_{43} + k_{23}k_{31}k_{43} \\
\Omega_2^{\text{4-state}} &= k_{12}k_{31}k_{43} + k_{12}k_{32}k_{43} + k_{13}k_{32}k_{43} \\
\Omega_3^{\text{4-state}} &= k_{12}k_{23}k_{43} + k_{13}k_{21}k_{43} + k_{13}k_{23}k_{43} \\
\Omega_4^{\text{4-state}} &= k_{12}k_{23}k_{34} + k_{13}k_{21}k_{34} + k_{13}k_{23}k_{34}.
\end{aligned}
$$

To get the normalized state probabilities the expressions need to be normalized by their respective $\Sigma$ expressions:

$$
\begin{aligned}
\Sigma^{\text{3-state}} &= k_{21}k_{31} + k_{21}k_{32} + k_{23}k_{31} \\
&+ k_{12}k_{31} + k_{12}k_{32} + k_{13}k_{32} \\
&+ k_{12}k_{23} + k_{13}k_{21} + k_{13}k_{23} \\
\Sigma^{\text{4-state}} &= k_{21}k_{31}k_{43} + k_{21}k_{32}k_{43} + k_{23}k_{31}k_{43} \\
&+ k_{12}k_{31}k_{43} + k_{12}k_{32}k_{43} + k_{13}k_{32}k_{43} \\
&+ k_{12}k_{23}k_{43} + k_{13}k_{21}k_{43} + k_{13}k_{23}k_{43} \\
&+ k_{12}k_{23}k_{34} + k_{13}k_{21}k_{34} + k_{13}k_{23}k_{34}.
\end{aligned}
$$

If we compare the state multiplicity expressions for the 3 and 4 state models, we get the following relationships:

$$
\begin{aligned}
\Omega_1^{\text{4-state}} &= k_{43}\,\Omega_1^{\text{3-state}} \\
\Omega_2^{\text{4-state}} &= k_{43}\,\Omega_2^{\text{3-state}} \\
\Omega_3^{\text{4-state}} &= k_{43}\,\Omega_3^{\text{3-state}} \\
\Omega_4^{\text{4-state}} &= k_{34}\,\Omega_3^{\text{3-state}}.
\end{aligned}
$$

Looking at the normalization factors, the 4-state normalization expression can be expressed in terms of the 3-state expressions, $\Sigma^{\text{4-state}} = k_{43}\,\Sigma^{\text{3-state}} + k_{34}\,\Omega_3^{\text{3-state}}$. With the aforementioned relationships considered, the net cycle flux expressions for cycle $k$ for both models are expressed:

$$
\begin{aligned}
J_K^{\text{3-state}} &= \left(\Pi_k^{+} - \Pi_k^{-}\right)/\Sigma^{\text{3-state}} \\
J_K^{\text{4-state}} &= \frac{k_{43}\left(\Pi_k^{+} - \Pi_k^{-}\right)}{k_{43}\,\Sigma^{\text{3-state}} + k_{34}\,\Omega_3^{\text{3-state}}},
\end{aligned}
$$

where $K$ is the cycle common to both models. Under the condition $k_{34}, k_{43} > 0$, the ratio of the net cycle fluxes is expressed:

$$\frac{J_k^{\text{4-state}}}{J_k^{\text{3-state}}} = \frac{k_{43}\,\Sigma^{\text{3-state}}}{k_{43}\,\Sigma^{\text{3-state}} + k_{34}\,\Omega_3^{\text{3-state}}} = \frac{\Omega_1^{\text{3-state}} + \Omega_2^{\text{3-state}} + \Omega_3^{\text{3-state}}}{\Omega_1^{\text{3-state}} + \Omega_2^{\text{3-state}} + (1 + \frac{k_{34}}{k_{43}})\Omega_3^{\text{3-state}}} \leq 1 \tag{S2}$$
$$J_k^{\text{4-state}} \leq J_k^{\text{3-state}}.$$

Generally speaking $J_k^{\text{4-state}} < J_k^{\text{3-state}}$. The equality is only met under the limit as $k_{34}$ approaches zero,

$$\lim_{k_{34}\to 0} p_4^{\text{4-state}} = 0, \text{ and } \lim_{k_{34}\to 0} J_k^{\text{4-state}} = J_k^{\text{3-state}}.$$

Conversely, for the limit as $k_{43}$ approaches zero,

$$\lim_{k_{43}\to 0} p_4^{\text{4-state}} = 1, \text{ and } \lim_{k_{43}\to 0} J_k^{\text{4-state}} = 0.$$

Lastly, we can use Eq. S2 to look at the flux ratio for the case where $k_{34} = k_{43}$:

$$\frac{J_k^{\text{4-state}}}{J_k^{\text{3-state}}} = \frac{\Omega_1^{\text{3-state}} + \Omega_2^{\text{3-state}} + \Omega_3^{\text{3-state}}}{\Omega_1^{\text{3-state}} + \Omega_2^{\text{3-state}} + (1 + \frac{k_{34}}{k_{43}})\Omega_3^{\text{3-state}}} = \frac{\Sigma^{\text{3-state}}}{\Sigma^{\text{3-state}} + \Omega_3^{\text{3-state}}}.$$

Therefore, with the exception of the case where $k_{34}$ approaches zero, adding a dangling node will reduce the net cycle flux regardless of the magnitudes of the transition rates between the dangling node and the original diagram.

### 3 Min-Max Normalization

The transition fluxes and state probabilities can be visualized by plotting the kinetic diagram and varying the edge widths and node sizes according to the values of interest. When plotting steady-state observables (e.g. probabilities, transition fluxes) over the kinetic diagram, to scale the diagram nodes and edges such that deviations are visibly distinguishable we use min-max normalization. The simple case is expressed

$$x_{\text{scaled}} = \frac{(x - x_{\text{min}})}{x_{\text{max}} - x_{\text{min}}},$$

where $x$ is the value to be scaled (out of a set of $x$ values), $x_{\text{min}}$ and $x_{\text{max}}$ are the minimum and maximum values for the data set, respectively. This scales any data set between the values [0, 1], but can be generalized to scale between an arbitrary minimum and maximum. This generalized version is expressed

$$x_{\text{scaled}} = \frac{(x - x_{\text{min}})(b - a)}{x_{\text{max}} - x_{\text{min}}} + a, \tag{S3}$$

where $a$ and $b$ are the lower and upper scaled values for the output scaled values, respectively. This is useful for our case since the nodes and edges must be scaled according to different boundaries in the plotting code.

Flux values may span many orders of magnitude so we used a modified version of Eq.

S3 where base-10 log scaling is used to re-scale the values before min-max normalization is applied:

$$x_{\text{scaled}} = \frac{(\log_{10} x - \log_{10} x_{\min})(b-a)}{\log_{10} x_{\max} - \log_{10} x_{\min}} + a. \tag{S4}$$

For our purposes, scaling nodes via state probability values Eq. S3 is used with $a = 60$, $b = 500$, $x_{\min} = 0$, and $x_{\max} = 1$. For scaling edges via transition fluxes, Eq. S4 is used with $a = 1.5$ and $b = 3.0$. $x_{\min}$ and $x_{\max}$ values are set based on the minimum and maximum transition flux values in the diagram. For scaling edges via net transition fluxes, Eq. S3 is used with $a = 0.2$ and $b = 1.5$. $x_{\min}$ and $x_{\max}$ values are set based on the minimum and maximum net transition flux values in the diagram. Using this method, transition and net transition flux edges cannot be compared directly within the same graph as they are scaled with different $a$ and $b$ values. Transition and net transition fluxes cannot be compared across graphs with different parameters since their $x_{\min}$ and $x_{\min}$ are case-specific.

### 4 KDA vs. KAPattern Expression Disagreement

For validation purposes we compared the outputs of `KDA` to the expressions created by the `KAPattern` software.[1] To compare expressions, we selected a subset of moderately complex diagrams with 3 to 7 states and used `SymPy`[2] to make the comparison. For some of our test models there were disagreements between the `KDA` and `KAPattern` expressions. Here we will include one such case where disagreements were found and a discussion why the `KAPattern` expression is incorrect. The code used to compare expressions along with the terminal output is located in the `KDA` examples repository (github.com/Becksteinlab/kda-examples) in the `kda_paper/fig_10__kda_validation_and_performance/kapattern_comparison` directory.

For the 7-state test model (Fig. S2) expression disagreements were observed for states 1, 6 and 7. To discover the source of the disagreements the expressions were compared by eye. For example, for state 1 the difference between both expressions has four remaining terms,

$$\begin{aligned} p_1^{\text{KDA}} - p_1^{\text{KAPattern}} = & \\ & + k_{25}k_{34}k_{41}k_{54}k_{65}k_{73} \text{ (KDA)} \\ & + k_{25}k_{34}k_{41}k_{54}k_{67}k_{73} \text{ (KDA)} \\ & - k_{25}k_{34}k_{41}k_{45}k_{54}k_{65}k_{73} \text{ (KAPattern)} \\ & - k_{25}k_{34}k_{41}k_{45}k_{54}k_{67}k_{73} \text{ (KAPattern)}. \end{aligned}$$

Every term in a state probability expression comes from a directional diagram and thus must only contain 6 rates for a 7-state diagram. Additionally, each directional diagram represents a directed path along the kinetic diagram and thus can never include both forward and reverse rates from a given transition. For these reasons, the `KAPattern`-generated terms represent invalid directional diagrams since they 1.) include a total of 7 rates, and 2.) include both forward and reverse rates for the $4 \leftrightarrow 5$ transition (i.e. $k_{45}$ and $k_{54}$). For all expression disagreements found (states 1 and 6 for the 6-state diagram and states 1, 6 and 7 for the 7-state diagram) the `KAPattern` rate-products break one or both directional diagram rules for each case.

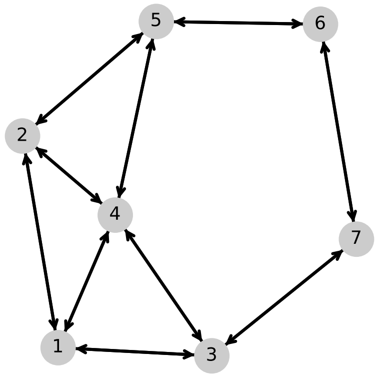


**Figure S2:** 7-state randomly generated kinetic diagram from the `KDA` validation data (index 7_17).

### 5 EmrE Flux Relationships

Due to the internal consistency of Hill's diagram method, the various fluxes can be expressed in terms of the others. The operational fluxes can be expressed in terms of either net cycle fluxes or net transition fluxes and the net transition fluxes can be expressed in terms of net cycle fluxes.[3] Here we will first derive the net transition flux relationships for the 8-state model of EmrE[4] (Fig. 12a) directly from the master equations, then show how these identities are equal to the operational fluxes.

Starting with the kinetic differential equations expressed in terms of net transition fluxes at steady-state (i.e. $\dot{p}_i = 0$), we develop identities between individual net transition fluxes and sums of two others:

$$\begin{aligned}
\dot{p}_1 &= J_{7,1} + J_{2,1} + J_{3,1} = 0 \implies J_{7,1} = J_{1,2} + J_{1,3} \\
\dot{p}_2 &= J_{1,2} + J_{8,2} + J_{4,2} = 0 \implies J_{8,2} = J_{2,1} + J_{2,4} \\
\dot{p}_3 &= J_{1,3} + J_{4,3} + J_{5,3} = 0 \\
\dot{p}_4 &= J_{2,4} + J_{3,4} + J_{6,4} = 0 \implies J_{6,4} = J_{4,2} + J_{4,3} \\
\dot{p}_5 &= J_{3,5} + J_{6,5} + J_{7,5} = 0 \\
\dot{p}_6 &= J_{4,6} + J_{5,6} + J_{8,6} = 0 \implies J_{6,4} = J_{5,6} + J_{8,6} \\
\dot{p}_7 &= J_{5,7} + J_{8,7} + J_{1,7} = 0 \implies J_{7,1} = J_{5,7} + J_{8,7} \\
\dot{p}_8 &= J_{2,8} + J_{6,8} + J_{7,8} = 0 \implies J_{8,2} = J_{6,8} + J_{7,8},
\end{aligned}$$

where we make use of the property $J_{ij} = -J_{ji}$ for any net transition flux. The chosen relationships are expressed in terms of the alternating access transitions from inside to outside, anticipating relationships between their sums. Summing the appropriate expressions (i.e. ones with canceling transitions), we find the following relationships:

$$J_{7,1} + J_{8,2} = J_{1,3} + J_{2,4} = J_{6,8} + J_{5,7} \tag{S5}$$

$$J_{6,4} + J_{8,2} = J_{7,8} + J_{5,6} = J_{2,1} + J_{4,3}. \tag{S6}$$

Upon inspection, the expressions in Eq. S5 are the sums of net transition fluxes for transitions that transport bound protons from inside to outside, unbind external protons, and bind internal protons, respectively. Similarly, the expressions in Eq. S6 are sums of net transition fluxes for transitions which transport bound drug molecules from inside to outside, bind internal drug molecules, and unbind external drug molecules, respectively. It is not surprising then that the expressions in Eqs. S5 and S6 will be shown to be equal to the operational fluxes for protons and drug molecules, respectively.

To prove these relationships we will show the net transition fluxes expressed in terms of the net cycle fluxes. Many combinations of transitions (and their corresponding net transition fluxes) can be combined to get the operational fluxes, but the specific transitions shown here are chosen due to their specific biochemical processes (i.e. alternating access, ligand binding/unbinding). In Fig. 12a in the main paper, the transitions $7 \to 1$, $8 \to 2$, and $6 \to 4$ are the alternating access transitions moving ligands from inside to outside. In terms of net cycle fluxes, their transition fluxes are expressed as

$$\begin{aligned} J_{7,1} &= J_{\mathbf{1}} + J_{\mathbf{2}} + J_{\mathbf{3}} + J_{\mathbf{4}} + J_{\mathbf{5}} + J_{\mathbf{6}} + J_{\mathbf{7}} \\ &\quad + J_{\mathbf{8}} + J_{\mathbf{9}} + J_{\mathbf{10}} + J_{\mathbf{11}} + J_{\mathbf{12}} + J_{\mathbf{13}} + J_{\mathbf{14}} \\ J_{8,2} &= -J_{\mathbf{5}} - J_{\mathbf{6}} - J_{\mathbf{7}} - J_{\mathbf{8}} - J_{\mathbf{9}} - J_{\mathbf{12}} \\ &\quad + J_{\mathbf{16}} + J_{\mathbf{17}} + J_{\mathbf{19}} + J_{\mathbf{20}} + J_{\mathbf{22}} + J_{\mathbf{23}} + J_{\mathbf{24}} + J_{\mathbf{25}} \\ J_{6,4} &= -J_{\mathbf{3}} - J_{\mathbf{4}} + J_{\mathbf{5}} - J_{\mathbf{10}} - J_{\mathbf{11}} + J_{\mathbf{12}} \\ &\quad + J_{\mathbf{15}} + J_{\mathbf{18}} - J_{\mathbf{19}} - J_{\mathbf{20}} - J_{\mathbf{22}} - J_{\mathbf{25}} + J_{\mathbf{26}} + J_{\mathbf{27}}. \end{aligned}$$

Summing the appropriate transition pairs according to Eqs. S5-S6 yields identical result as Eqs. 17-18 in the main paper, confirming the first operational flux relationship for each ligand.

For the the net transition fluxes related to the binding/unbinding of the ligands, we focus on the transition pairs $1 \to 3$, $2 \to 4$, and $6 \to 8$, $5 \to 7$ for the protons, and transition pairs $7 \to 8$, $5 \to 6$, $2 \to 1$, $4 \to 3$ for the drug molecules. The net transition fluxes for the proton binding/unbinding transitions expressed in terms of net cycle fluxes are

$$\begin{aligned} J_{1,3} &= +J_{\mathbf{2}} + J_{\mathbf{3}} + J_{\mathbf{5}} + J_{\mathbf{6}} + J_{\mathbf{8}} + J_{\mathbf{10}} + J_{\mathbf{14}} \\ &\quad + J_{\mathbf{15}} + J_{\mathbf{16}} + J_{\mathbf{17}} + J_{\mathbf{18}} + J_{\mathbf{19}} + J_{\mathbf{20}} + J_{\mathbf{21}} \\ J_{2,4} &= +J_{\mathbf{1}} + J_{\mathbf{4}} - J_{\mathbf{5}} - J_{\mathbf{6}} - J_{\mathbf{8}} + J_{\mathbf{11}} + J_{\mathbf{13}} \\ &\quad - J_{\mathbf{15}} - J_{\mathbf{18}} - J_{\mathbf{21}} + J_{\mathbf{22}} + J_{\mathbf{23}} + J_{\mathbf{24}} + J_{\mathbf{25}} \\ J_{6,8} &= +J_{\mathbf{1}} + J_{\mathbf{2}} + J_{\mathbf{3}} + J_{\mathbf{4}} - J_{\mathbf{8}} - J_{\mathbf{9}} - J_{\mathbf{12}} \\ &\quad - J_{\mathbf{15}} + J_{\mathbf{17}} + J_{\mathbf{19}} + J_{\mathbf{24}} + J_{\mathbf{25}} - J_{\mathbf{26}} - J_{\mathbf{28}} \\ J_{5,7} &= +J_{\mathbf{8}} + J_{\mathbf{9}} + J_{\mathbf{10}} + J_{\mathbf{11}} + J_{\mathbf{12}} + J_{\mathbf{13}} + J_{\mathbf{14}} \\ &\quad + J_{\mathbf{15}} + J_{\mathbf{16}} + J_{\mathbf{20}} + J_{\mathbf{22}} + J_{\mathbf{23}} + J_{\mathbf{26}} + J_{\mathbf{28}}, \end{aligned}$$

and the net transition fluxes for the drug binding/unbinding transitions expressed in terms of net cycle fluxes are

$$
\begin{aligned}
J_{7,8} &= -\mathbf{J_1} - \mathbf{J_2} - \mathbf{J_3} - \mathbf{J_4} - \mathbf{J_5} - \mathbf{J_6} - \mathbf{J_7} \\
&\quad + \mathbf{J_{15}} + \mathbf{J_{16}} + \mathbf{J_{20}} + \mathbf{J_{22}} + \mathbf{J_{23}} + \mathbf{J_{26}} + \mathbf{J_{28}} \\
J_{5,6} &= +\mathbf{J_1} + \mathbf{J_2} + \mathbf{J_5} - \mathbf{J_8} - \mathbf{J_9} - \mathbf{J_{10}} - \mathbf{J_{11}} \\
&\quad + \mathbf{J_{17}} + \mathbf{J_{18}} - \mathbf{J_{20}} - \mathbf{J_{22}} + \mathbf{J_{24}} + \mathbf{J_{27}} - \mathbf{J_{28}} \\
J_{2,1} &= -\mathbf{J_1} - \mathbf{J_4} - \mathbf{J_7} - \mathbf{J_9} - \mathbf{J_{11}} - \mathbf{J_{12}} - \mathbf{J_{13}} \\
&\quad + \mathbf{J_{15}} + \mathbf{J_{16}} + \mathbf{J_{17}} + \mathbf{J_{18}} + \mathbf{J_{19}} + \mathbf{J_{20}} + \mathbf{J_{21}} \\
J_{4,3} &= +\mathbf{J_1} - \mathbf{J_3} - \mathbf{J_6} - \mathbf{J_8} - \mathbf{J_{10}} + \mathbf{J_{12}} + \mathbf{J_{13}} \\
&\quad - \mathbf{J_{19}} - \mathbf{J_{20}} - \mathbf{J_{21}} + \mathbf{J_{23}} + \mathbf{J_{24}} + \mathbf{J_{26}} + \mathbf{J_{27}}.
\end{aligned}
$$

Summing the net transition fluxes for the appropriate transition pairs yields the result in Eqs. 17-18, confirming the remaining net transition flux expressions in Eqs. S5-S6 are equal to the operational fluxes:

$$J_{\mathrm{H^+}} = J_{7,1} + J_{8,2} = J_{1,3} + J_{2,4} = J_{6,8} + J_{5,7} \tag{S7}$$

$$J_{\mathrm{D}} = J_{6,4} + J_{8,2} = J_{7,8} + J_{5,6} = J_{2,1} + J_{4,3}. \tag{S8}$$

### 6 EmrE Net Cycle Fluxes

In our analysis of EmrE, operational fluxes were calculated by summing distinct sets of net cycle fluxes. Here we include the numerical results for all net cycle fluxes evaluated under both alternating access rate biasing conditions (Table S1) and substrate off-rate biasing conditions (Table S2). For the substrate off-rate biasing case, we include only the $k_{\mathrm{AA}} = 100\ \mathrm{s}^{-1}$ results.

### 7 Operational Flux Comparison Numerical Data

Here we include the operational flux numerical data calculated using both net cycle fluxes and net transition fluxes for two different models. Table S3 contains the numerical data for the 6-state antiporter model and Table S4 contains the numerical data for the 8-state model of EmrE.

**Table S1:** EmrE Net Cycle Fluxes for Alternating Access Rate Biasing. Fluxes have units of cycle completions per second and are labeled according to Fig. 13 in the main paper. Values of 0 are numerically identical to zero.

| Cycle | $R_{AA} = 10^{-8}$ | $R_{AA} = 1$ | $R_{AA} = 10^{8}$ |
|---|---|---|---|
| 1 | 0 | $1.1 \times 10^{-9}$ | 0 |
| 2 | $6.0 \times 10^{-9}$ | $6.4 \times 10^{-5}$ | $1.9 \times 10^{-8}$ |
| 3 | $2.0 \times 10^{-14}$ | $1.4 \times 10^{-2}$ | $6.0 \times 10^{-2}$ |
| 4 | $6.6 \times 10^{-16}$ | $4.5 \times 10^{-3}$ | $1.9 \times 10^{-2}$ |
| 5 | 0 | 0 | 0 |
| 6 | 0 | 0 | 0 |
| 7 | 0 | 0 | 0 |
| 8 | 0 | 0 | 0 |
| 9 | 0 | 0 | 0 |
| 10 | $3.8 \times 10^{-16}$ | $4.5 \times 10^{-3}$ | $1.9 \times 10^{-2}$ |
| 11 | $1.9 \times 10^{-14}$ | $1.4 \times 10^{-3}$ | $6.0 \times 10^{-3}$ |
| 12 | 0 | 0 | 0 |
| 13 | $6.0 \times 10^{-8}$ | $2.0 \times 10^{-4}$ | $1.9 \times 10^{-8}$ |
| 14 | $4.2 \times 10^{-4}$ | $2.0 \times 10^{0}$ | $4.2 \times 10^{-4}$ |
| 15 | 0 | 0 | 0 |
| 16 | $1.9 \times 10^{-2}$ | $4.5 \times 10^{-3}$ | $6.6 \times 10^{-16}$ |
| 17 | $6.0 \times 10^{-3}$ | $1.4 \times 10^{-3}$ | $1.9 \times 10^{-14}$ |
| 18 | 0 | 0 | 0 |
| 19 | $1.9 \times 10^{-8}$ | $1.9 \times 10^{-4}$ | $6.0 \times 10^{-8}$ |
| 20 | 0 | $1.1 \times 10^{-9}$ | 0 |
| 21 | 0 | 0 | 0 |
| 22 | $1.9 \times 10^{-8}$ | $5.9 \times 10^{-5}$ | $6.0 \times 10^{-9}$ |
| 23 | $6.0 \times 10^{-2}$ | $1.4 \times 10^{-2}$ | $2.0 \times 10^{-14}$ |
| 24 | $1.9 \times 10^{-2}$ | $4.5 \times 10^{-3}$ | $3.8 \times 10^{-16}$ |
| 25 | $1.0 \times 10^{-4}$ | $4.9 \times 10^{-1}$ | $1.0 \times 10^{-4}$ |
| 26 | 0 | 0 | 0 |
| 27 | 0 | 0 | 0 |
| 28 | 0 | 0 | 0 |

**Table S2:** EmrE Net Cycle Fluxes for Substrate Off-Rate Biasing. Fluxes have units of cycle completions per second and are labeled according to Fig. 13 in the main paper. All fluxes are evaluated with $k_{\mathrm{AA}} = 100\ \mathrm{s}^{-1}$. Values of 0 are numerically identical to zero.

| Cycle | $R_{\mathrm{off}} = 10^{-10}$ | $R_{\mathrm{off}} = 1$ | $R_{\mathrm{off}} = 10^{2}$ | $R_{\mathrm{off}} = 10^{10}$ |
|---|---|---|---|---|
| 1 | $1.8 \times 10^{-12}$ | $1.1 \times 10^{-8}$ | $1.3 \times 10^{-10}$ | 0 |
| 2 | $7.5 \times 10^{-8}$ | $7.1 \times 10^{-4}$ | $3.8 \times 10^{-5}$ | $1.7 \times 10^{-12}$ |
| 3 | $2.4 \times 10^{-6}$ | $1.6 \times 10^{-2}$ | $3.4 \times 10^{-4}$ | $5.5 \times 10^{-15}$ |
| 4 | $9.7 \times 10^{-2}$ | $6.1 \times 10^{-3}$ | $7.1 \times 10^{-6}$ | 0 |
| 5 | 0 | 0 | 0 | 0 |
| 6 | 0 | 0 | 0 | 0 |
| 7 | 0 | 0 | 0 | 0 |
| 8 | 0 | 0 | 0 | 0 |
| 9 | 0 | 0 | 0 | 0 |
| 10 | $7.3 \times 10^{-12}$ | $5.5 \times 10^{-3}$ | $1.6 \times 10^{-3}$ | $3.4 \times 10^{-10}$ |
| 11 | $3.0 \times 10^{-7}$ | $2.1 \times 10^{-3}$ | $4.4 \times 10^{-5}$ | $7.0 \times 10^{-16}$ |
| 12 | 0 | 0 | 0 | 0 |
| 13 | $9.5 \times 10^{-8}$ | $2.1 \times 10^{-3}$ | $1.9 \times 10^{-4}$ | $1.7 \times 10^{-12}$ |
| 14 | $1.0 \times 10^{-3}$ | $2.3 \times 10^{1}$ | $1.0 \times 10^{1}$ | $1.4 \times 10^{-3}$ |
| 15 | 0 | 0 | 0 | 0 |
| 16 | $9.7 \times 10^{-12}$ | $6.2 \times 10^{-3}$ | $9.1 \times 10^{-4}$ | $1.7 \times 10^{-8}$ |
| 17 | $3.0 \times 10^{-13}$ | $2.1 \times 10^{-3}$ | $2.7 \times 10^{-3}$ | $1.7 \times 10^{-6}$ |
| 18 | 0 | 0 | 0 | 0 |
| 19 | $2.3 \times 10^{-12}$ | $1.9 \times 10^{-3}$ | $7.0 \times 10^{-4}$ | $1.7 \times 10^{-10}$ |
| 20 | 0 | $1.1 \times 10^{-8}$ | $1.3 \times 10^{-8}$ | $4.2 \times 10^{-15}$ |
| 21 | 0 | 0 | 0 | 0 |
| 22 | $3.0 \times 10^{-13}$ | $6.4 \times 10^{-4}$ | $5.5 \times 10^{-4}$ | $1.7 \times 10^{-10}$ |
| 23 | $2.3 \times 10^{-12}$ | $1.6 \times 10^{-2}$ | $1.8 \times 10^{-2}$ | $1.7 \times 10^{-6}$ |
| 24 | $3.7 \times 10^{-16}$ | $5.4 \times 10^{-3}$ | $5.4 \times 10^{-2}$ | $1.7 \times 10^{-4}$ |
| 25 | $1.9 \times 10^{-5}$ | $5.4 \times 10^{-1}$ | $1.5 \times 10^{-1}$ | $3.4 \times 10^{-8}$ |
| 26 | 0 | 0 | 0 | 0 |
| 27 | 0 | 0 | 0 | 0 |
| 28 | 0 | 0 | 0 | 0 |

**Table S3:** 6-State Antiporter Operational Flux Calculation Method Comparison. Flux values are computed as IEEE-754 doubles but shown with reduced precision. All values are expressed with units of $\mathrm{s}^{-1}$. Values of 0 are numerically identical to zero.

| $k_{\mathrm{leak}}$ | $J_{\mathrm{H}^+}$ | $J_{\mathrm{H}^+} - J_{1,2}$ | $J_{\mathrm{Na}^+}$ | $J_{\mathrm{Na}^+} - J_{6,1}$ |
|---|---|---|---|---|
| 0.01 | $2.05 \times 10^{1}$ | 0 | $2.05 \times 10^{1}$ | 0 |
| 1 | $2.06 \times 10^{1}$ | 0 | $2.02 \times 10^{1}$ | 0 |
| 100 | $2.78 \times 10^{1}$ | 0 | $-1.67$ | 0 |

**Table S4:** EmrE Operational Flux Calculation Method Comparison. Flux values are computed as IEEE-754 doubles but shown with reduced precision and units of $s^{-1}$. Values of 0 are numerically identical to zero.

| | $J_{H^+}$ | $J_{H^+} - (J_{1,3} + J_{2,4})$ | $J_D$ | $J_D - (J_{2,1} + J_{4,3})$ |
|---|---|---|---|---|
| $R_{AA} = 10^{-8}$ | $1.04 \times 10^{-1}$ | $2.19 \times 10^{-13}$ | $1.04 \times 10^{-1}$ | $2.18 \times 10^{-13}$ |
| $R_{AA} = 1$ | 2.57 | 0 | 0 | $5.55 \times 10^{-16}$ |
| $R_{AA} = 10^{8}$ | $1.04 \times 10^{-1}$ | $3.34 \times 10^{-12}$ | $-1.04 \times 10^{-1}$ | $-3.43 \times 10^{-13}$ |
| $R_{off} = 10^{-10\dagger}$ | $9.79 \times 10^{-2}$ | $4.77 \times 10^{-14}$ | $-9.69 \times 10^{-2}$ | 0 |
| $R_{off} = 1^{\dagger}$ | $2.36 \times 10^{1}$ | $-9.95 \times 10^{-14}$ | $1.24 \times 10^{-18}$ | $7.01 \times 10^{-16}$ |
| $R_{off} = 100^{\dagger}$ | $1.03 \times 10^{1}$ | 0 | $7.30 \times 10^{-2}$ | $3.05 \times 10^{-16}$ |

† $R_{off}$ values are evaluated with $k_{AA} = 100\ s^{-1}$.

### 8 Operational Flux Expression Comparison Code

Here we include Python code used to compare the operational flux expressions generated using net cycle fluxes or net transition fluxes for two different models. The scripts are also available in the `KDA` examples repository (github.com/Becksteinlab/kda-examples) in the `kda_paper/operational_flux_comparisons` directory.

#### 8.1 6-state Antiporter Model Operational Flux Expression Comparison Code

For the leakage variant of the 6-state antiporter model (Fig. 11b in the main paper, model $G_{leak}$) the operational fluxes can be expressed in terms of net cycle fluxes:

$$J_{H^+} = J_A + J_C$$
$$J_{Na^+} = J_B + J_C,$$

or single net transition fluxes

$$J_{H^+} = J_{1,2} = J_{2,3} = J_{3,4}$$
$$J_{Na^+} = J_{6,1} = J_{5,6} = J_{4,5}.$$

Using `KDA` and `SymPy`[2] we compare the algebraic expressions directly by executing the code below.

```
import numpy as np
import networkx as nx
from sympy import symbols
from kda import graph_utils, calculations as calcs

# define connectivity matrix
K = np.array(
    [
```

```
        [0, 1, 0, 1, 0, 1],
        [1, 0, 1, 0, 0, 0],
        [0, 1, 0, 1, 0, 0],
        [1, 0, 1, 0, 1, 0],
        [0, 0, 0, 1, 0, 1],
        [1, 0, 0, 0, 1, 0],
    ]
)
# create the 6-state kinetic diagram
G_leak = nx.MultiDiGraph()
# populate edge data using KDA utility
graph_utils.generate_edges(G_leak, K)
# create symbols for current variables
(k12, k21, k23, k32, k34, k43, k45,
    k54, k56, k65, k61, k16, k14, k41) = symbols(
    "k12 k21 k23 k32 k34 k43 k45 k54 k56 k65 k61 k16 k14, k41")
# create variables for substitution
(H_on, H_off, Na_on, Na_off, k_conf,
k_leak,  H_in, H_out, c_Na) = symbols(
    "H_on, H_off, Na_on, Na_off, k_conf, k_leak,  H_in, H_out, c_Na")
# define variable substitution mapping
model = {
    k12:H_on*H_out, k21:H_off,
    k23:k_conf, k32:k_conf,
    k34:H_off, k43:H_on*H_in,
    k45:Na_on*c_Na, k54:Na_off,
    k56:k_conf, k65:k_conf,
    k61:Na_off, k16:Na_on*c_Na,
    k14:k_leak, k41:k_leak,
}
# collect the cycles using KDA
cycles = graph_utils.find_all_unique_cycles(G_leak)
# label cycles according to figure
cycle_labels = ["a", "c", "b"]
# set the cycle directions (CW)
cycle_orders = [[0, 1], [0, 1], [5, 0]]
# generate net cycle flux expressions
G_leak_cycles = {}
for label, _cycle, _order in zip(cycle_labels, cycles, cycle_orders):
    func = calcs.calc_net_cycle_flux(
        G_leak, cycle=_cycle, order=_order, key='name', output_strings=
   True)
    # perform variable substitutions
    func = func.subs(model).simplify()
    G_leak_cycles[label] = {"cycle": _cycle, "order": _order, "func": func
   }
# assign net cycle flux expressions
J_a = G_leak_cycles["a"]["func"]
J_b = G_leak_cycles["b"]["func"]
J_c = G_leak_cycles["c"]["func"]
# generate state probability expressions
prob_strs = calcs.calc_state_probs(
    G_leak, key='name', output_strings=True)
p1, p2, p3, p4, p5, p6 = prob_strs
```

```
# calculate operational fluxes using cycles
J_H_cycle = (J_a + J_c).simplify()
J_Na_cycle = (J_b + J_c).simplify()
# calculate operational fluxes using transition fluxes
# J_H = p1 * k12 - p2 * k21 = J12
J_H_trans = (p1.subs(model) * model[k12] - p2.subs(model) * model[k21]).
    simplify()
# J_Na = p6 * k61 - p1 * k16 = J61
J_Na_trans = (p6.subs(model) * model[k61] - p1.subs(model) * model[k16]).
    simplify()

# compare operational flux expressions
# created using cycles and transitions
print((J_H_cycle - J_H_trans).simplify() == 0)
print((J_Na_cycle - J_Na_trans).simplify() == 0)
```

#### 8.2 8-state EmrE Model Operational Flux Expression Comparison Code

For the 8-state model of EmrE (Fig. 12a in the main paper) the operational fluxes can be expressed in terms of net cycle fluxes:

$$\begin{aligned}
J_{\mathrm{H}^+} &= + J_{\mathbf{1}} + J_{\mathbf{2}} + J_{\mathbf{3}} + J_{\mathbf{4}} + J_{\mathbf{10}} + J_{\mathbf{11}} + J_{\mathbf{13}} + J_{\mathbf{14}} \\
&\quad + J_{\mathbf{16}} + J_{\mathbf{17}} + J_{\mathbf{19}} + J_{\mathbf{20}} + J_{\mathbf{22}} + J_{\mathbf{23}} + J_{\mathbf{24}} + J_{\mathbf{25}} \\
J_{\mathrm{D}} &= - J_{\mathbf{3}} - J_{\mathbf{4}} - J_{\mathbf{6}} - J_{\mathbf{7}} - J_{\mathbf{8}} - J_{\mathbf{9}} - J_{\mathbf{10}} - J_{\mathbf{11}} \\
&\quad + J_{\mathbf{15}} + J_{\mathbf{16}} + J_{\mathbf{17}} + J_{\mathbf{18}} + J_{\mathbf{23}} + J_{\mathbf{24}} + J_{\mathbf{26}} + J_{\mathbf{27}},
\end{aligned}$$

or sums of net transition fluxes (Eqs. S7-S8). Using `KDA` and `SymPy`[2] we compare the algebraic expressions directly by executing the code below.

```
import sys
# raise the recursion limit so SymPy doesn't raise a RecursionError
# when parsing the EmrE normalization expression (sigma)
sys.setrecursionlimit(5000)
import numpy as np
import networkx as nx
from sympy import symbols
from sympy.parsing.sympy_parser import parse_expr
from kda import graph_utils, diagrams, calculations as calcs

# define connectivity matrix
K = np.array(
    [
        [0, 1, 1, 0, 0, 0, 1, 0],
        [1, 0, 0, 1, 0, 0, 0, 1],
        [1, 0, 0, 1, 1, 0, 0, 0],
        [0, 1, 1, 0, 0, 1, 0, 0],
        [0, 0, 1, 0, 0, 1, 1, 0],
        [0, 0, 0, 1, 1, 0, 0, 1],
        [1, 0, 0, 0, 1, 0, 0, 1],
        [0, 1, 0, 0, 0, 1, 1, 0],
```

```
    ]
)
# create the 8-state kinetic diagram
G = nx.MultiDiGraph()
# populate edge data using KDA utility
graph_utils.generate_edges(G, K)
# create symbols for current variables
(k31, k13, k57, k75, k42, k24, k68, k86, k34, k43, k56, k65,
  k12, k21, k78, k87, k71, k17, k53, k35, k64, k46, k82, k28) = symbols(
  "k31 k13 k57 k75 k42 k24 k68 k86 k34 k43 k56 k65
     k12 k21 k78 k87 k71 k17 k53 k35 k64 k46 k82 k28")
# create variables for substitution
(H_on, H_off, D_on, D_off, H_in, H_out,
    D_in, D_out, k_AA_anti, k_AA_sym) = symbols(
    "H_on H_off D_on D_off H_in H_out D_in D_out k_AA_anti k_AA_sym")
# define variable substitution mapping
model = {
    k31: H_on * H_out, k13: H_off,
    k57: H_on * H_in, k75: H_off,
    k42: H_on * H_out, k24: H_off,
    k68: H_on * H_in, k86: H_off,
    k34: D_on * D_out, k43: D_off,
    k56: D_on * D_in, k65: D_off,
    k12: D_on * D_out, k21: D_off,
    k78: D_on * D_in, k87: D_off,
    k71: k_AA_anti, k17: k_AA_anti,
    k53: k_AA_sym, k35: k_AA_sym,
    k64: k_AA_anti, k46: k_AA_anti,
    k82: k_AA_sym, k28: k_AA_sym,
}

# collect the directional edges beforehand
dir_edges = diagrams.generate_directional_diagrams(G, return_edges=True)
# create the normalization factor for net cycle fluxes
sigma = calcs.calc_sigma(G, dir_edges, key="name", output_strings=True)
sigma = parse_expr(sigma)
sigma = sigma.subs(model).simplify()

# define cycle info
EmrE_cycles = {
  1: {"cycle": [0, 1, 3, 2, 4, 5, 7, 6], "order": [6, 0]},
  2: {"cycle": [0, 6, 7, 5, 4, 2], "order": [6, 0]},
  3: {"cycle": [0, 6, 7, 5, 3, 2], "order": [6, 0]},
  4: {"cycle": [0, 6, 7, 5, 3, 1], "order": [6, 0]},
  6: {"cycle": [0, 6, 7, 1, 3, 2], "order": [6, 0]},
  7: {"cycle": [0, 6, 7, 1], "order": [6, 0]},
  8: {"cycle": [0, 2, 3, 1, 7, 5, 4, 6], "order": [6, 0]},
  9: {"cycle": [0, 6, 4, 5, 7, 1], "order": [6, 0]},
  10: {"cycle": [0, 6, 4, 5, 3, 2], "order": [6, 0]},
  11: {"cycle": [0, 6, 4, 5, 3, 1], "order": [6, 0]},
  13: {"cycle": [0, 6, 4, 2, 3, 1], "order": [6, 0]},
  14: {"cycle": [0, 6, 4, 2], "order": [6, 0]},
  15: {"cycle": [0, 1, 3, 5, 7, 6, 4, 2], "order": [0, 2]},
  16: {"cycle": [0, 2, 4, 6, 7, 1], "order": [0, 2]},
```

```
  17: {"cycle": [0, 2, 4, 5, 7, 1], "order": [0, 2]},
  18: {"cycle": [0, 2, 4, 5, 3, 1], "order": [0, 2]},
  19: {"cycle": [0, 2, 3, 5, 7, 1], "order": [0, 2]},
  20: {"cycle": [0, 1, 7, 6, 4, 5, 3, 2], "order": [0, 2]},
  22: {"cycle": [1, 7, 6, 4, 5, 3], "order": [1, 3]},
  23: {"cycle": [1, 7, 6, 4, 2, 3], "order": [2, 4]},
  24: {"cycle": [1, 7, 5, 4, 2, 3], "order": [2, 4]},
  25: {"cycle": [1, 7, 5, 3], "order": [1, 3]},
  26: {"cycle": [2, 3, 5, 7, 6, 4], "order": [2, 4]},
  27: {"cycle": [2, 3, 5, 4], "order": [2, 4]},
  }
# generate net cycle flux expression numerators
for idx, cycle_info in EmrE_cycles.items():
  pi_diff = calcs.calc_pi_difference(G, cycle=cycle_info["cycle"],
    order=cycle_info["order"], key="name", output_strings=True)
  flux_diags = diagrams.generate_flux_diagrams(G, cycle=cycle_info["cycle"
   ])
  sigma_K = calcs.calc_sigma_K(G, cycle=cycle_info["cycle"],
    flux_diags=flux_diags, key="name", output_strings=True)
  if sigma_K == 1:
    func = parse_expr(pi_diff)
  else:
    func = parse_expr(pi_diff) * parse_expr(sigma_K)
  # perform variable substitutions
  func = func.subs(model).simplify()
  EmrE_cycles[idx]["func"] = func
# define contributing cycles for both ligands
H_cycles = [1, 2, 3, 4, 10, 11, 13, 14, 16, 17, 19, 20, 22, 23, 24, 25]
D_cycles = [3, 4, 6, 7, 8, 9, 10, 11, 15, 16, 17, 18, 23, 24, 26, 27]
J_H_cycle = 0
for idx in H_cycles:
  J_H_cycle += EmrE_cycles[idx]["func"]
J_D_cycle = 0
for idx in D_cycles:
  # negative contributors
  if idx in (3, 4, 6, 7, 8, 9, 10, 11):
    J_D_cycle -= EmrE_cycles[idx]["func"]
  else:
    J_D_cycle += EmrE_cycles[idx]["func"]
# normalize the operational fluxes (cycles)
J_H_cycle = (J_H_cycle / sigma).simplify()
J_D_cycle = (J_D_cycle / sigma).simplify()
# generate state probability expressions
prob_strs = calcs.calc_state_probs(G, key='name', output_strings=True)
p1, p2, p3, p4, p5, p6, p7, p8 = prob_strs
# calculate operational fluxes using transition fluxes
J_13 = p1.subs(model) * model[k13] - p3.subs(model) * model[k31]
J_24 = p2.subs(model) * model[k24] - p4.subs(model) * model[k42]
J_21 = p2.subs(model) * model[k21] - p1.subs(model) * model[k12]
J_43 = p4.subs(model) * model[k43] - p3.subs(model) * model[k34]
J_H_trans = (J_13 + J_24).simplify()
J_D_trans = (J_21 + J_43).simplify()
# compare operational flux expressions
# created using cycles and transitions
```

```
print((J_H_cycle - J_H_trans).simplify() == 0)
print((J_D_cycle - J_D_trans).simplify() == 0)
```

#### References


(1) Qi, F.; Dash, R. K.; Han, Y.; Beard, D. A. Generating rate equations for complex enzyme systems by a computer-assisted systematic method. *BMC Bioinformatics* **2009**, *10*, 238.

(2) Meurer, A. et al. SymPy: symbolic computing in Python. *PeerJ Computer Science* **2017**, *3*, e103.

(3) Hill, T. L. *Free Energy Transduction and Biochemical Cycle Kinetics*; Springer-Verlag: New York, 1989.

(4) Hussey, G. A.; Thomas, N. E.; Henzler-Wildman, K. A. Highly coupled transport can be achieved in free-exchange transport models. *The Journal of General Physiology* **2020**, *152*, e201912437.
